## Supplement for "Post-translational modifications of *Drosophila melanogaster* HOX protein, Sex combs reduced"

**Figure S1. Coverage map of embryonic SCRTT.** Each blue line underneath the primary protein sequence of SCRTT represents a chemically distinct peptide identified by MS/MS analysis. The modifications are indicated by letters or symbols on the blue lines. The C-terminal triple tag sequence is represented by the faded region. On the right is the legend for all modifications shown in the figure. Amino acid substitutions have been excluded from the figure. A. Analysis of tryptic peptides. A1, A2, A3 and A4 are independent MS/MS analyses. B. Analysis of chymotryptic peptides. A1 and A2 are independent MS/MS analyses. C. Analysis of thermolytic peptides. A1 and A2 are independent MS/MS analyses.

**A**

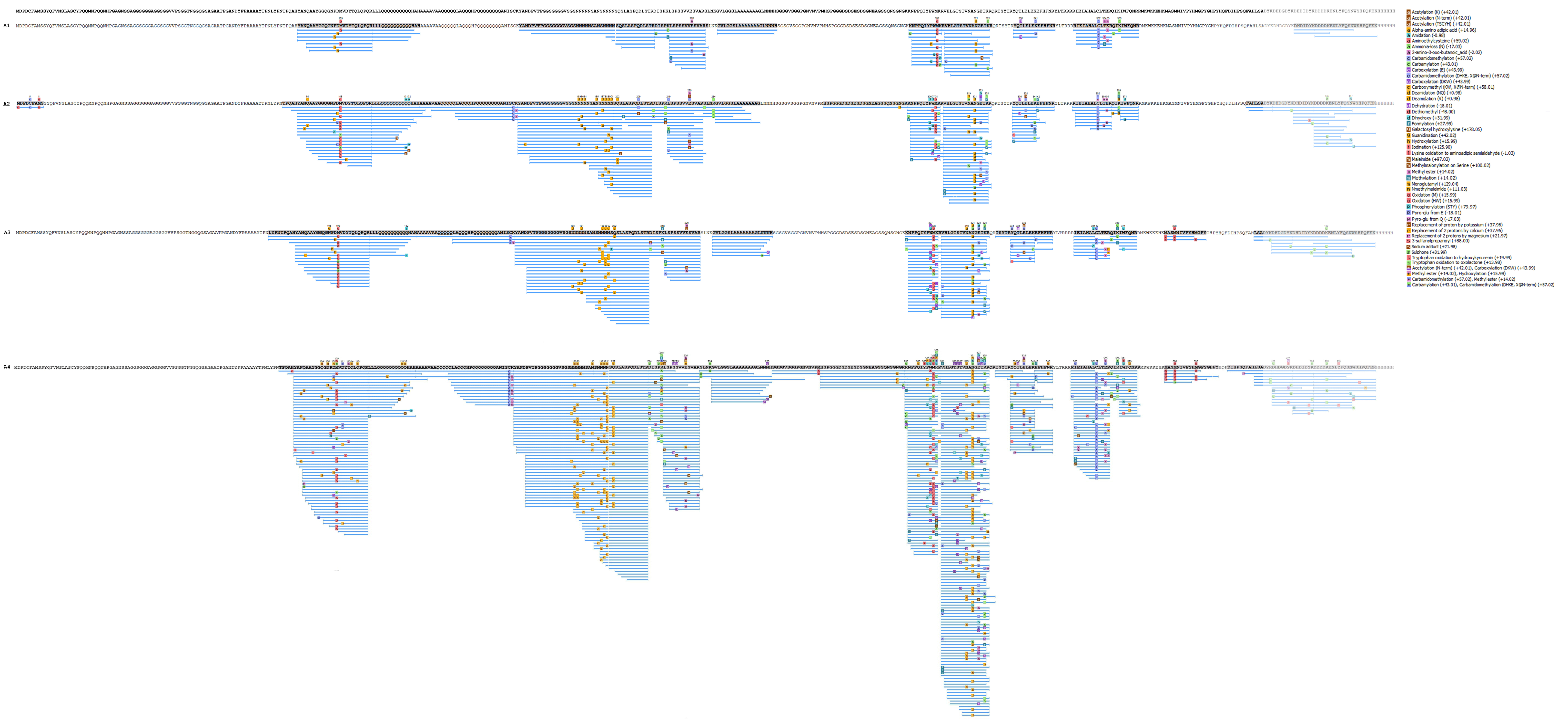
A1
A2
A3
A4
Acetylation (K) (+42.01)
Acetylation (N-term) (+42.01)
Acetylation (TSCYH) (+42.01)
Alpha-amino adipic acid (+14.96)
Amidation (-0.98)
Aminoethylcysteine (+59.02)
Ammonia-loss (N) (-17.03)
2-amino-3-oxo-butanoic_acid (-2.02)
Carbamidomethylation (+57.02)
Carbamylation (+43.01)
Carboxylation (E) (+43.99)
Carbamidomethylation (DHKE, X@N-term) (+57.02)
Carboxylation (DKW) (+43.99)
Carboxymethyl (KW, X@N-term) (+58.01)
Deamidation (NQ) (+0.98)
Deamidation (R) (+0.98)
Dehydration (-18.01)
Dethiomethyl (-48.00)
Dihydroxy (+31.99)
Formylation (+27.99)
Galactosyl hydroxylysine (+178.05)
Guanidination (+42.02)
Hydroxylation (+15.99)
Iodination (+125.90)
Lysine oxidation to aminoadipic semialdehyde (-1.03)
Maleimide (+97.02)
Methylmalonylation on Serine (+100.02)
Methyl ester (+14.02)
Methylation (+14.02)
Monoglutamyl (+129.04)
Nmethylmaleimide (+111.03)
Oxidation (M) (+15.99)
Oxidation (HW) (+15.99)
Phosphorylation (STY) (+79.97)
Pyro-glu from E (-18.01)
Pyro-glu from Q (-17.03)
Replacement of proton by potassium (+37.96)
Replacement of 2 protons by calcium (+37.95)
Replacement of 2 protons by magnesium (+21.97)
3-sulfanylpropanoyl (+88.00)
Sodium adduct (+21.98)
Sulphone (+31.99)
Tryptophan oxidation to hydroxykynurenin (+19.99)
Tryptophan oxidation to oxolactone (+13.98)
Acetylation (N-term) (+42.01), Carboxylation (DKW) (+43.99)
Methyl ester (+14.02), Hydroxylation (+15.99)
Carbamidomethylation (+57.02), Methyl ester (+14.02)
Carbamylation (+43.01), Carbamidomethylation (DHKE, X@N-term) (+57.02)

**B**

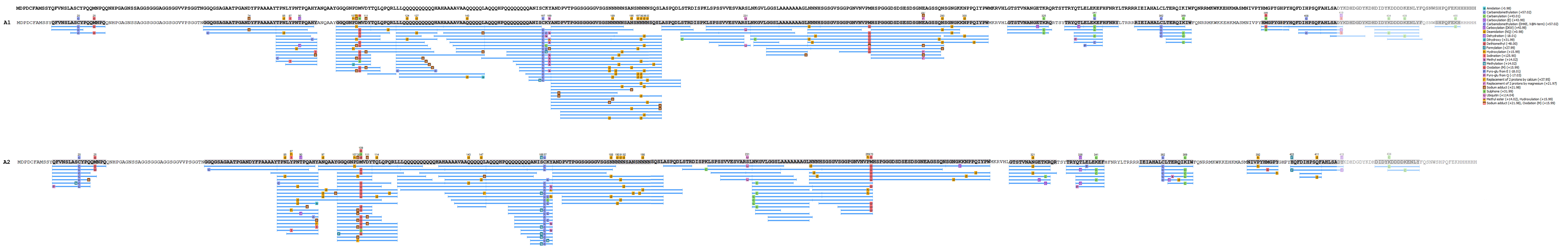

C

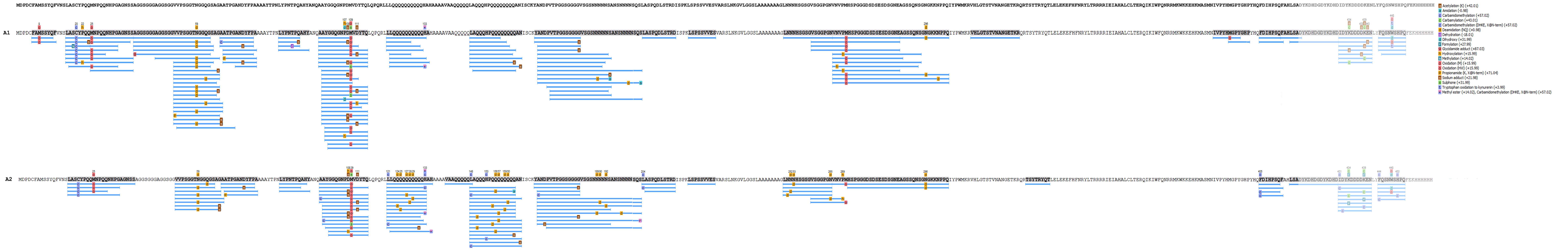

A1
A2
a Acetylation (K) (+42.01)
a Amidation (-0.98)
c Carbamidomethylation (+57.02)
c Carbamylation (+43.01)
c Carbamidomethylation (DHKE, X@N-term) (+57.02)
d Deamidation (NQ) (+0.98)
d Dehydration (-18.01)
d Dihydroxy (+31.99)
f Formylation (+27.99)
g Glycidamide adduct (+87.03)
h Hydroxylation (+15.99)
m Methylation (+14.02)
o Oxidation (M) (+15.99)
o Oxidation (HW) (+15.99)
p Propionamide (K, X@N-term) (+71.04)
s Sodium adduct (+21.98)
s Sulphone (+31.99)
t Tryptophan oxidation to kynurenin (+3.99)
* Methyl ester (+14.02), Carbamidomethylation (DHKE, X@N-term) (+57.02)

**Figure S2. Coverage map of commercially purchased bovine α-casein.** Each blue line underneath the primary protein sequence of α-casein represents a chemically distinct tryptic peptide identified by MS/MS analysis. The modifications are indicated by letters or symbols on the blue lines. On the right is the legend for all modifications shown in the figure. Amino acid substitutions have been excluded from the figure. A. Analysis of α-casein without $TiO_2$ treatment. B. Analysis of α-casein with $TiO_2$ treatment.

A

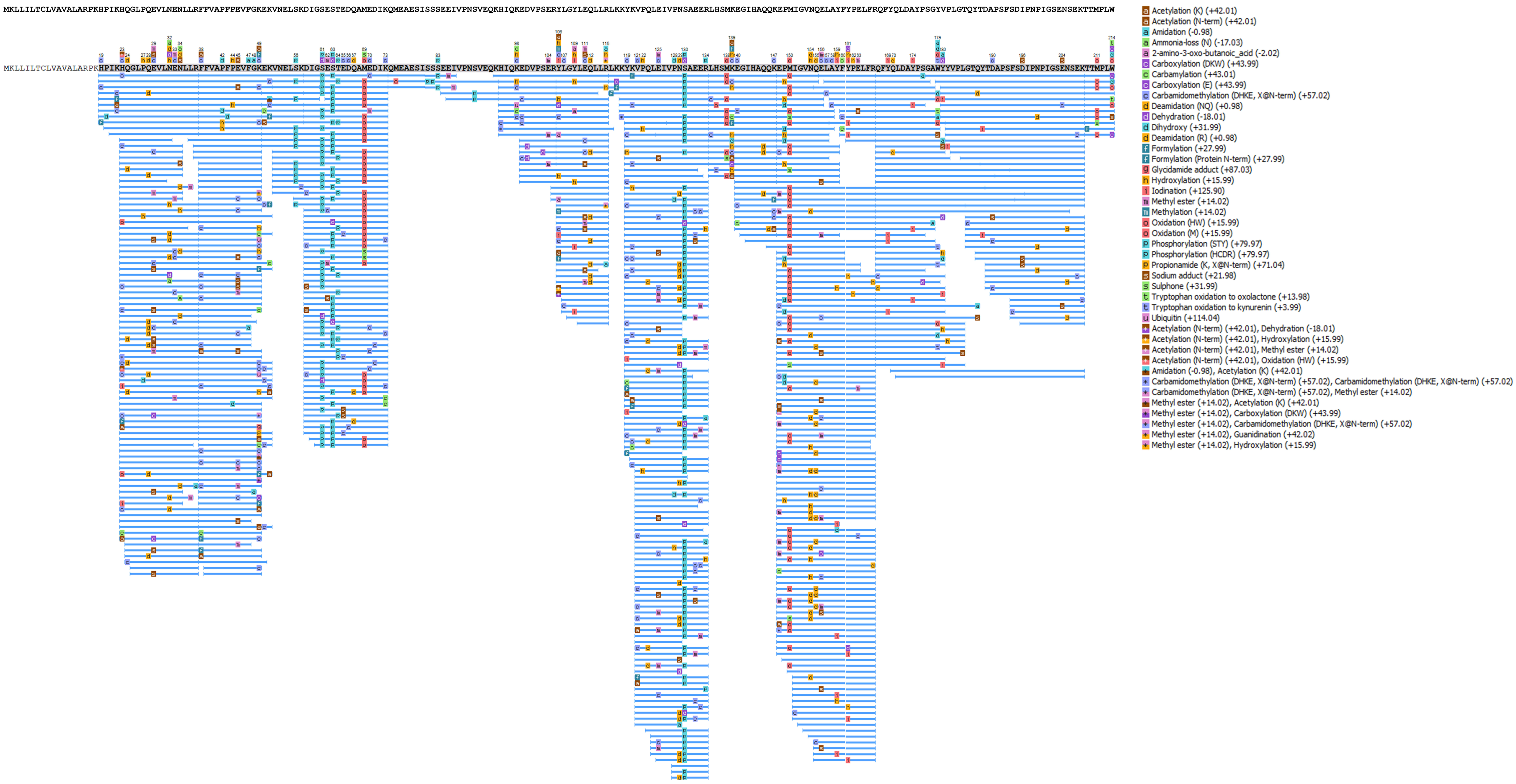
MKLLILTCLVAVALARPKHPIKHQGLPQEVLNENLLRFFVAPFPEVFGKEKVNELSKDIGSESTEDQAMEDIKQMEAESISSSEEIVPNSVEQKHIQKEDVPSERYLGYLEQLLRLKKYKVPQLEIVPNSAEERLHSMKEGIHAQQKEPMIGVNQELAYFYPELFRQFYQLDAYPSGYVPLGTQYTDAPSFSDIPNPIGSENSEKTTMPLW
Acetylation (K) (+42.01)
Acetylation (N-term) (+42.01)
Amidation (-0.98)
Ammonia-loss (N) (-17.03)
2-amino-3-oxo-butanoic_acid (-2.02)
Carboxylation (DKW) (+43.99)
Carbamylation (+43.01)
Carboxylation (E) (+43.99)
Carbamidomethylation (DHKE, X@N-term) (+57.02)
Deamidation (NQ) (+0.98)
Dehydration (-18.01)
Dihydroxy (+31.99)
Deamidation (R) (+0.98)
Formylation (+27.99)
Formylation (Protein N-term) (+27.99)
Glycidamide adduct (+87.03)
Hydroxylation (+15.99)
Iodination (+125.90)
Methyl ester (+14.02)
Methylation (+14.02)
Oxidation (HW) (+15.99)
Oxidation (M) (+15.99)
Phosphorylation (STY) (+79.97)
Phosphorylation (HCDR) (+79.97)
Propionamide (K, X@N-term) (+71.04)
Sodium adduct (+21.98)
Sulphone (+31.99)
Tryptophan oxidation to oxolactone (+13.98)
Tryptophan oxidation to kynurenin (+3.99)
Ubiquitin (+114.04)
Acetylation (N-term) (+42.01), Dehydration (-18.01)
Acetylation (N-term) (+42.01), Hydroxylation (+15.99)
Acetylation (N-term) (+42.01), Methyl ester (+14.02)
Acetylation (N-term) (+42.01), Oxidation (HW) (+15.99)
Amidation (-0.98), Acetylation (K) (+42.01)
Carbamidomethylation (DHKE, X@N-term) (+57.02), Carbamidomethylation (DHKE, X@N-term) (+57.02)
Carbamidomethylation (DHKE, X@N-term) (+57.02), Methyl ester (+14.02)
Methyl ester (+14.02), Acetylation (K) (+42.01)
Methyl ester (+14.02), Carboxylation (DKW) (+43.99)
Methyl ester (+14.02), Carbamidomethylation (DHKE, X@N-term) (+57.02)
Methyl ester (+14.02), Guanidination (+42.02)
Methyl ester (+14.02), Hydroxylation (+15.99)

**B**

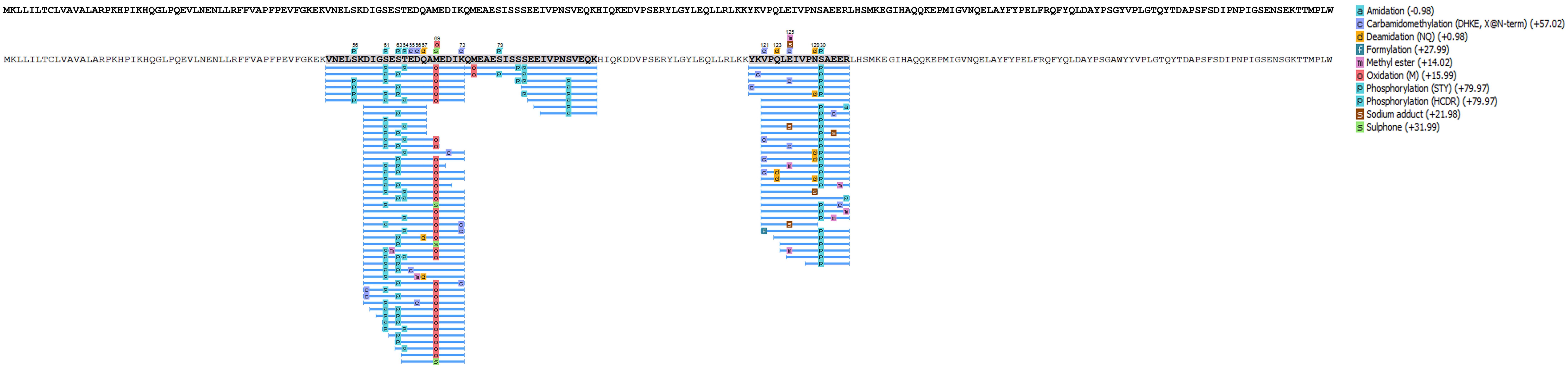

MKLLILTCLVAVALARPKHPIKHQGLPQEVLNENLLRFFVAPFPEVFGKEKVNELSKDIGSESTEDQAMEDIKQMEAESISSSEEIVPNSVEQKHIQKEDVPSERYLGYLEQLLRLKKYKVPQLEIVPNSAEERLHSMKEGIHAQQKEPMIGVNQELAYFYPELFRQFYQLDAYPSGYVPLGTQYTDAPSFSDIPNPIGSENSEKTTMPLW
a Amidation (-0.98)
c Carbamidomethylation (DHKE, X@N-term) (+57.02)
d Deamidation (NQ) (+0.98)
f Formylation (+27.99)
m Methyl ester (+14.02)
o Oxidation (M) (+15.99)
p Phosphorylation (STY) (+79.97)
p Phosphorylation (HCDR) (+79.97)
s Sodium adduct (+21.98)
s Sulphone (+31.99)

**Figure S3. Phosphorylation of Serine 185, Serine 201, Threonine 315, Serine 316, Threonine 317 and Threonine 324 residues of SCRTT.** $MS^2$ spectra for the three phosphopeptides identified by LC-MS/MS are shown. A. Phosphorylation of Serine 185. B. Phosphorylation of Serine 201. The inset box shows fragment ions with *m/z* 1690 to 1790. C. Phosphorylation of Threonine 315. D. Phosphorylation of Serine 316. E. Phosphorylation of Threonine 317. The inset box shows fragment ions with *m/z* 900 to 1400. F. Phosphorylation of Threonine 324. The peptide sequence and *m/z* ratio are indicated on the top of the spectra. Positions of fragmentation are shown with vertical lines in the peptide sequence. The box on the right summarizes the evidences of phosphorylation. The relevant fragment ions and their *m/z* ratios supporting phosphorylation are labelled in the spectra.

**A** $MS^2$ *m/z* **1118.73**

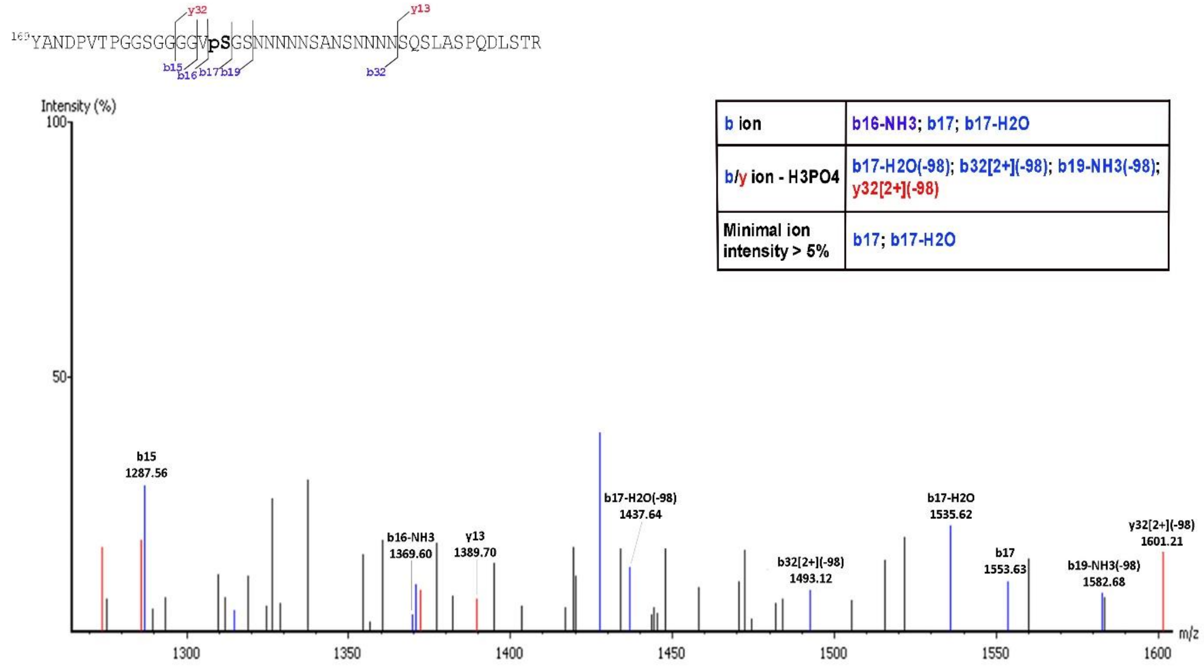

**B** MS² *m/z* 1118.98

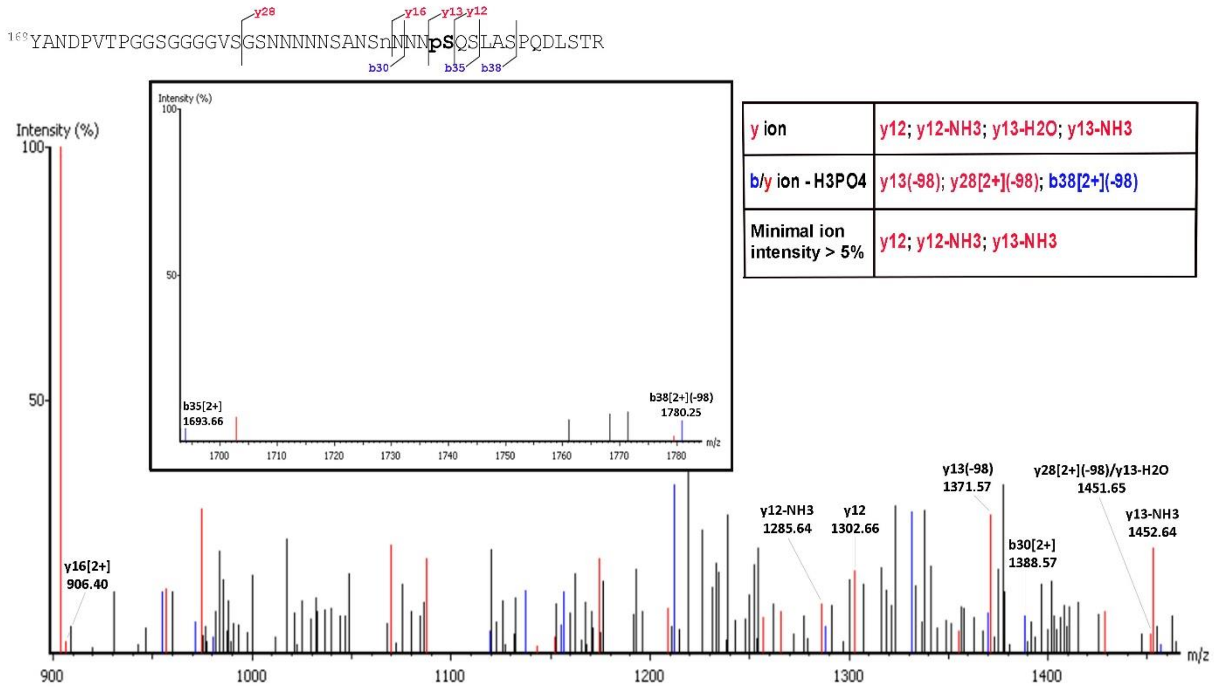


| y ion | y12; y12-NH3; y13-H2O; y13-NH3 |
|---|---|
| b/y ion - H3PO4 | y13(-98); y28[2+](-98); b38[2+](-98) |
| Minimal ion intensity > 5% | y12; y12-NH3; y13-NH3 |

**C** MS² *m/z* 441.96

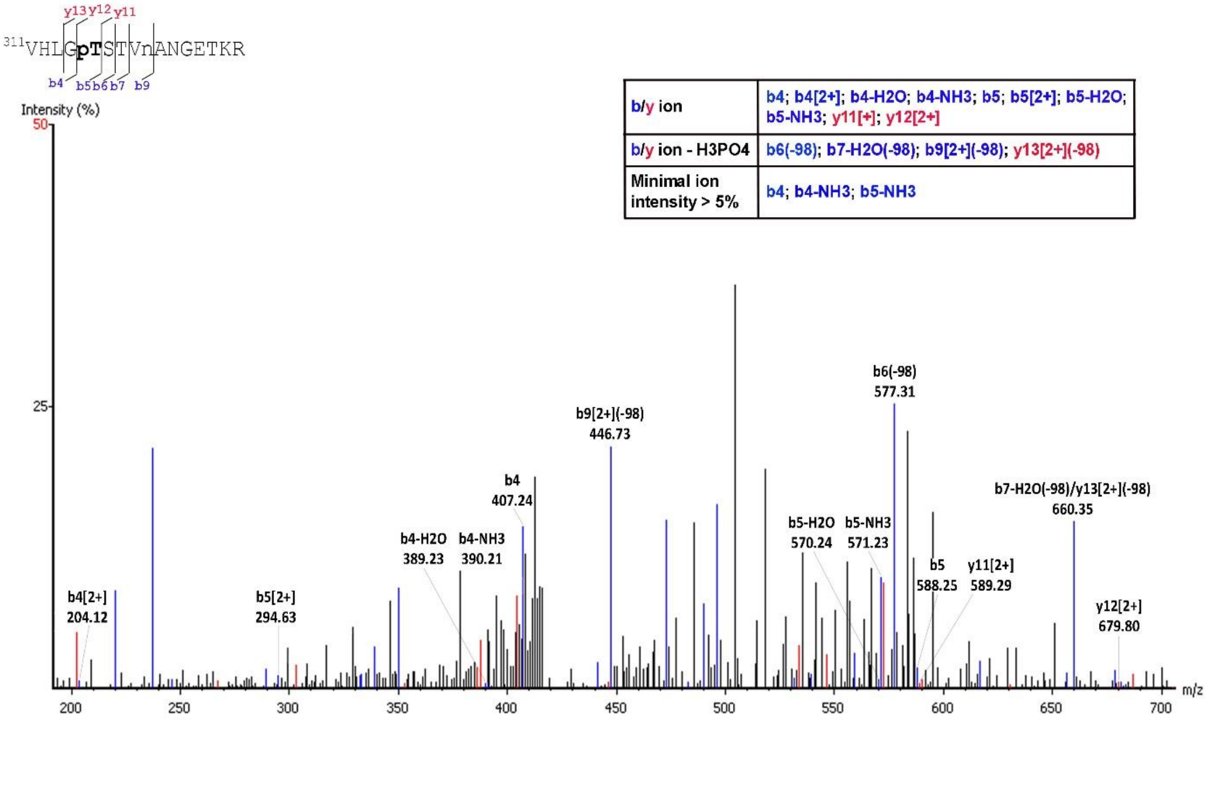


| b/y ion | b4; b4[2+]; b4-H2O; b4-NH3; b5; b5[2+]; b5-H2O; b5-NH3; y11[+]; y12[2+] |
|---|---|
| b/y ion - H3PO4 | b6(-98); b7-H2O(-98); b9[2+](-98); y13[2+](-98) |
| Minimal ion intensity > 5% | b4; b4-NH3; b5-NH3 |

**D** MS$^2$ *m/z* 442.21

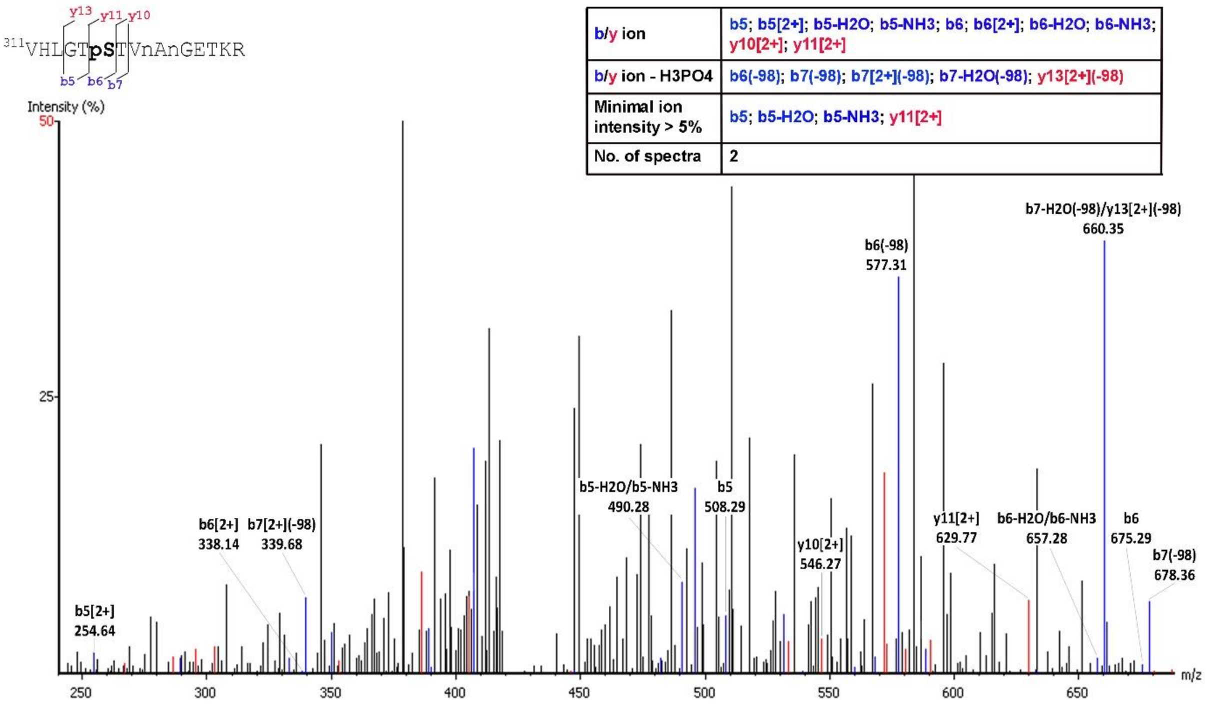


**E** MS$^2$ *m/z* 441.96

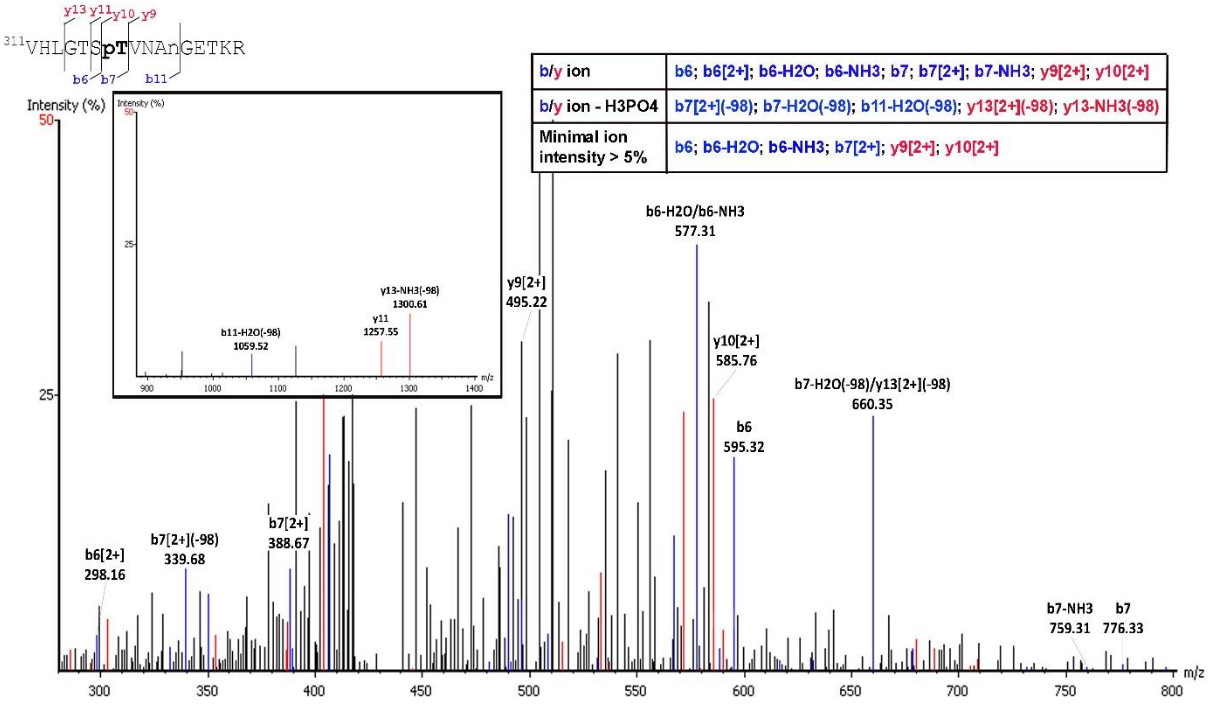

**F MS$^2$ *m/z* 442.21**

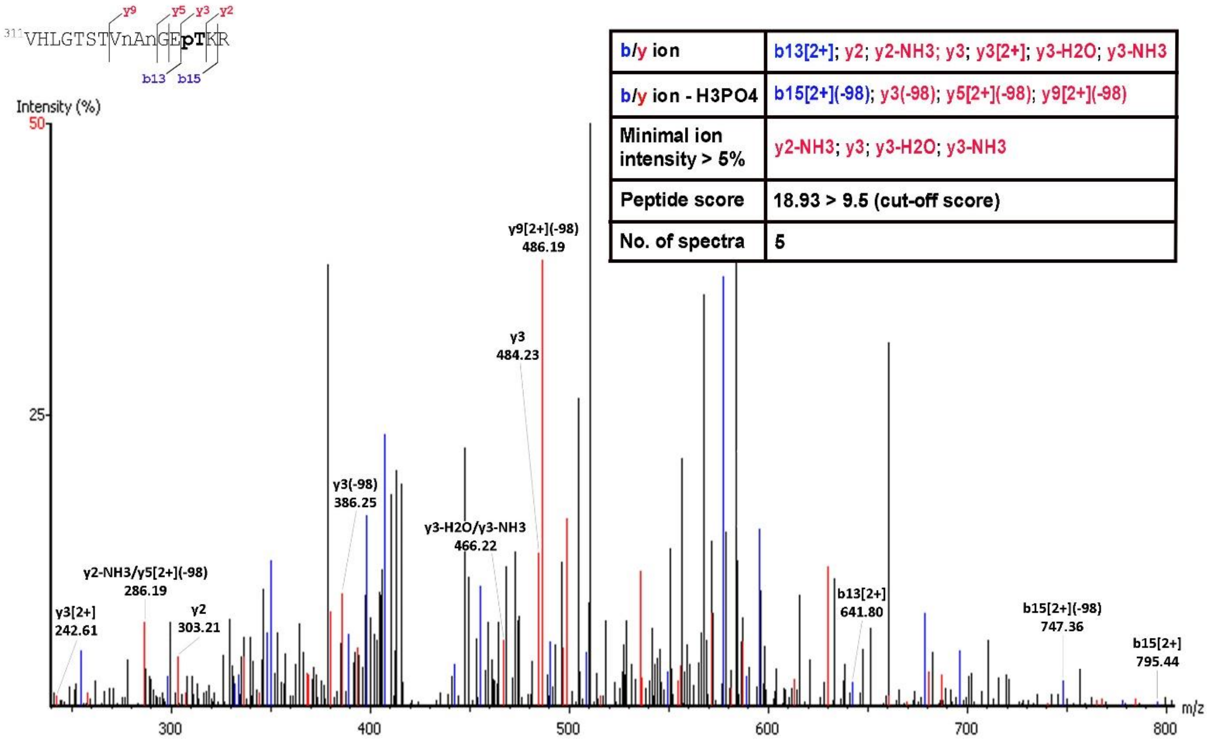


| | |
|---|---|
| b/y ion | b13[2+]; y2; y2-NH3; y3; y3[2+]; y3-H2O; y3-NH3 |
| b/y ion - H3PO4 | b15[2+](-98); y3(-98); y5[2+](-98); y9[2+](-98) |
| Minimal ion intensity > 5% | y2-NH3; y3; y3-H2O; y3-NH3 |
| Peptide score | 18.93 > 9.5 (cut-off score) |
| No. of spectra | 5 |

**Figure S4. Acetylation of Lysine 218, Serine 223, Serine 227, Lysine 309, Lysine 434 and Lysine 439 residues of SCRTT.** $MS^2$ spectra of the peptide identified by LC-MS/MS is shown. A. Acetylation of Lysine 218. B. Acetylation of Serine 223. C. Acetylation of Serine 227. D. Acetylation of Lysine 309. The inset box shows fragment ions with *m/z* 1150 to 1350. E. Acetylation of Lysine 434. F. Acetylation of Lysine 439. The inset box shows fragment ions with *m/z* 390 to 580. The peptide sequence and *m/z* ratio are indicated at the top of the spectra. Positions of fragmentation are shown with vertical lines in the peptide sequence. The box on the right summarizes the evidence confirming acetylation. The relevant fragment ions and their *m/z* ratios supporting acetylation are labelled in the spectra.

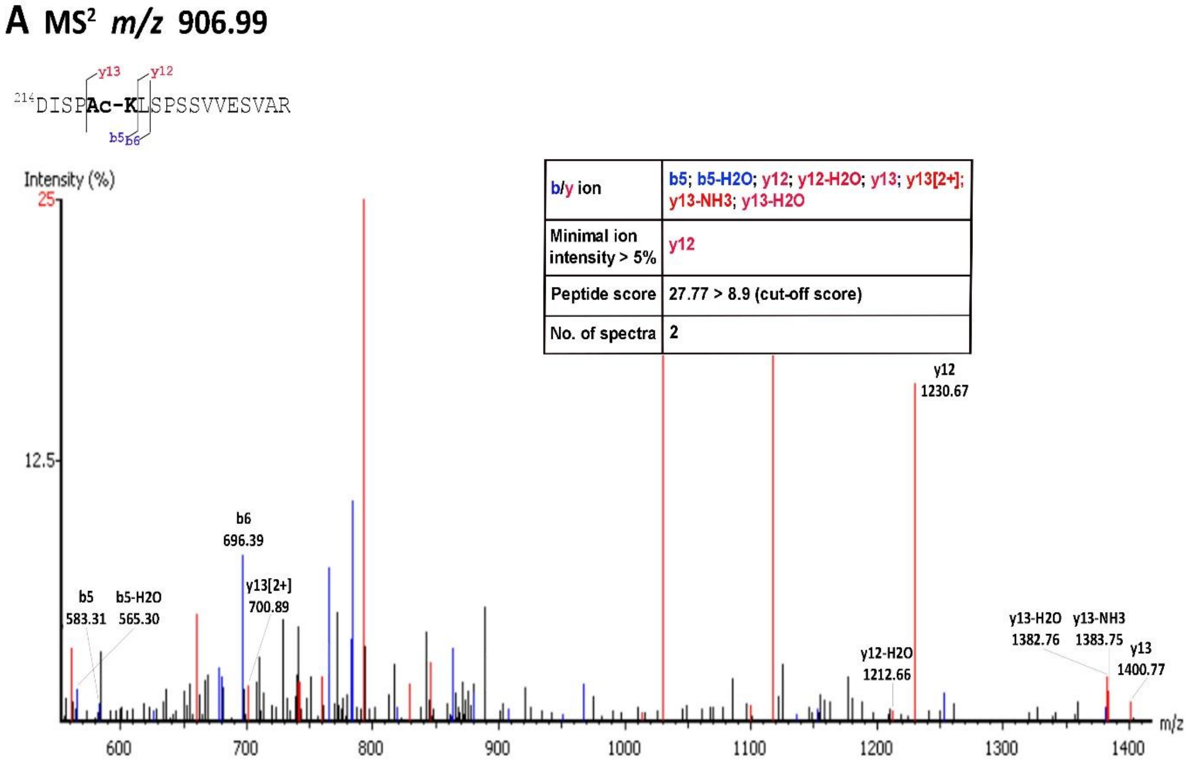

**B** MS$^2$ *m/z* 636.84

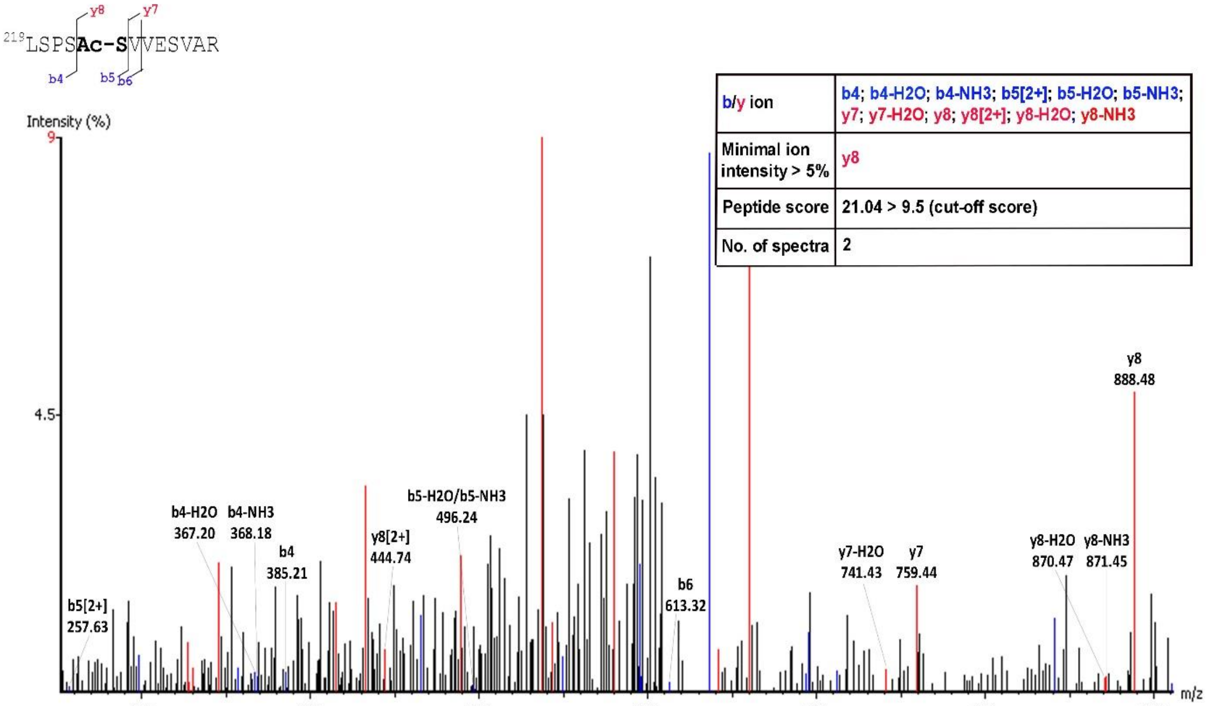


**C** MS$^2$ *m/z* 680.36

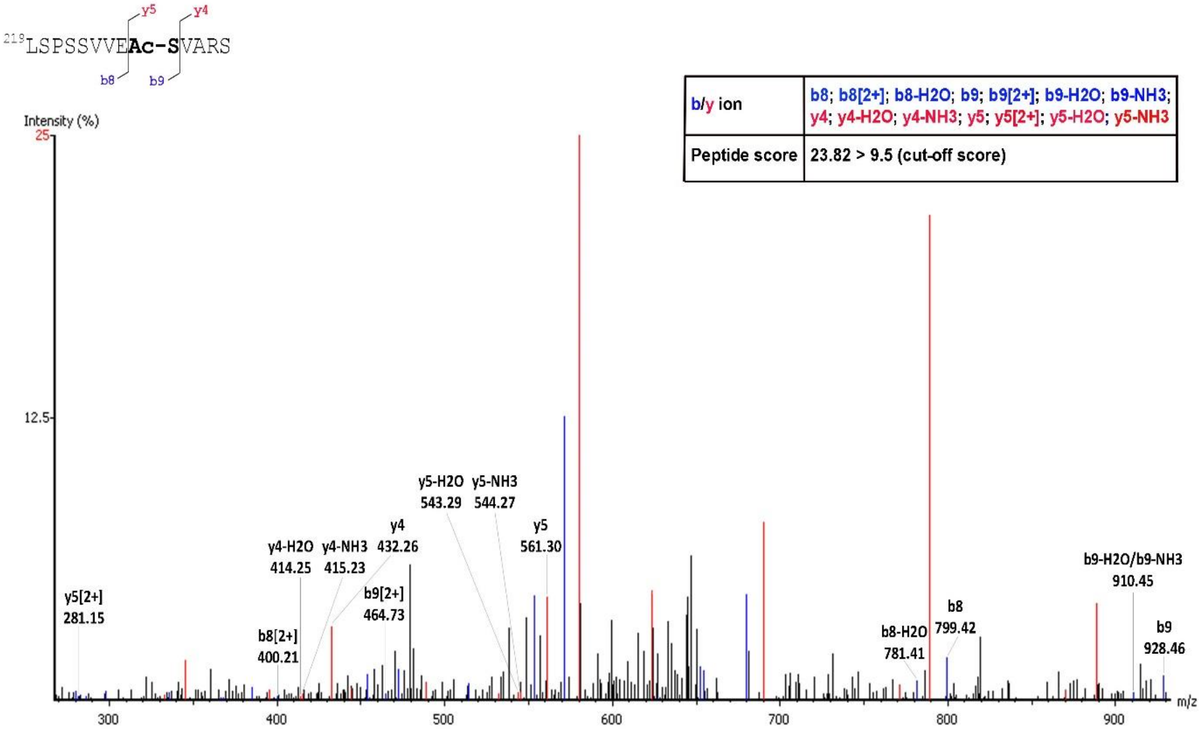

**D MS$^2$ *m/z* 758.38**

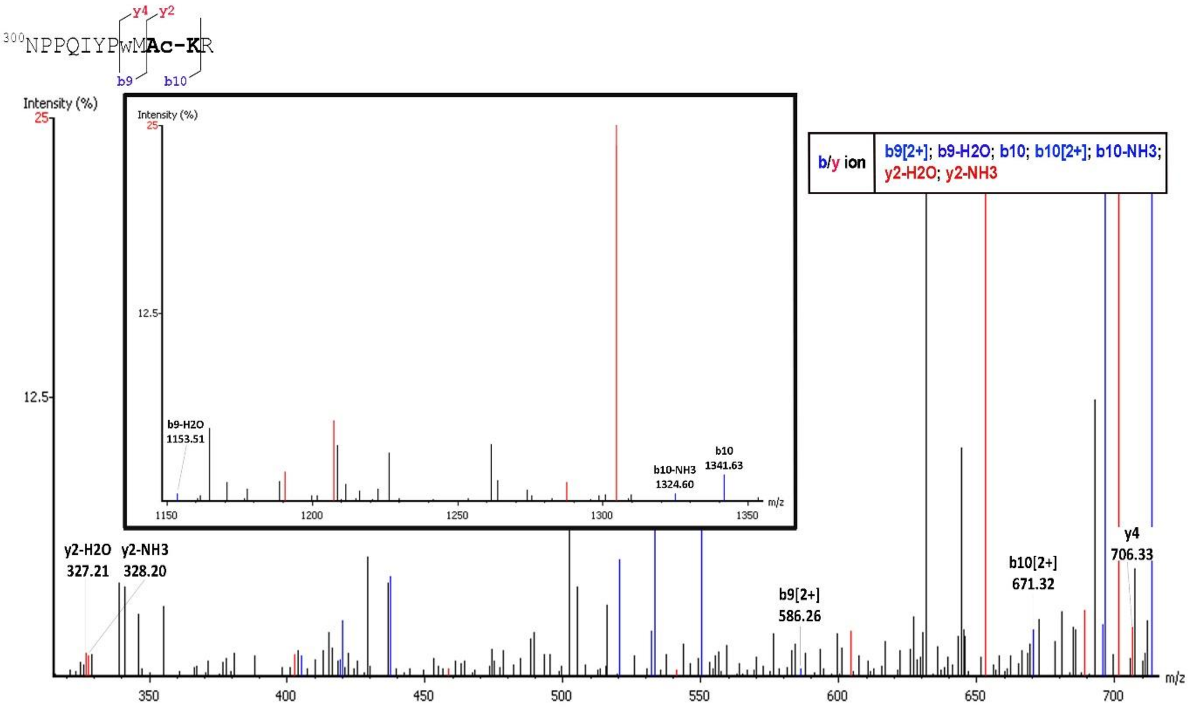


**E MS$^2$ *m/z* 706.3**

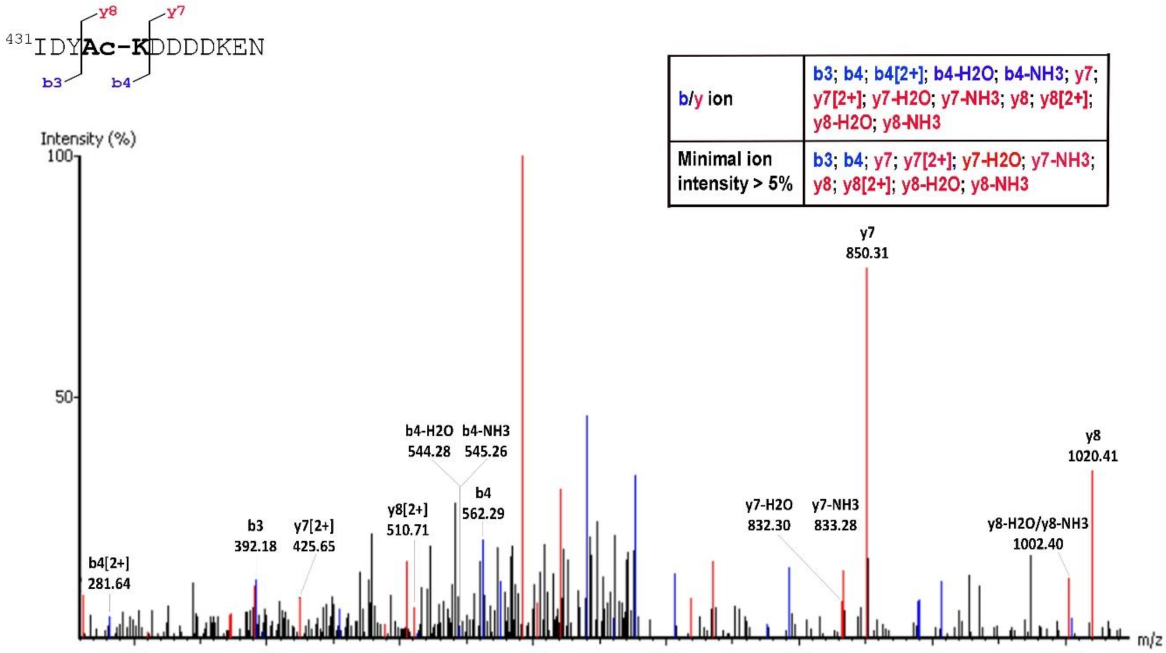

**F MS$^2$ *m/z* 706.3**

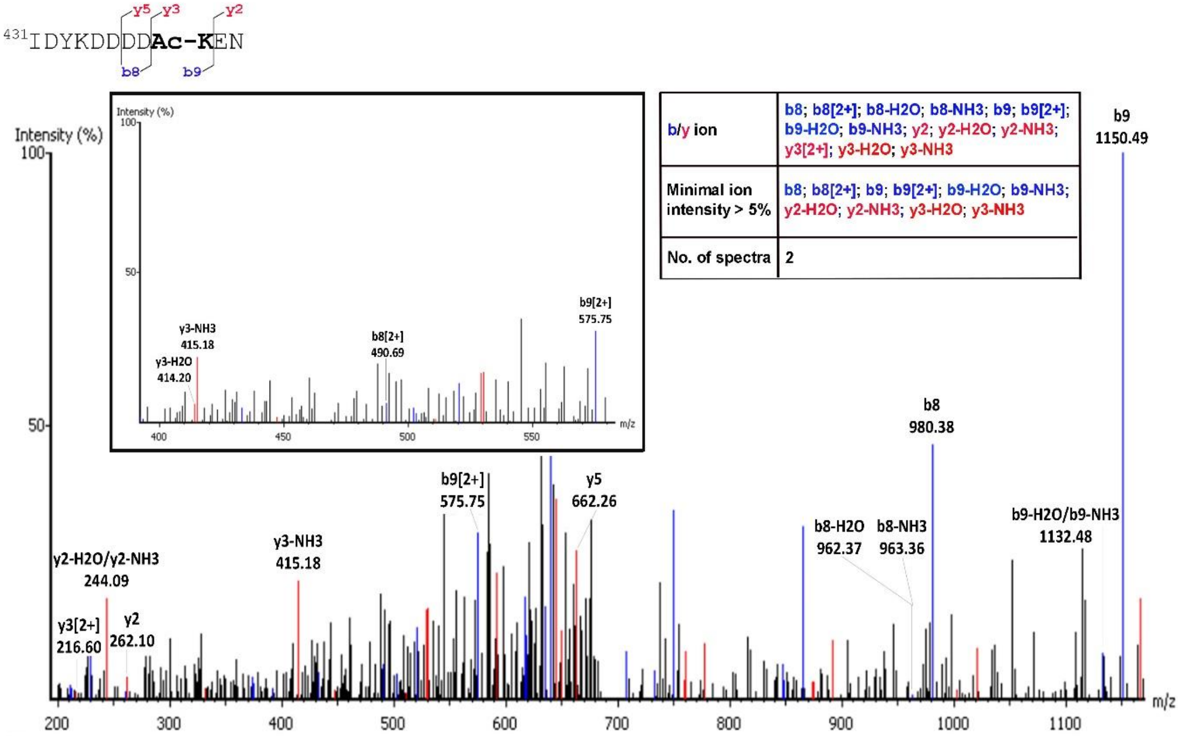


| | |
|---|---|
| b/y ion | b8; b8[2+]; b8-H2O; b8-NH3; b9; b9[2+]; b9-H2O; b9-NH3; y2; y2-H2O; y2-NH3; y3[2+]; y3-H2O; y3-NH3 |
| Minimal ion intensity > 5% | b8; b8[2+]; b9; b9[2+]; b9-H2O; b9-NH3; y2-H2O; y2-NH3; y3-H2O; y3-NH3 |
| No. of spectra | 2 |

**Figure S5. Formylation of Lysine 218, 309, 325, 341, 369, 434 and 439 residues of SCRTT.** MS$^2$ spectra of the peptide identified by LC-MS/MS is shown. A. Formylation of Lysine 218. The inset box shows fragment ions with *m/z* 300 to 700. B. Formylation of Lysine 309. The inset box shows fragment ions with *m/z* 1100 to 1300. C. Formylation of Lysine 325. The inset box shows fragment ions with *m/z* 700 to 1400. D. Formylation of Lysine 341. E. Formylation of Lysine 369. The inset box shows fragment ions with *m/z* 700 to 1100. F. Formylation of Lysine 434. The inset box shows fragment ions with *m/z* 830 to 1010. G. Formylation of Lysine 439. The inset box shows fragment ions with *m/z* 950 to 1200. The peptide sequence and *m/z* ratio are indicated at the top of the spectra. Positions of fragmentation are shown with vertical lines in the peptide sequence. The box on the right summarizes the evidence confirming formylation. The relevant fragment ions and their *m/z* ratios supporting formylation are labelled in the spectra.

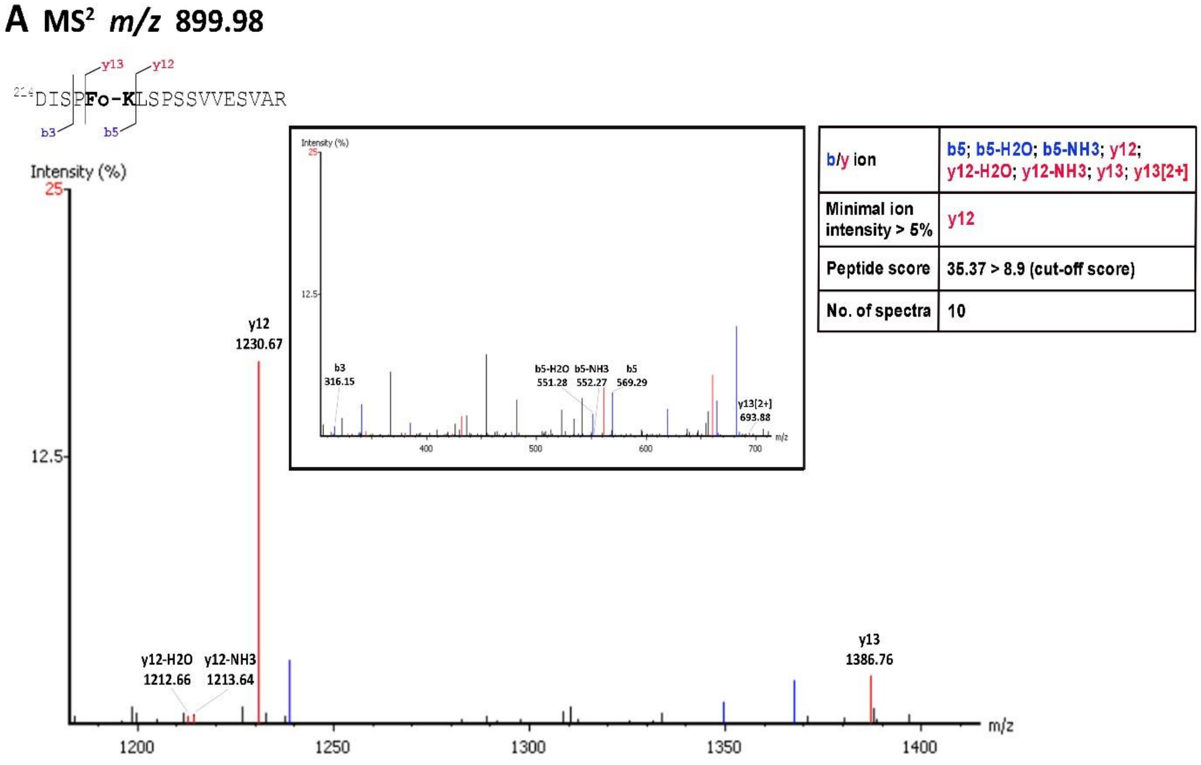

**B MS$^2$ *m/z* 737.37**

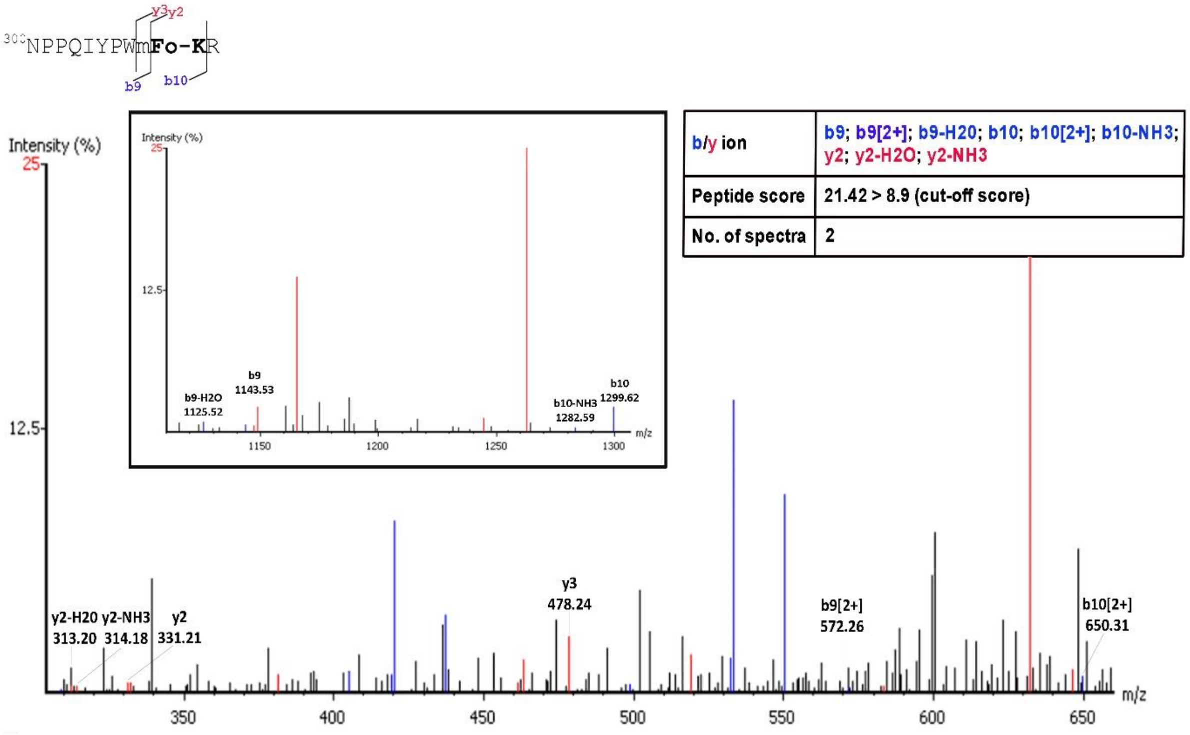


**C MS$^2$ *m/z* 571.3**

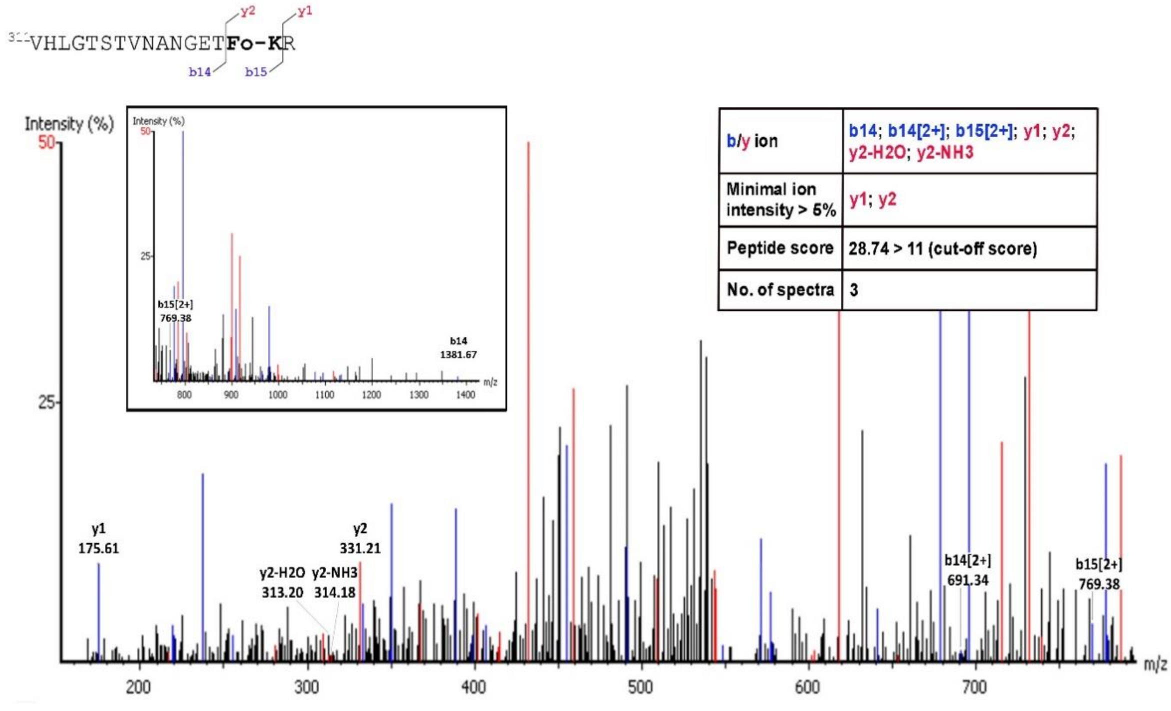

**D** MS[2] *m/z* 627.98

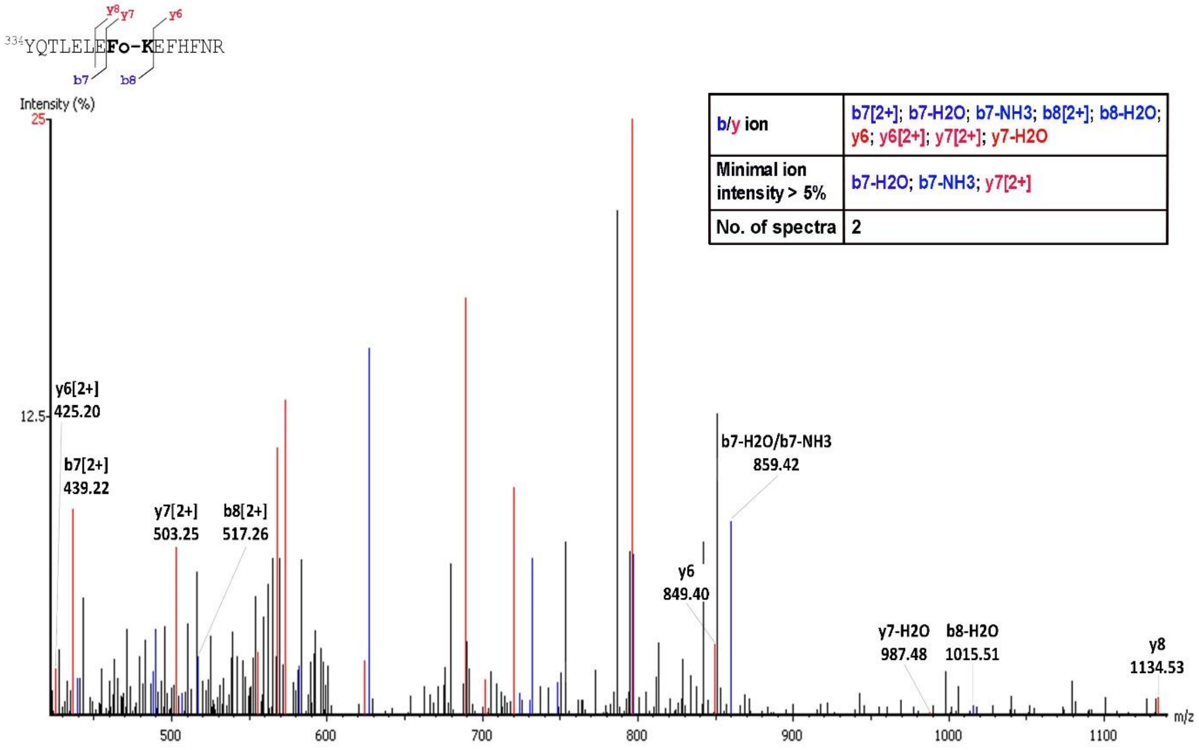


| b/y ion | b7[2+]; b7-H2O; b7-NH3; b8[2+]; b8-H2O; y6; y6[2+]; y7[2+]; y7-H2O |
|---|---|
| Minimal ion intensity > 5% | b7-H2O; b7-NH3; y7[2+] |
| No. of spectra | 2 |

**E** MS[2] *m/z* 630.85

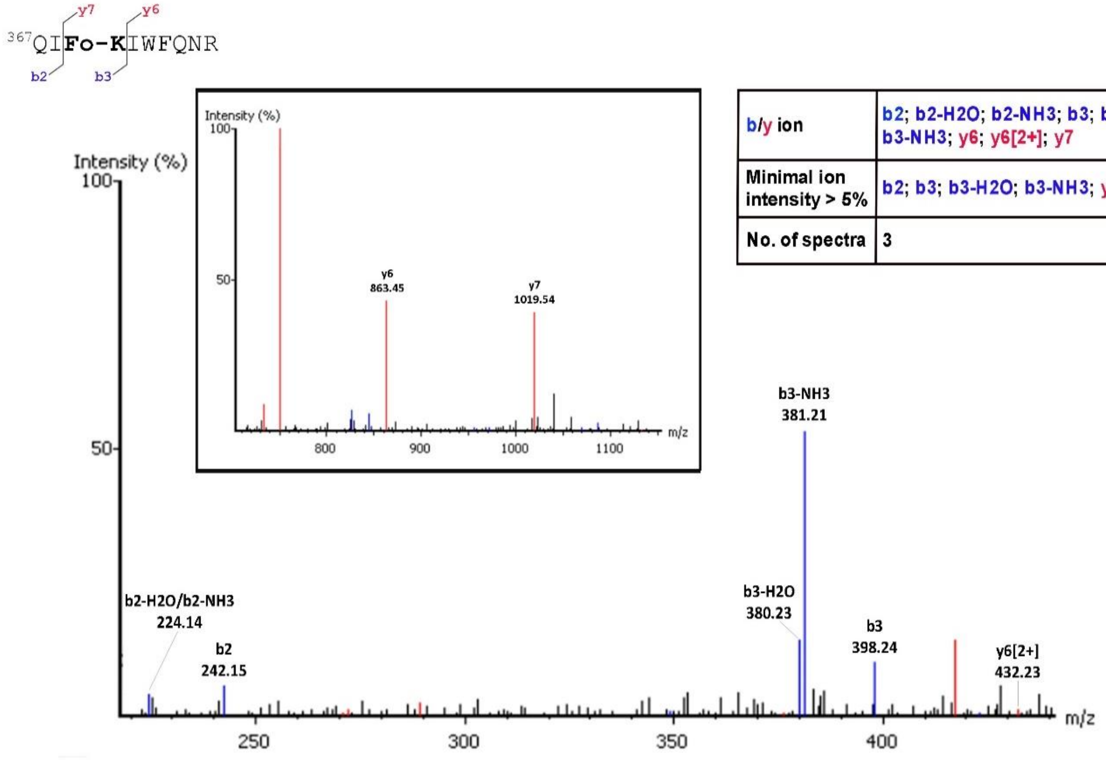


| b/y ion | b2; b2-H2O; b2-NH3; b3; b3-H2O; b3-NH3; y6; y6[2+]; y7 |
|---|---|
| Minimal ion intensity > 5% | b2; b3; b3-H2O; b3-NH3; y6; y7 |
| No. of spectra | 3 |

**F** MS[2] *m/z* 699.29

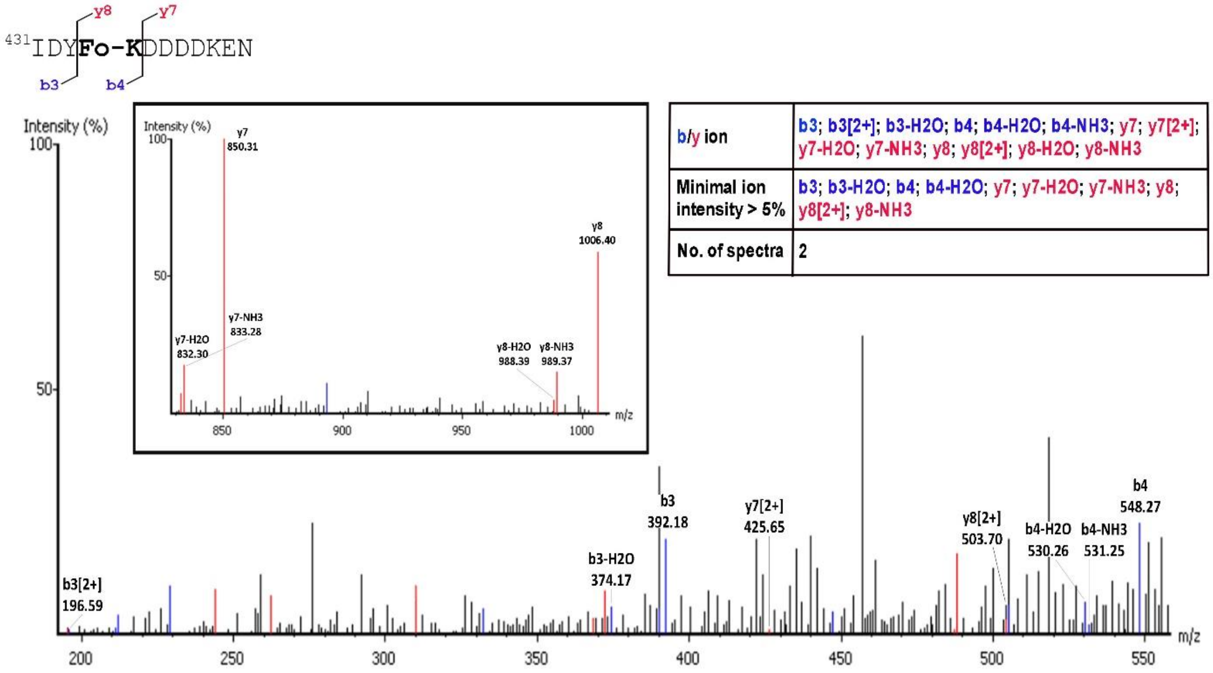


| b/y ion | b3; b3[2+]; b3-H2O; b4; b4-H2O; b4-NH3; y7; y7[2+]; y7-H2O; y7-NH3; y8; y8[2+]; y8-H2O; y8-NH3 |
|---|---|
| Minimal ion intensity > 5% | b3; b3-H2O; b4; b4-H2O; y7; y7-H2O; y7-NH3; y8; y8[2+]; y8-NH3 |
| No. of spectra | 2 |

**G** MS[2] *m/z* 699.29

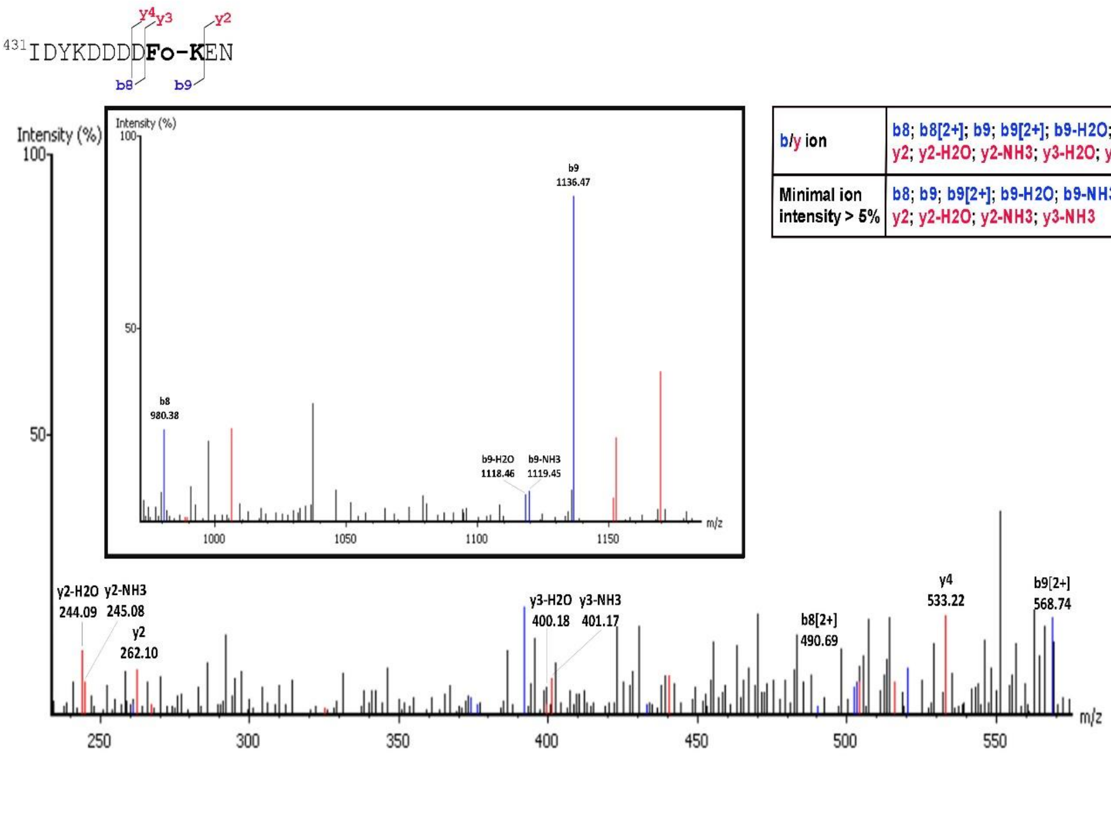


| b/y ion | b8; b8[2+]; b9; b9[2+]; b9-H2O; b9-NH3; y2; y2-H2O; y2-NH3; y3-H2O; y3-NH3 |
|---|---|
| Minimal ion intensity > 5% | b8; b9; b9[2+]; b9-H2O; b9-NH3; y2; y2-H2O; y2-NH3; y3-NH3 |

**Figure S6. Methylation of Serine 19, Serine 166, Lysine 168 and Threonine 364 residues of SCRTT.** $MS^2$ spectra of the peptide identified by LC-MS/MS is shown. A. Methylation of Lysine 19. B. Methylation of Serine 166. C. Methylation of Lysine 168. D. Methylation of Threonine 364. The inset box shows fragment ions with *m/z* 1000 to 1150. The peptide sequence and *m/z* ratio are indicated at the top of the spectra. Positions of fragmentation are shown with vertical lines in the peptide sequence. The box on the right summarizes the evidence confirming methylation. The relevant fragment ions and their *m/z* ratios supporting methylation are labelled in the spectra.

**A** $MS^2$ ***m/z* 714.34**

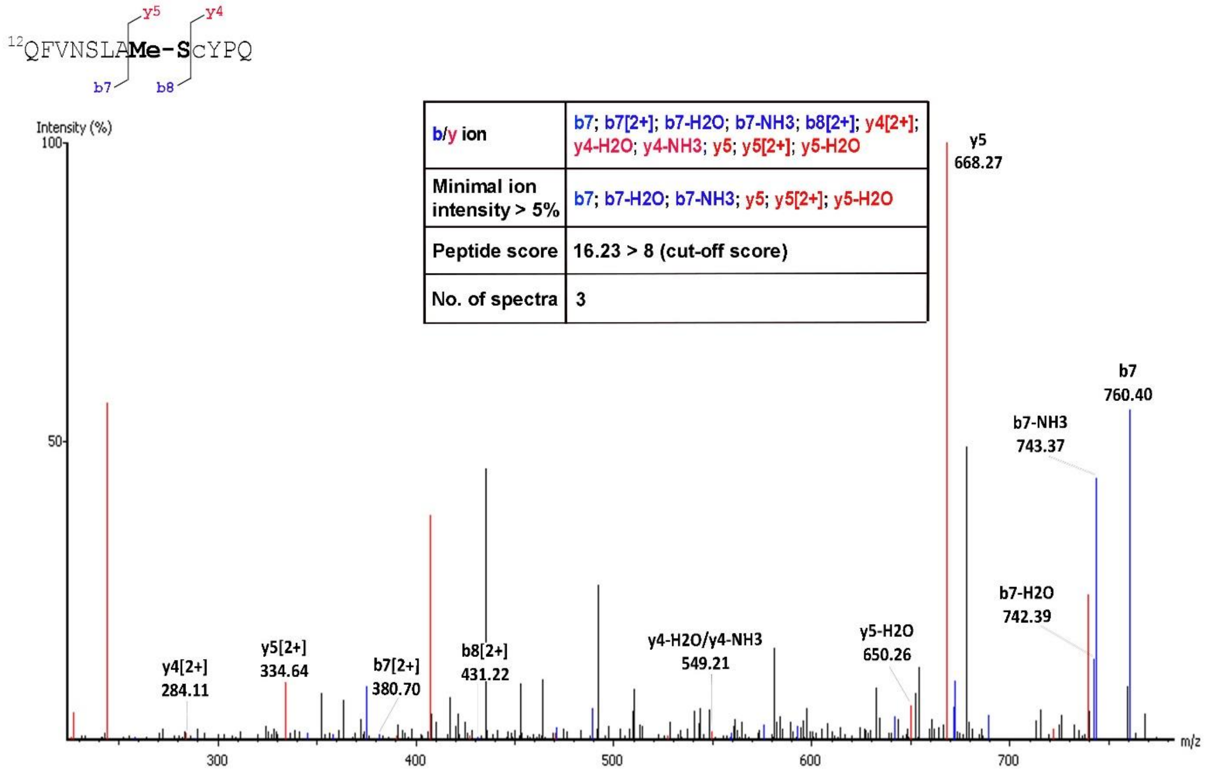

**B** MS$^2$ *m/z* 435.21

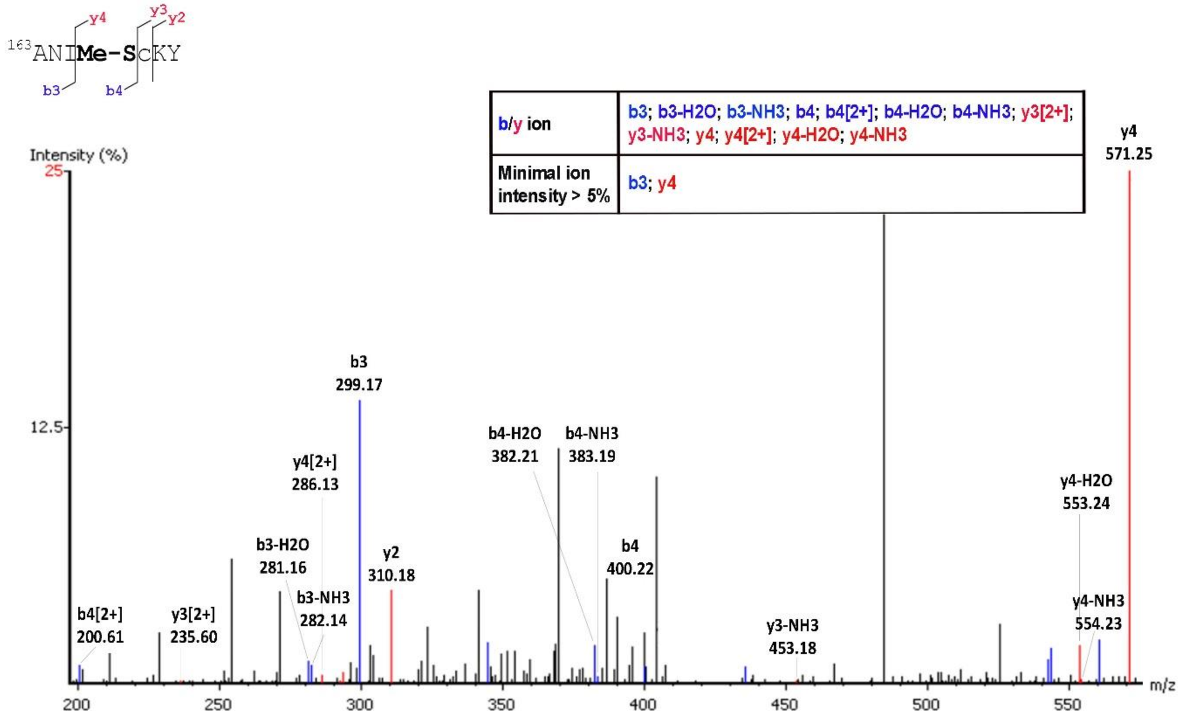


**C** MS$^2$ *m/z* 435.21

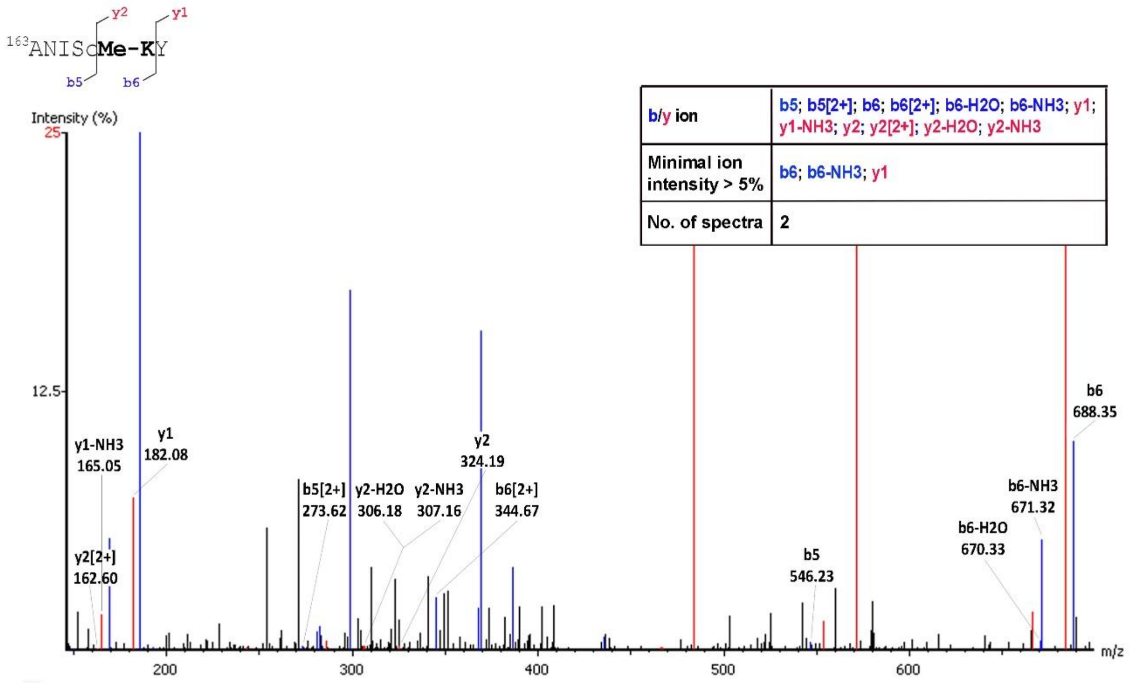

**D** MS$^2$ *m/z* 720.39

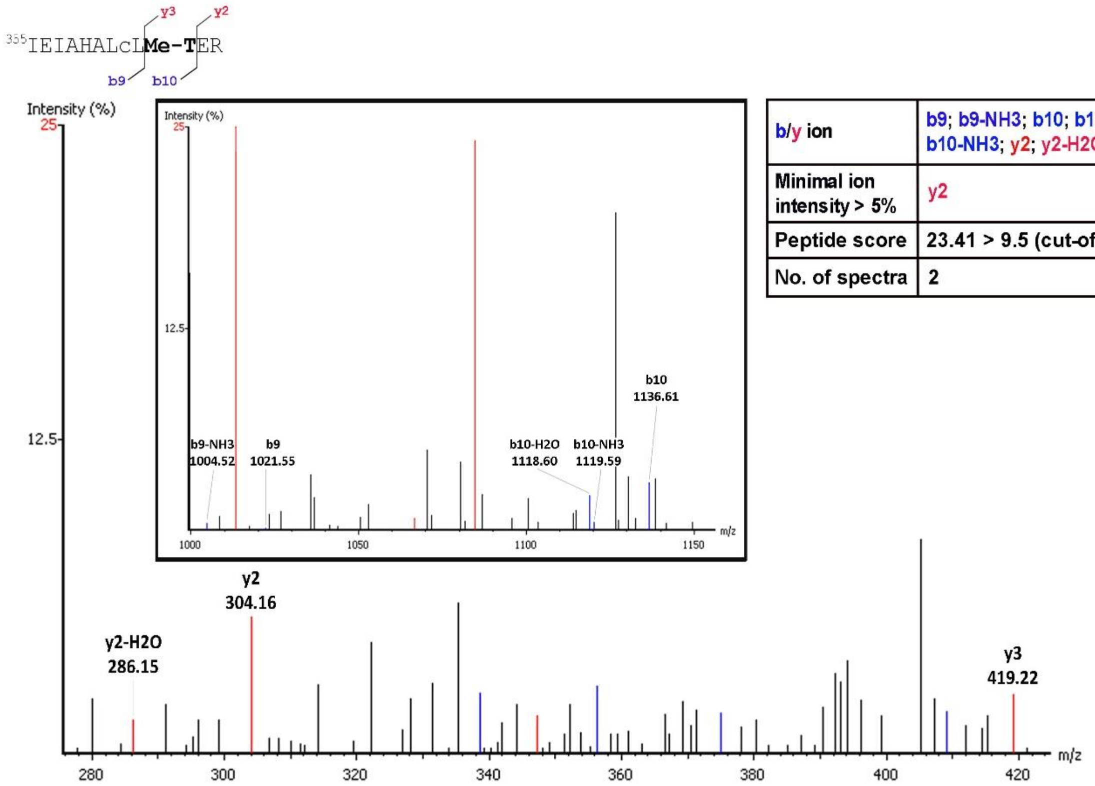


| b/y ion | b9; b9-NH3; b10; b10-H2O; b10-NH3; y2; y2-H2O; y3 |
|---|---|
| Minimal ion intensity > 5% | y2 |
| Peptide score | 23.41 > 9.5 (cut-off score) |
| No. of spectra | 2 |

**Figure S7. Carboxylation of Aspartic acid 108, Lysine 298, Tryptophan 307, Lysine 309, Glutamic acid 323, Lysine 325 and Lysine 369 residues of SCRTT.** $MS^2$ spectra of the peptide identified by LC-MS/MS is shown. A. Carboxylation of Aspartic acid 108. B. Carboxylation of Lysine 298. The inset box shows fragment ions with *m/z* 470 to 618. C. Carboxylation of Tryptophan 307. The inset box shows fragment ions with *m/z* 780 to 1030. D. Carboxylation of Lysine 309. The inset box shows fragment ions with *m/z* 990 to 1300. E. Carboxylation of Glutamic acid 323. F. Carboxylation of Lysine 325. G. Carboxylation of Lysine 369. The inset box shows fragment ions with *m/z* 840 to 1020. The peptide sequence and *m/z* ratio are indicated at the top of the spectra. Positions of fragmentation are shown with vertical lines in the peptide sequence. The box on the right summarizes the evidence confirming carboxylation. The relevant fragment ions and their *m/z* ratios supporting carboxylation are labelled in the spectra.

**A** $MS^2$ *m/z* **827.71**

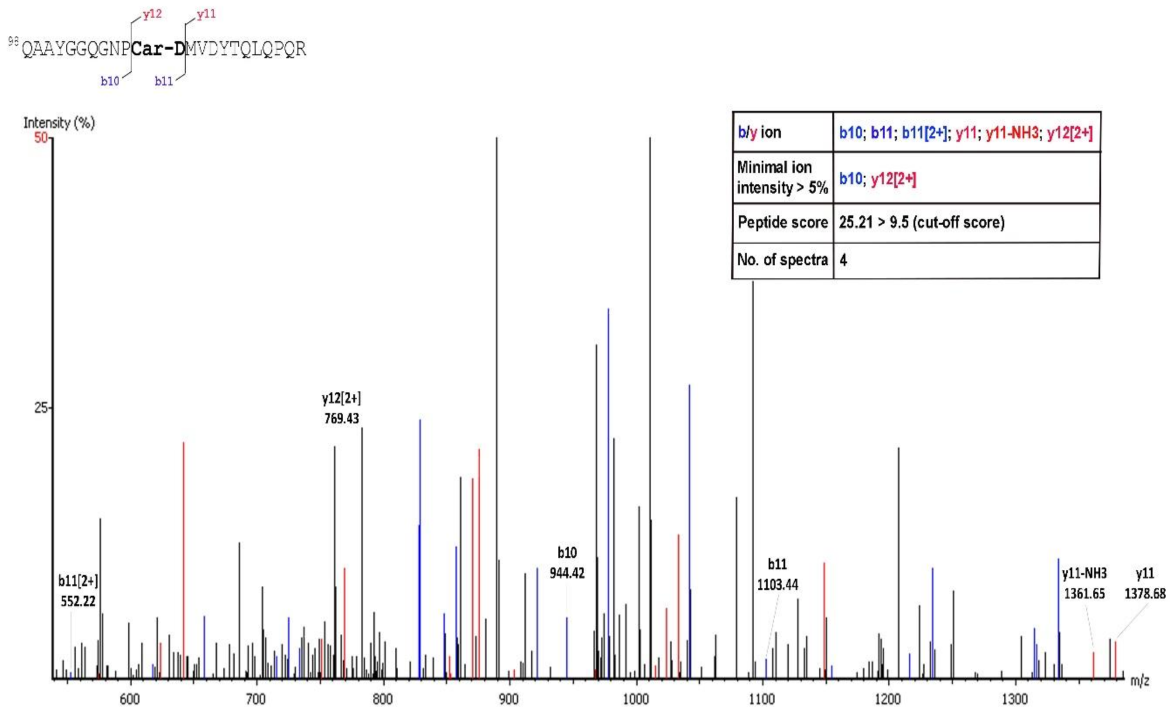

**B** MS$^2$ *m/z* 815.4

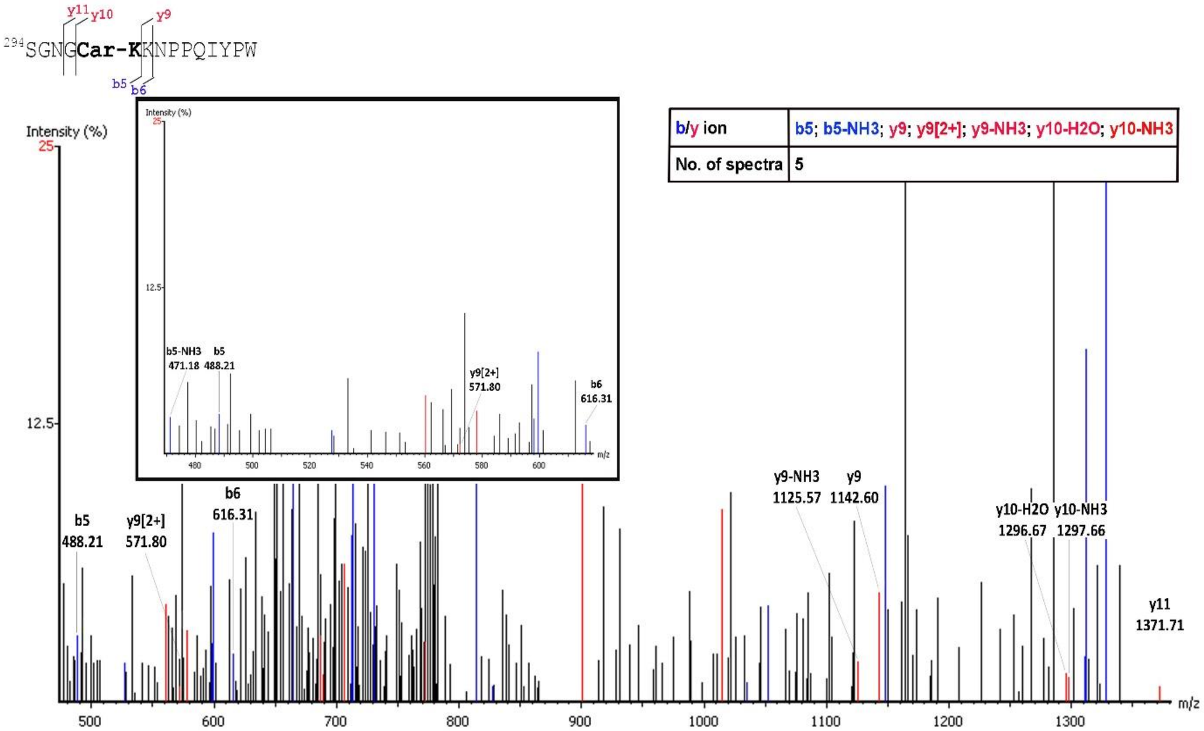


**C** MS$^2$ *m/z* 491.92

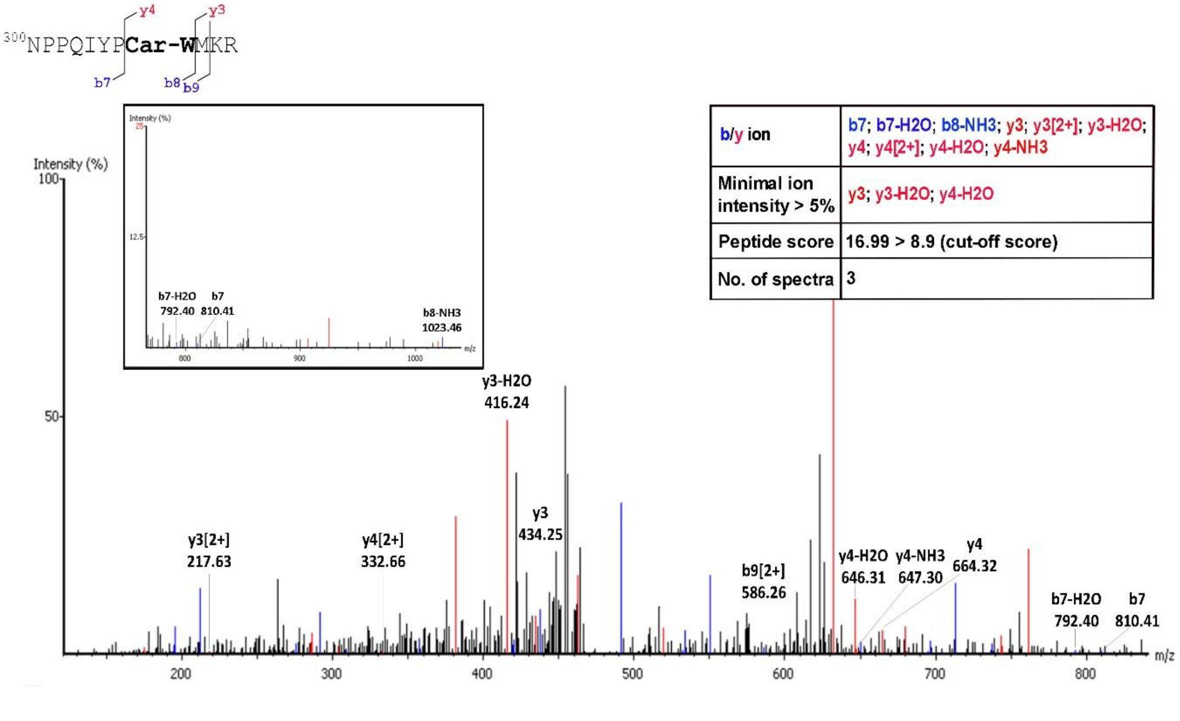

**D MS$^2$ *m/z* 737.37**

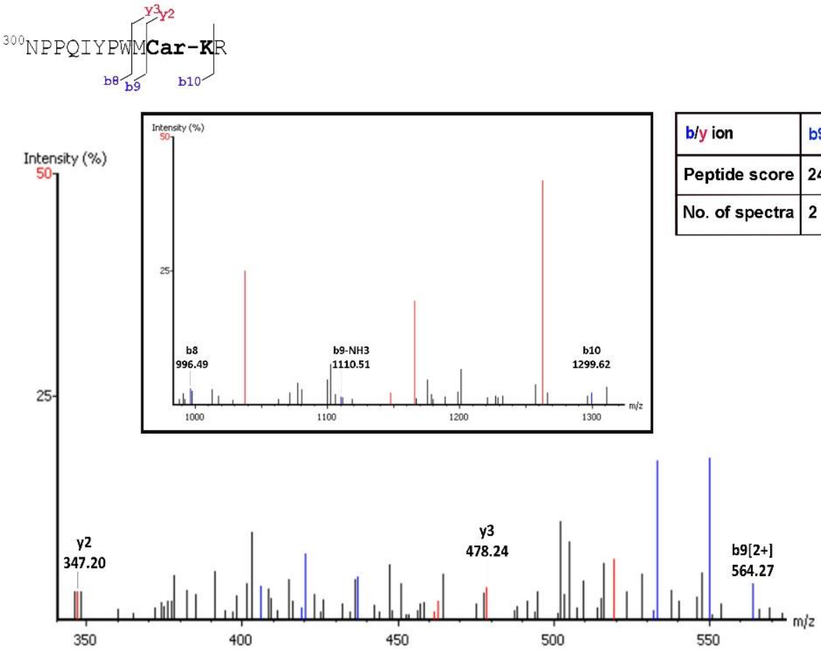


| b/y ion | b9[2+]; b9-NH3; b10; y2 |
|---|---|
| Peptide score | 24.22 > 11 (cut-off score) |
| No. of spectra | 2 |

**E MS$^2$ *m/z* 864.93**

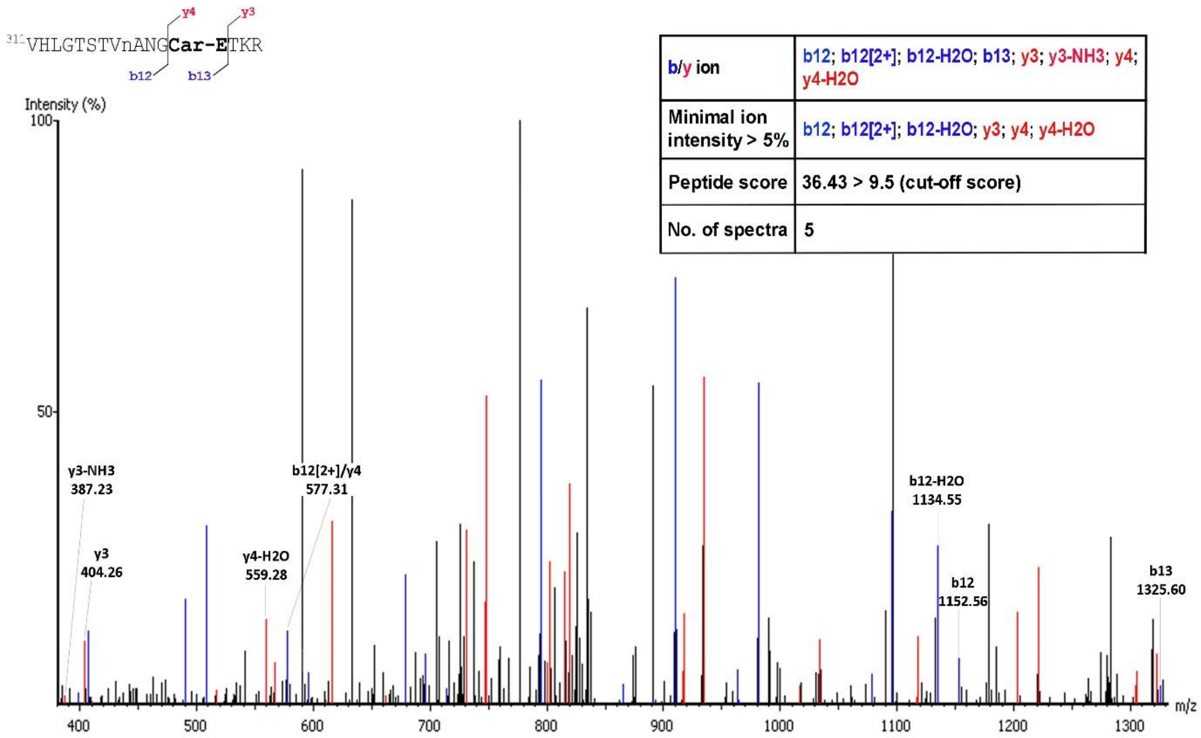


| b/y ion | b12; b12[2+]; b12-H2O; b13; y3; y3-NH3; y4; y4-H2O |
|---|---|
| Minimal ion intensity > 5% | b12; b12[2+]; b12-H2O; y3; y4; y4-H2O |
| Peptide score | 36.43 > 9.5 (cut-off score) |
| No. of spectra | 5 |

## F MS$^2$ *m/z* 595.63

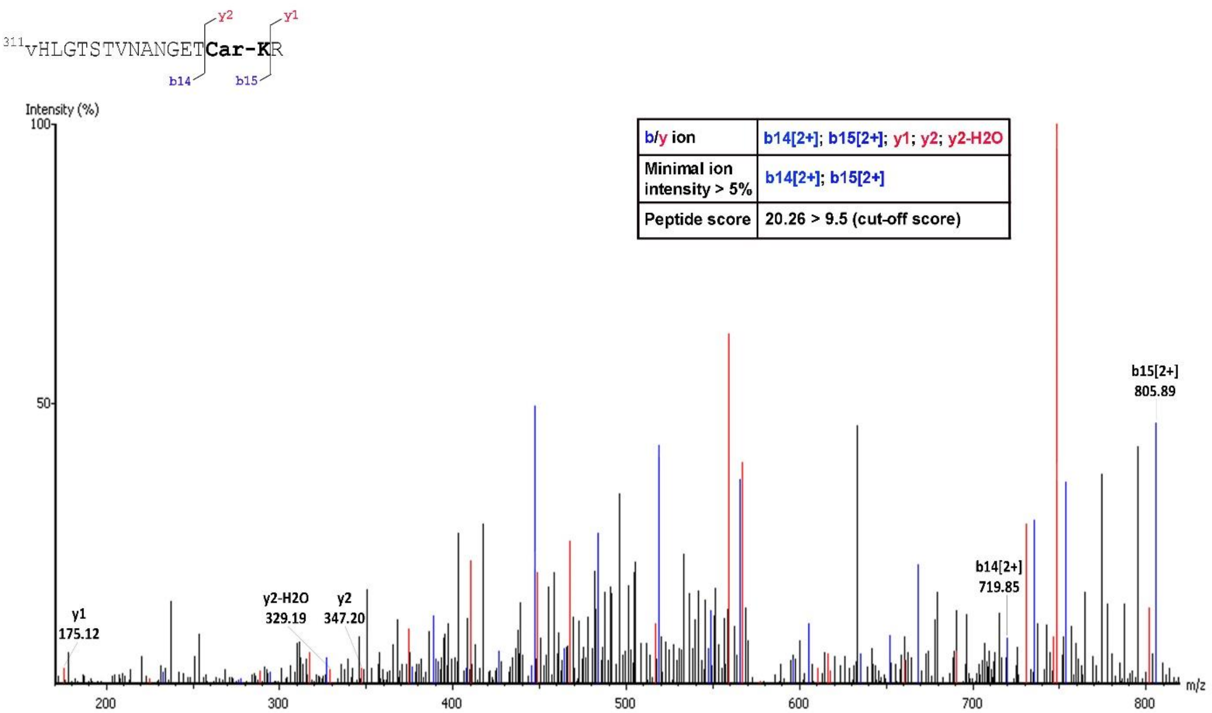


## G MS$^2$ *m/z* 638.85

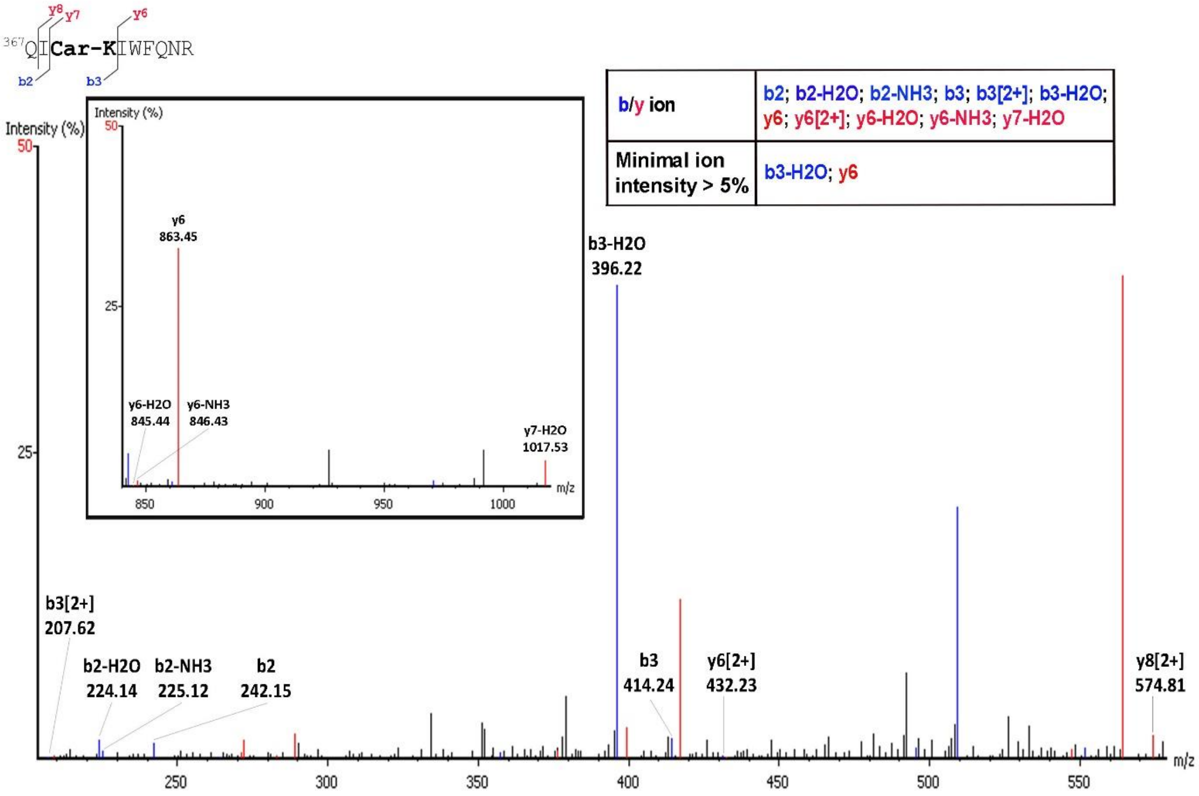

**Figure S8. Hydroxylation of Proline 22, Tyrosine 87, Proline 107, Aspartic Acid 108, Aspartic acid 111, Proline 269, Proline 306, Arginine 310, Asparagine 321, Lysine 325, Tyrosine 334, Arginine 366, Proline 392 and Tyrosine 398 residues of SCRTT.** $MS^2$ spectra of the peptide identified by LC-MS/MS is shown. A. Hydroxylation of Proline 22. The inset box shows fragment ions with *m/z* 835 to 920. B. Hydroxylation of Tyrosine 87. C. Hydroxylation of Proline 107. The inset box shows fragment ions with *m/z* 670 to 1000. D. Hydroxylation of Aspartic acid 108. The inset box shows fragment ions with *m/z* 620 to 900. E. Hydroxylation of Aspartic acid 111. The inset box shows fragment ions with *m/z* 420 to 600. F. Hydroxylation of Proline 269. The inset box shows fragment ions with *m/z* 420 to 940. G. Hydroxylation of Proline 306. H. Hydroxylation of R310. I. Hydroxylation of N321. J. Hydroxylation of K325. K. Hydroxylation of Y334. The inset box shows fragment ions with *m/z* 788 to 1040. L. Hydroxylation of R366. M. Hydroxylation of P392. The inset box shows fragment ions with *m/z* 750 to 890. N. Hydroxylation of Y398. The inset box shows fragment ions with *m/z* 945 to 1010. The peptide sequence and *m/z* ratio are indicated at the top of the spectra. Positions of fragmentation are shown with vertical lines in the peptide sequence. The box on the right summarizes the evidence confirming hydroxylation. The relevant fragment ions and their *m/z* ratios supporting hydroxylation are labelled in the spectra.

**A** $MS^2$ *m/z* **801.35**

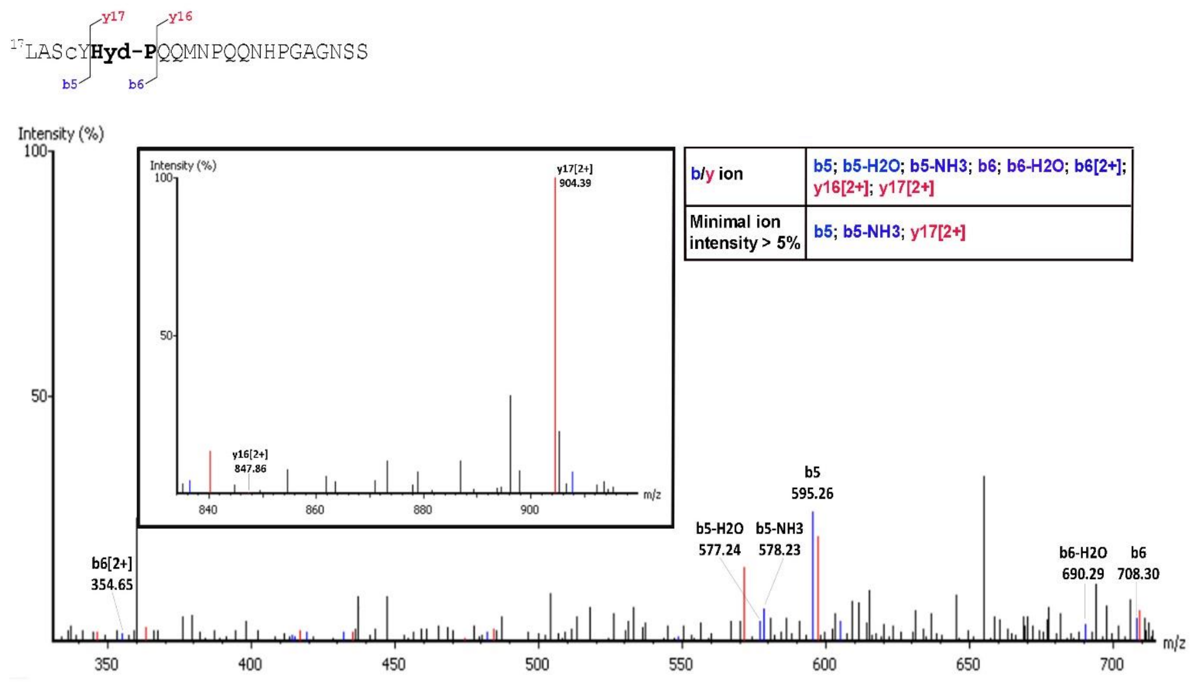

**B** MS[2] *m/z* 766.36

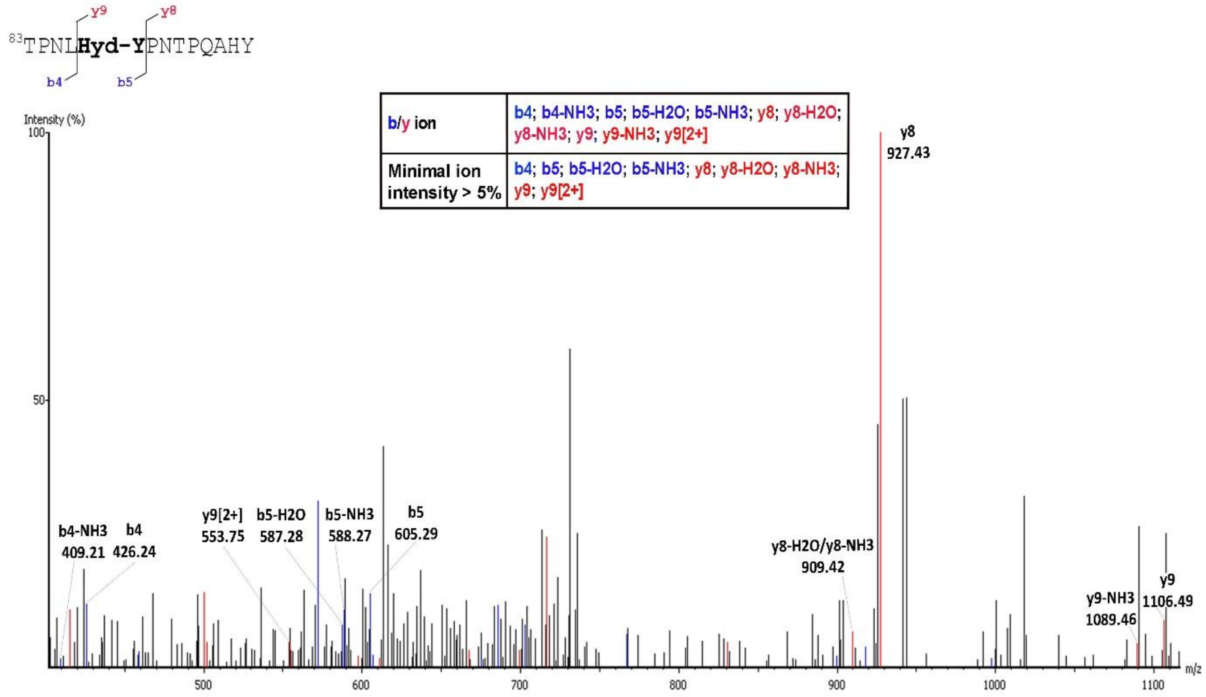


**C** MS[2] *m/z* 780.82

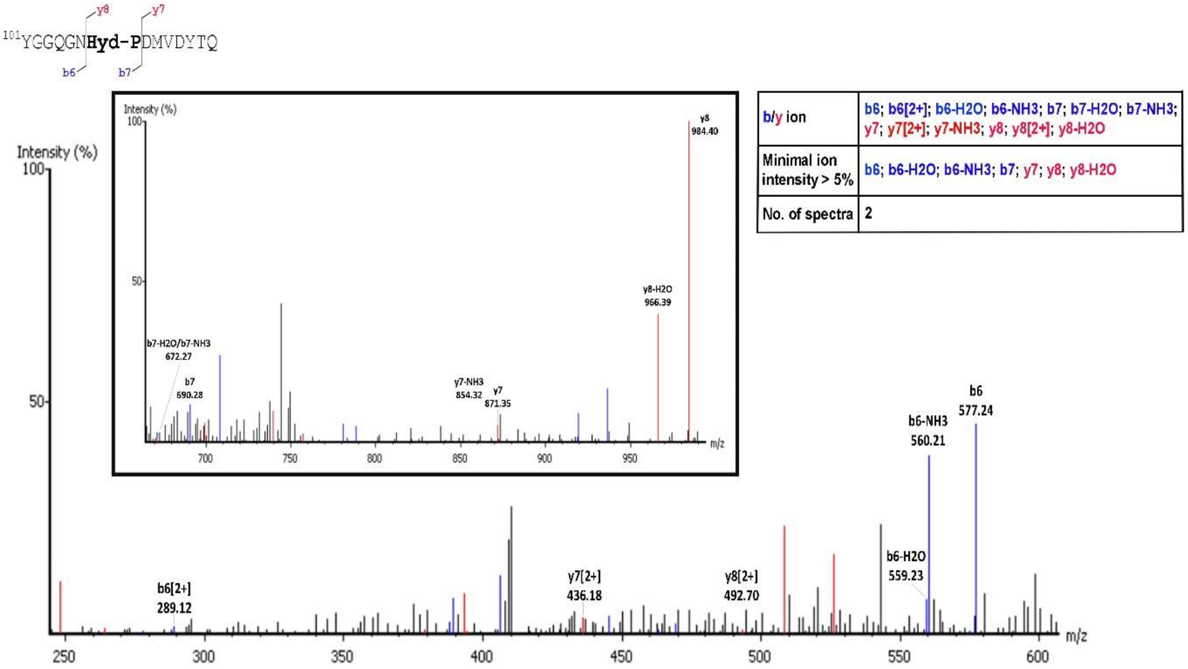

**D MS$^2$ *m/z* 699.29**

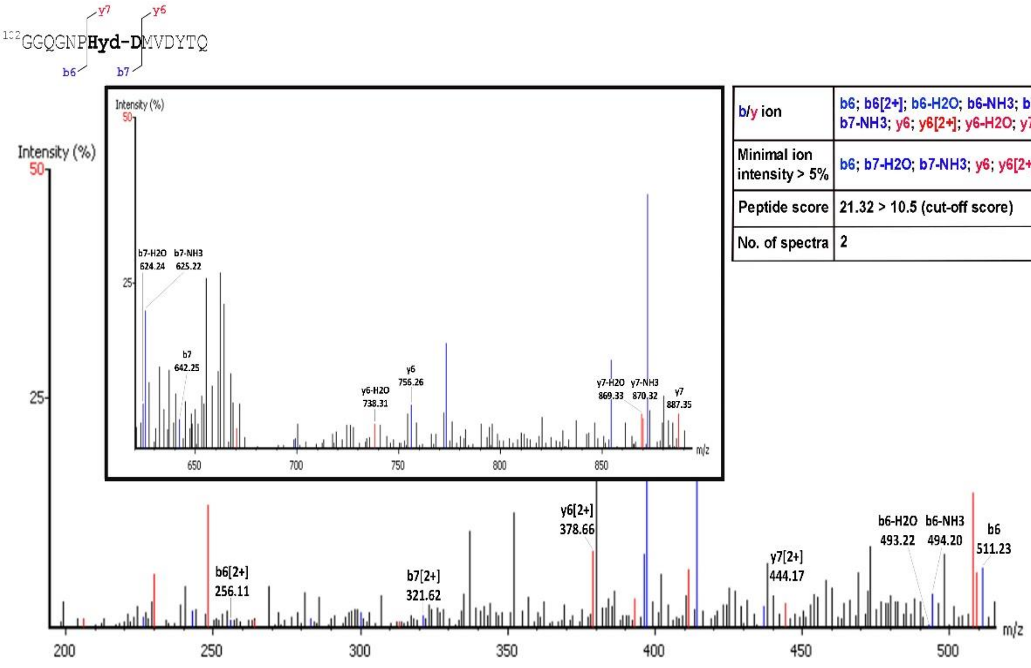


| b/y ion | b6; b6[2+]; b6-H2O; b6-NH3; b7; b7[2+]; b7-H2O; b7-NH3; y6; y6[2+]; y6-H2O; y7; y7[2+]; y7-H2O; y7-NH3 |
|---|---|
| Minimal ion intensity > 5% | b6; b7-H2O; b7-NH3; y6; y6[2+]; y7; y7-H2O |
| Peptide score | 21.32 > 10.5 (cut-off score) |
| No. of spectra | 2 |

**E MS$^2$ *m/z* 673.98**

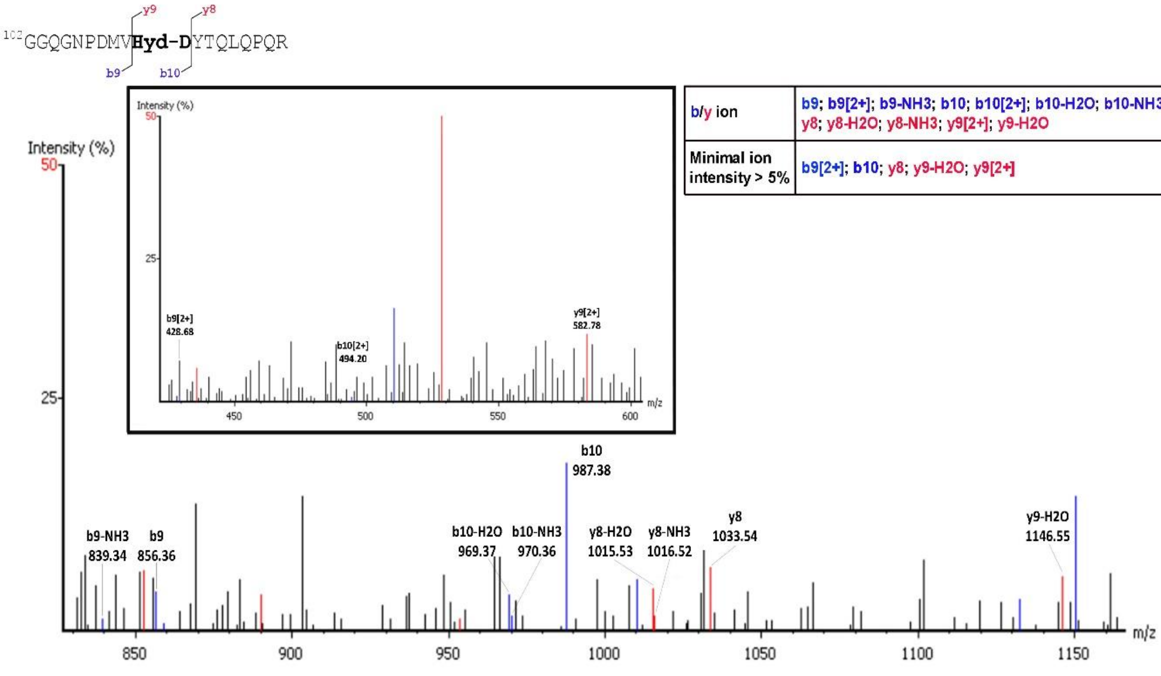


| b/y ion | b9; b9[2+]; b9-NH3; b10; b10[2+]; b10-H2O; b10-NH3; y8; y8-H2O; y8-NH3; y9[2+]; y9-H2O |
|---|---|
| Minimal ion intensity > 5% | b9[2+]; b10; y8; y9-H2O; y9[2+] |

**F MS² *m/z* 1095.43**

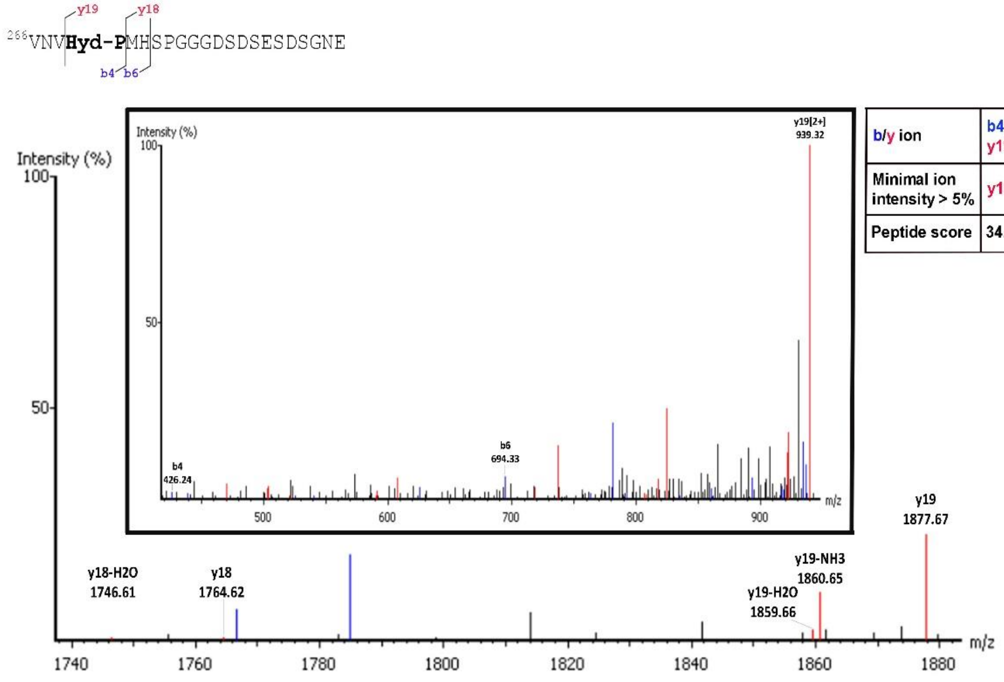


| b/y ion | b4; y18; y18-H2O; y19; y19[2+]; y19-H2O; y19-NH3 |
|---|---|
| Minimal ion intensity > 5% | y19; y19[2+]; y19-NH3 |
| Peptide score | 34.16 > 17.9 (cut-off score) |

**G MS² *m/z* 645.32**

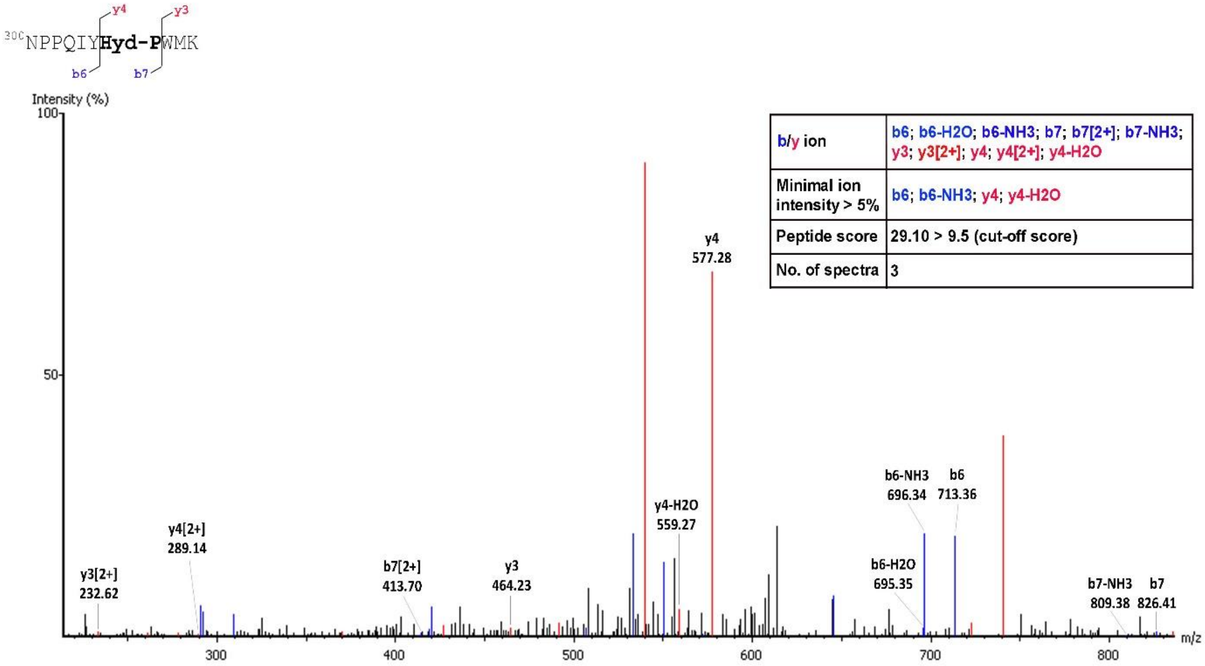


| b/y ion | b6; b6-H2O; b6-NH3; b7; b7[2+]; b7-NH3; y3; y3[2+]; y4; y4[2+]; y4-H2O |
|---|---|
| Minimal ion intensity > 5% | b6; b6-NH3; y4; y4-H2O |
| Peptide score | 29.10 > 9.5 (cut-off score) |
| No. of spectra | 3 |

**H** MS$^2$ *m/z* 482.58

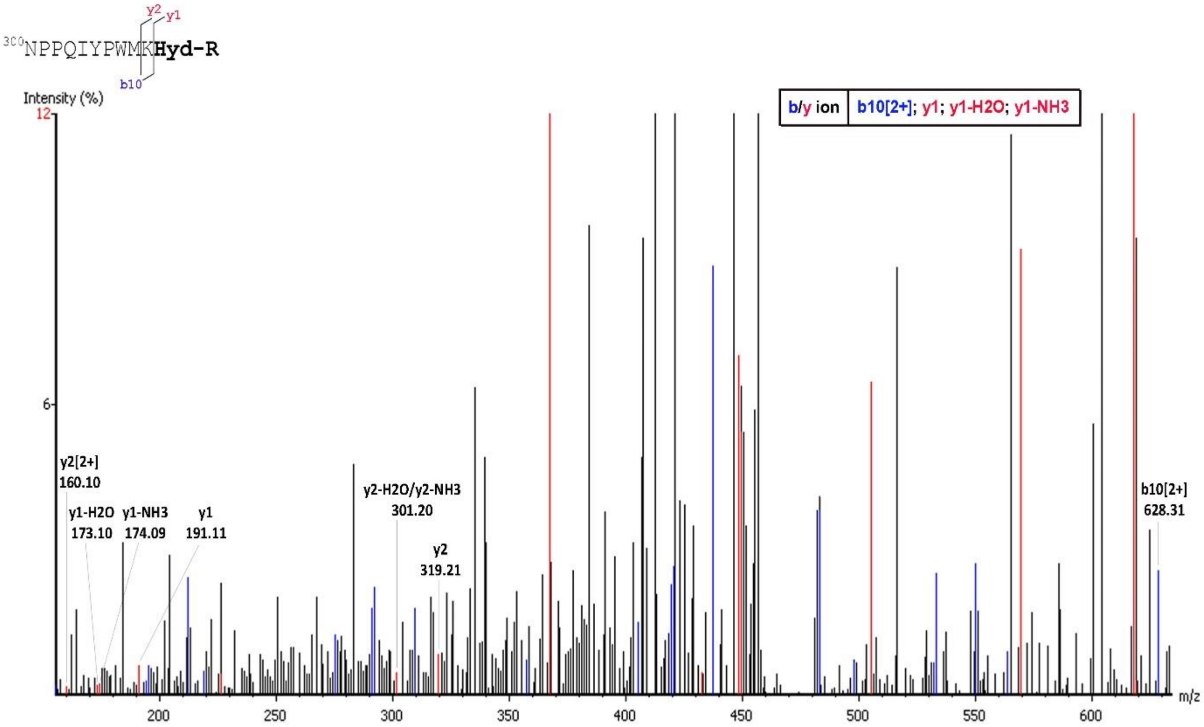


**I** MS$^2$ *m/z* 586.3

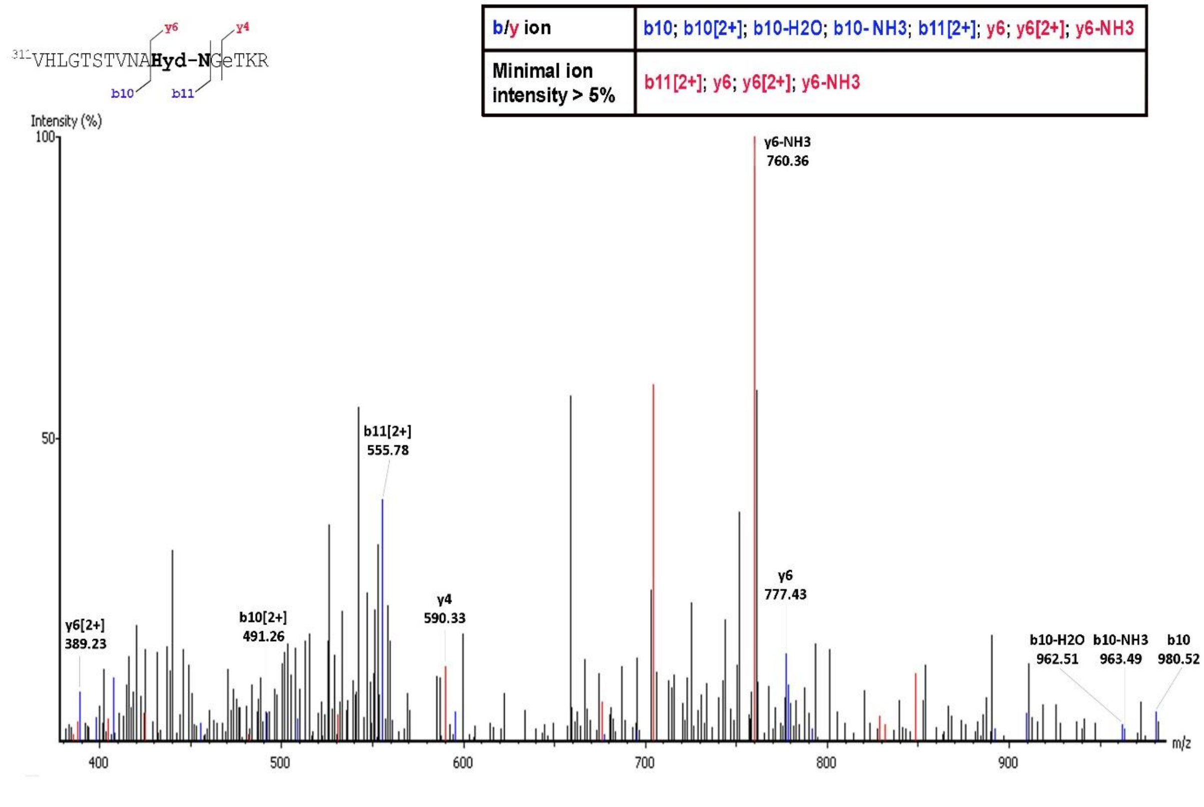

**J** MS[2] *m/z* 567.62

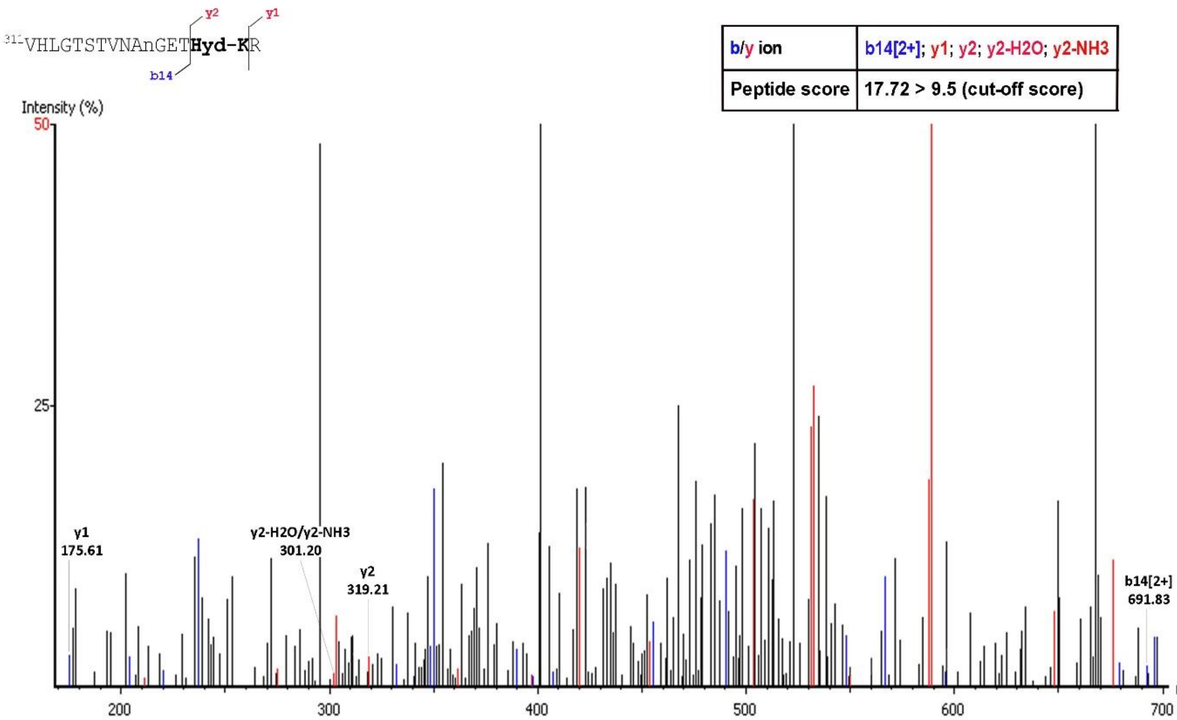


**K** MS[2] *m/z* 549.95

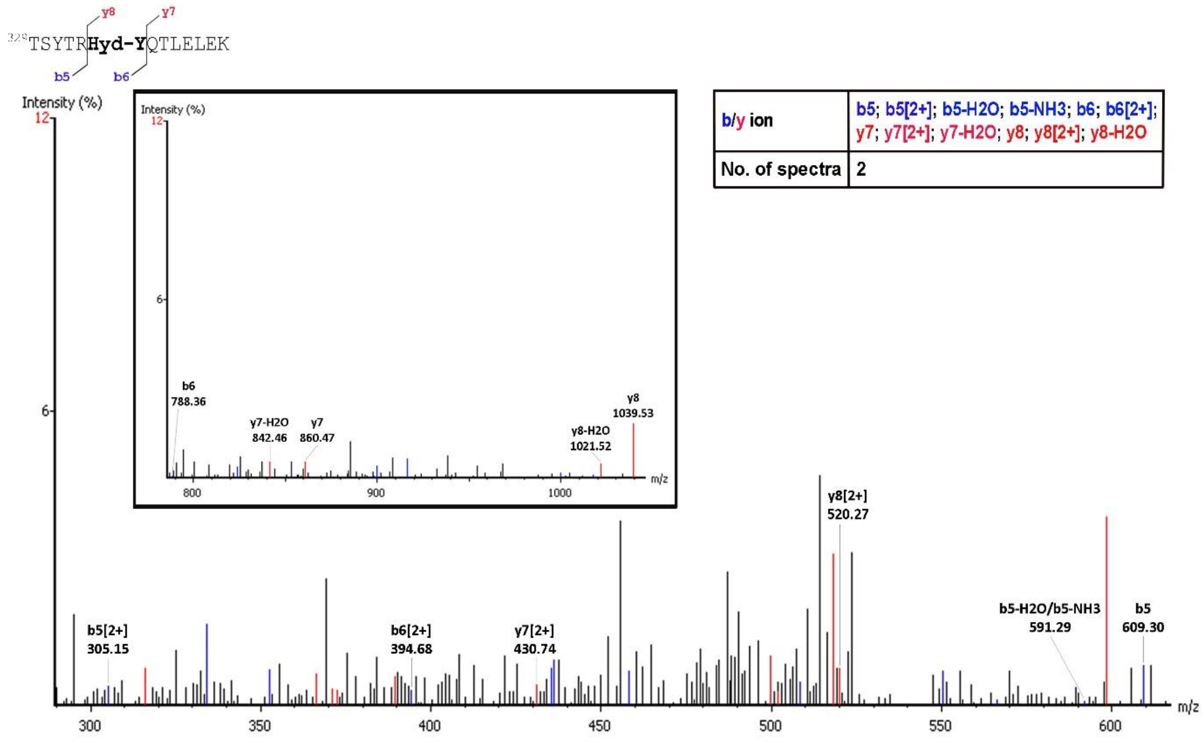

**L** MS$^2$ *m/z* 481.25

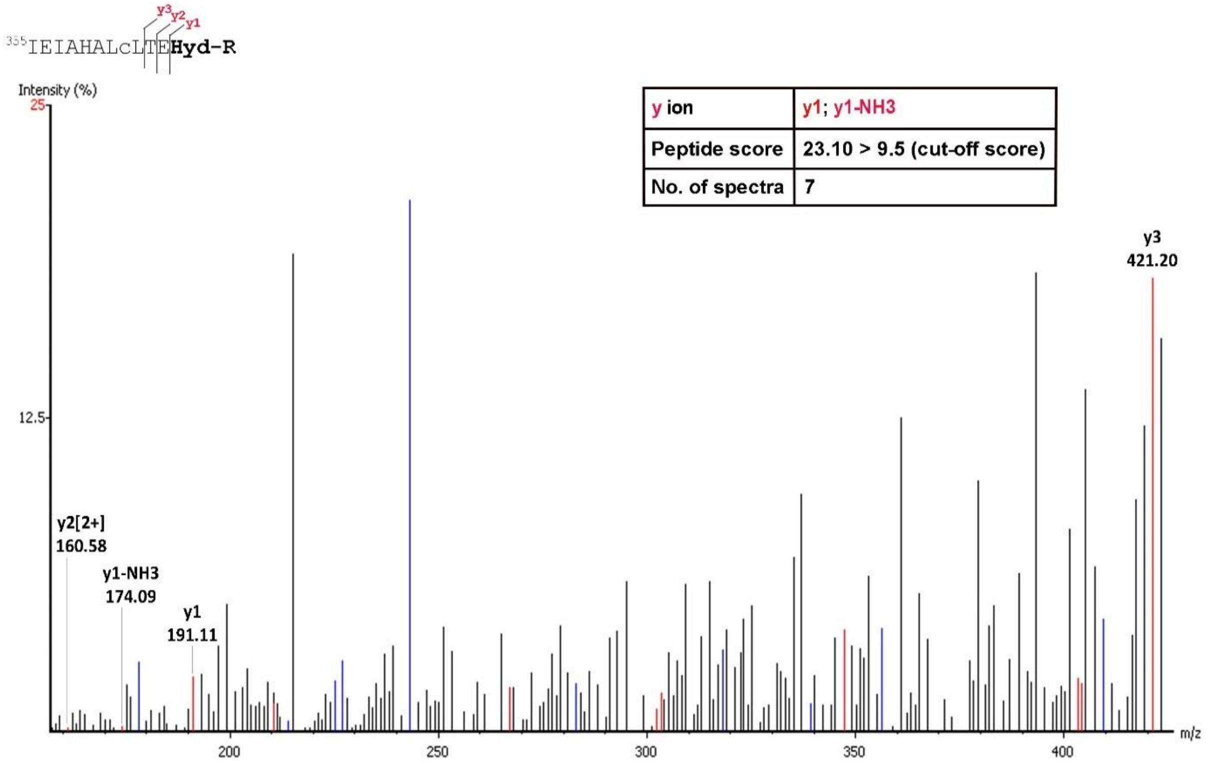


| | |
|---|---|
| y ion | y1; y1-NH3 |
| Peptide score | 23.10 > 9.5 (cut-off score) |
| No. of spectra | 7 |

**M** MS$^2$ *m/z* 603.78

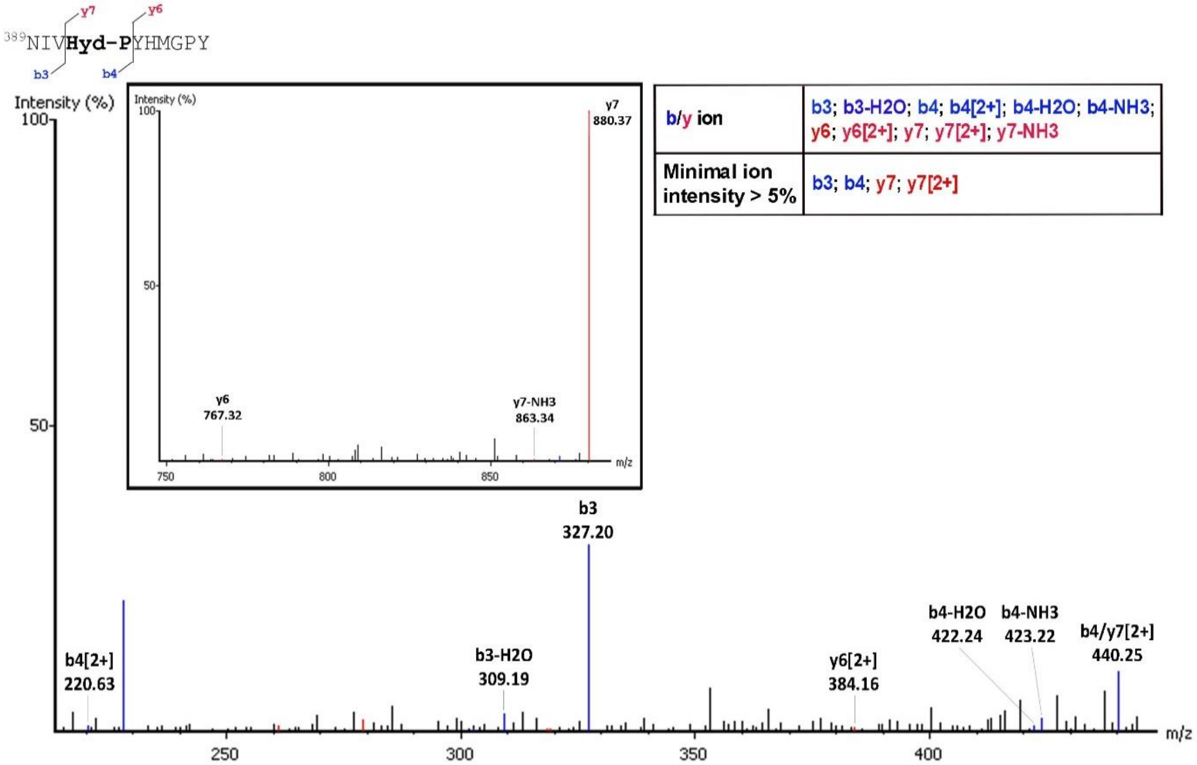


| | |
|---|---|
| b/y ion | b3; b3-H2O; b4; b4[2+]; b4-H2O; b4-NH3; y6; y6[2+]; y7; y7[2+]; y7-NH3 |
| Minimal ion intensity > 5% | b3; b4; y7; y7[2+] |

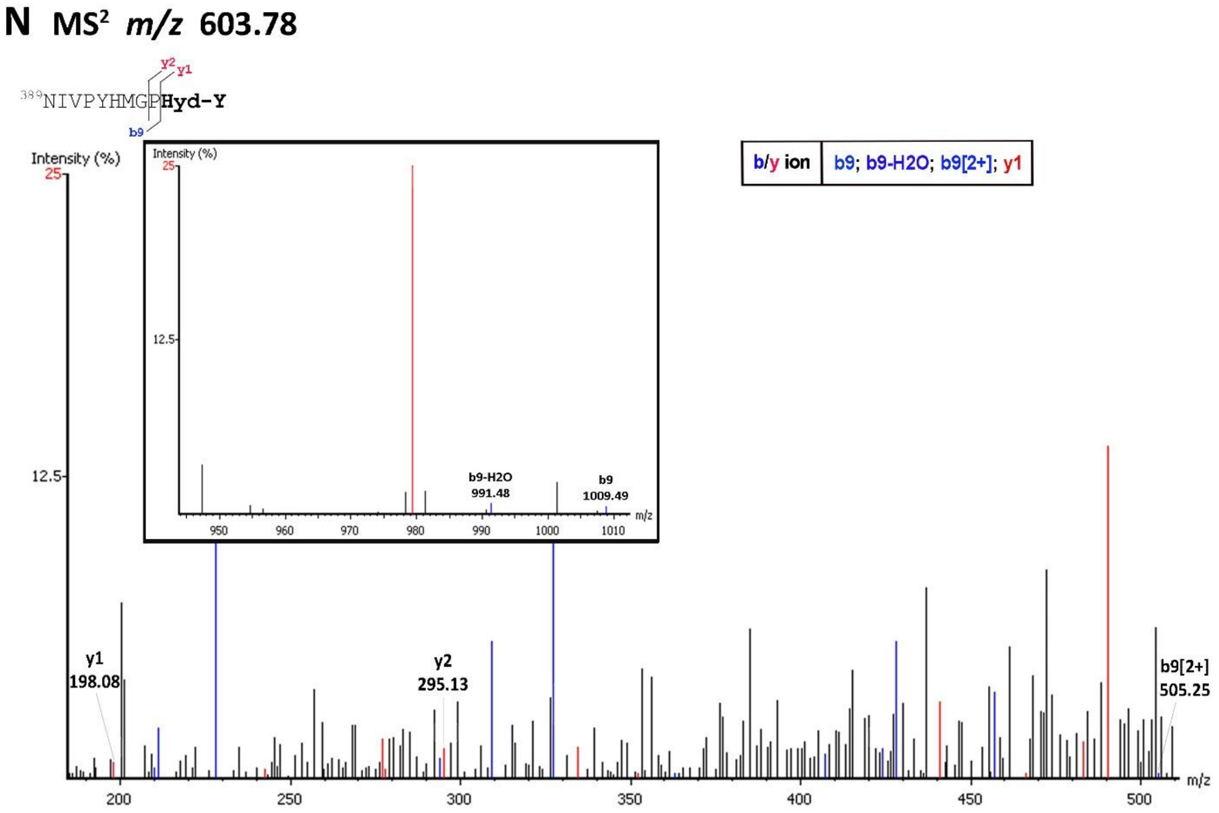
N MS2 m/z 603.78
389NIVPYHMGPHyd-Y
y2
y1
b9
Intensity (%)
25
12.5
b/y ion
b9; b9-H2O; b9[2+]; y1
b9-H2O
991.48
b9
1009.49
950
960
970
980
990
1000
1010
m/z
y1
198.08
y2
295.13
b9[2+]
505.25
200
250
300
350
400
450
500
m/z

**Table S1. Post-translational modifications of embryonic SCRTT identified by LC-MS/MS**

| Location | Peptide | n* | Protease | Site modified | Modification |
|---|---|---|---|---|---|
| 12-23 | Y.QFVNSLAS(+14.02)C(+57.02)YPQ.Q | 1 | Chymotrypsin | S19 | Methylation |
| 17-24 | S.LAS(+14.02)C(+57.02)YPQQ.M | 1 | Thermolysin | S19 | Methylation |
| 17-38 | S.LASC(+57.02)YP(+15.99)QQMNPQQNHPGAGNSS.A | 1 | Thermolysin | P22 | Hydroxylation |
| 83-95 | Y.TPNLY(+15.99)PNTPQAHY.A | 1 | Chymotrypsin | Y87 | Hydroxylation |
| 86-119 | N.LYPNTPQAHYANQAAYGGQGNPD(+15.99)MVDYTQLQPQR.L | 1 | Trypsin | D108 | Hydroxylation |
| 95-119 | H.YANQAAYGGQGNP(+15.99)DMVDYTQLQPQR.L | 4 | Trypsin | P107 | Hydroxylation |
| 95-119 | H.YANQAAYGGQGNPD(+15.99)MVDYTQLQPQR.L | 4 | Trypsin | D108 | Hydroxylation |
| 95-119 | H.YANQAAYGGQGNPD(+43.99)MVDYTQLQPQR.L | 1 | Trypsin | D108 | Carboxylation |
| 96-112 | Y.ANQAAYGGQGNP(+15.99)DMVDY.T | 1 | Chymotrypsin | P107 | Hydroxylation |
| 98-119 | N.QAAYGGQGNP(+15.99)DMVDYTQLQPQR.L | 1 | Trypsin | P107 | Hydroxylation |
| 98-119 | N.QAAYGGQGNPD(+43.99)MVDYTQLQPQR.L | 1 | Trypsin | D108 | Carboxylation |
| 98-119 | N.QAAYGGQGNPD(+15.99)MVDYTQLQPQR.L | 1 | Trypsin | D108 | Hydroxylation |
| 99-119 | Q.AAYGGQGNPD(+15.99)MVDYTQLQPQR.L | 1 | Trypsin | D108 | Hydroxylation |
| 101-114 | A.YGGQGNP(+15.99)DMVDYTQ.L | 1 | Thermolysin | P107 | Hydroxylation |
| 102-112 | Y.GGQGNPD(+15.99)MVDY.T | 2 | Chymotrypsin | D108 | Hydroxylation |
| 102-114 | Y.GGQGNPD(+15.99)MVDYTQ.L | 1 | Thermolysin | D108 | Hydroxylation |
| 102-115 | Y.GGQGNPD(+15.99)MVDYTQL.Q | 1 | Chymotrypsin | D108 | Hydroxylation |
| 102-119 | Y.GGQGNPDMVD(+15.99)YTQLQPQR.L | 1 | Trypsin | D111 | Hydroxylation |
| 102-120 | Y.GGQGNP(+15.99)DMVDYTQLQPQRL.L | 1 | Chymotrypsin | P107 | Hydroxylation |
| 105-119 | Q.GNPDMVD(+15.99)YTQLQPQR.L | 1 | Trypsin | D111 | Hydroxylation |
| 149-169 | L.AQQQHPQQQQQQQQANIS(+14.02)C(+57.02)KY.A | 1 | Chymotrypsin | S166 | Methylation |

| | | | | | |
|---|---|---|---|---|---|
| 156-169 | Q.QQQQQQQANIS(+14.02)C(+57.02)KY.A | 1 | Chymotrypsin | S166 | Methylation |
| 156-169 | Q.QQQQQQQANISC(+57.02)K(+14.02)Y.A | 1 | Chymotrypsin | K168 | Methylation |
| 158-169 | Q.QQQQQANIS(+14.02)C(+57.02)KY.A | 1 | Chymotrypsin | S166 | Methylation |
| 160-169 | Q.QQQANIS(+14.02)C(+57.02)KY.A | 1 | Chymotrypsin | S166 | Methylation |
| 162-169 | Q.QANIS(+14.02)C(+57.02)KY.A | 1 | Chymotrypsin | S166 | Methylation |
| 163-169 | Q.ANIS(+14.02)C(+57.02)KY.A | 2 | Chymotrypsin | S166 | Methylation |
| 163-169 | Q.ANISC(+57.02)K(+14.02)Y.A | 1 | Chymotrypsin | K168 | Methylation |
| 169-213 | K.YANDPVTPGGSGGGGVS(+79.97)GSNNNNNSANSNNNNSQSLASPQDLSTR.D | 1 | Trypsin | S185 | Phosphorylation |
| 169-213 | K.YANDPVTPGGSGGGGVSGSNNNNNSANSN(+.98)NNNS(+79.97)QSLASPQDLSTR.D | 1 | Trypsin | S201 | Phosphorylation |
| 214-230 | R.DISPK(+27.99)LSPSSVVESVAR.S | 4 | Trypsin | K218 | Formylation |
| 214-230 | R.DISPK(+42.01)LSPSSVVESVAR.S | 2 | Trypsin | K218 | Acetylation |
| 219-230 | K.LSPSS(+42.01)VVESVAR.S | 1 | Trypsin | S223 | Acetylation |
| 219-230 | K.LSPSSVVES(+42.01)VAR.S | 1 | Trypsin | S227 | Acetylation |
| 219-231 | K.LSPSSVVES(+42.01)VARS.L | 1 | Trypsin | S227 | Acetylation |
| 252-270 | N.NNHSGSGVSGGPGNVNVP(+15.99)M.H | 1 | Chymotrypsin | P269 | Hydroxylation |
| 255-270 | H.SGSGVSGGPGNVNVP(+15.99)M.H | 1 | Chymotrypsin | P269 | Hydroxylation |
| 259-269 | G.VSGGPGN(+.98)VNVP(+15.99).M | 1 | Thermolysin | P269 | Hydroxylation |
| 266-287 | N.VNVP(+15.99)MHSPGGGDSDSESDSGNE.A | 1 | Thermolysin | P269 | Hydroxylation |
| 266-294 | N.VNVP(+15.99)MHSPGGGDSDSESDSGNEAGSSQNS.G | 1 | Thermolysin | P269 | Hydroxylation |
| 294-307 | N.SGNGK(+43.99)KNPPQIYPW.M | 1 | Chymotrypsin | K298 | Carboxylation |
| 300-309 | K.NPPQIYP(+15.99)WMK.R | 1 | Trypsin | P306 | Hydroxylation |
| 300-309 | K.NPPQIYPW(+43.99)MK.R | 1 | Trypsin | W307 | Carboxylation |
| 300-309 | K.NPPQIYPW(+43.99)M(+15.99)K.R | 2 | Trypsin | W307 | Carboxylation |
| 300-309 | K.NPPQIYPWMK(+43.99).R | 1 | Trypsin | K309 | Carboxylation |
| 300-309 | K.NPPQIYPWM(+15.99)K(+43.99).R | 2 | Trypsin | K309 | Carboxylation |
| 300-310 | K.NPPQIYP(+15.99)WMKR.V | 1 | Trypsin | P306 | Hydroxylation |
| 300-310 | K.NPPQIYPW(+43.99)MKR.V | 2 | Trypsin | W307 | Carboxylation |

| | | | | | |
|---|---|---|---|---|---|
| 300-310 | K.NPPQIYPW(+43.99)MK(+42.01)R.V | 1 | Trypsin | W307<br>K309 | Carboxylation<br>Acetylation |
| 300-310 | K.NPPQIYPWM(+15.99)K(+27.99)R.V | 2 | Trypsin | K309 | Formylation |
| 300-310 | K.NPPQIYPWMK(+27.99)R.V | 1 | Trypsin | K309 | Formylation |
| 300-310 | K.NPPQIYPWMK(+43.99)R.V | 3 | Trypsin | K309 | Carboxylation |
| 300-310 | K.NPPQIYPWMKR(+15.99).V | 1 | Trypsin | R310 | Hydroxylation |
| 310-326 | K.RVHLGTSTVNANGETK(+43.99)R.Q | 1 | Trypsin | K325 | Carboxylation |
| 311-326 | R.VHLGT(+79.97)STVN(+.98)ANGETKR.Q | 1 | Trypsin | T315 | Phosphorylation |
| 311-326 | R.VHLGTS(+79.97)TVN(+.98)AN(+.98)GETKR.Q | 1 | Trypsin | S316 | Phosphorylation |
| 311-326 | R.VHLGTST(+79.97)VNAN(+.98)GETKR.Q | 1 | Trypsin | T317 | Phosphorylation |
| 311-326 | R.VHLGTST(+79.97)VN(+.98)AN(+.98)GETKR.Q | 1 | Trypsin | T317 | Phosphorylation |
| 311-326 | R.VHLGTSTVNAN(+15.99)GE(+57.02)TKR.Q | 1 | Trypsin | N321 | Hydroxylation |
| 311-326 | R.VHLGTSTVNANGE(+43.99)TKR.Q | 3 | Trypsin | E323 | Carboxylation |
| 311-326 | R.VHLGTSTVN(+.98)ANGE(+43.99)TKR.Q | 1 | Trypsin | E323 | Carboxylation |
| 311-326 | R.V(+57.02)HLGTSTVNANGE(+43.99)TKR.Q | 1 | Trypsin | E323 | Carboxylation |
| 311-326 | R.VHLGTSTVNANGE(+43.99)TK(+57.02)R.Q | 1 | Trypsin | E323 | Carboxylation |
| 311-326 | R.VHLGTSTVN(+.98)AN(+.98)GET(+79.97)KR.Q | 1 | Trypsin | T324 | Phosphorylation |
| 311-326 | R.VHLGTSTVN(+.98)ANGET(+79.97)KR.Q | 1 | Trypsin | T324 | Phosphorylation |
| 311-326 | R.VHLGTSTVNAN(+.98)GET(+79.97)KR.Q | 2 | Trypsin | T324 | Phosphorylation |
| 311-326 | R.VHLGTSTVNAN(+.98)GETK(+15.99)R.Q | 1 | Trypsin | K325 | Hydroxylation |
| 311-326 | R.VHLGTSTVNANGETK(+27.99)R.Q | 3 | Trypsin | K325 | Formylation |
| 311-326 | R.VHLGTSTVNAN(+.98)GETK(+27.99)R.Q | 3 | Trypsin | K325 | Formylation |
| 311-326 | R.VHLGTSTVN(+.98)ANGETK(+43.99)R.Q | 1 | Trypsin | K325 | Carboxylation |
| 311-326 | R.VHLGTSTVNANGE(+57.02)TK(+43.99)R.Q | 1 | Trypsin | K325 | Carboxylation |
| 311-326 | R.V(+57.02)HLGTSTVNANGETK(+43.99)R.Q | 1 | Trypsin | K325 | Carboxylation |
| 314-326 | L.GTSTVNANGE(+43.99)TKR.Q | 1+2^ | Trypsin & Chymotrypsin | E323 | Carboxylation |
| 314-326 | L.GTSTVNANGETK(+43.99)R.Q | 1 | Chymotrypsin | K325 | Carboxylation |

| | | | | | |
|---|---|---|---|---|---|
| 329-341 | R.TSYTRY(+15.99)QTLELEK.E | 1 | Trypsin | Y334 | Hydroxylation |
| 334-347 | R.YQTLELEK(+27.99)EFHFNR.Y | 1 | Trypsin | K341 | Formylation |
| 354-366 | R.RIEIAHALC(+57.02)LT(+14.02)ER.Q | 1 | Trypsin | T364 | Methylation |
| 354-366 | R.RIEIAHALC(+57.02)LTER(+15.99).Q | 1 | Trypsin | R366 | Hydroxylation |
| 355-366 | R.IEIAHALC(+57.02)LT(+14.02)ER.Q | 2 | Trypsin | T364 | Methylation |
| 355-366 | R.IEIAHALC(+57.02)LTER(+15.99).Q | 1 | Trypsin | R366 | Hydroxylation |
| 367-375 | R.QIK(+27.99)IWFQNR.R | 3 | Trypsin | K369 | Formylation |
| 367-375 | R.QIK(+43.99)IWFQNR.R | 1 | Trypsin | K369 | Carboxylation |
| 389-398 | M.NIVP(+15.99)YHMGPY.G | 1 | Chymotrypsin | P392 | Hydroxylation |
| 389-398 | M.NIVPYHMGPY(+15.99).G | 1 | Chymotrypsin | Y398 | Hydroxylation |
| 431-441 | D.IDYK(+42.01)DDDDKEN.L | 1 | Thermolysin | K434 | Acetylation |
| 431-441 | D.IDYK(+27.99)DDDDKEN.L | 1 | Thermolysin | K434 | Formylation |
| 431-441 | D.IDYKDDDDK(+42.01)EN.L | 1 | Thermolysin | K439 | Acetylation |
| 431-441 | D.IDYKDDDDK(+27.99)EN.L | 1 | Thermolysin | K439 | Formylation |

* - number of spectrograms in which the peptide was identified

^ - 1 spectrogram for trypsin and 2 for chymotrypsin

**Table S2. Phosphosites in bovine α-casein identified by MS/MS**

| Phosphosites mapped in bovine α-casein | Previous report |
|---|---|
| S56, S61, S63, S79, S81, S82, S83, S90, S130 | Larsen *et al.*, 2005 |
| T64, R134, Y159 | - |

Phosphosites in green – confidence shown by PEAKS database

**Table S3. Phosphopeptides of α-casein identified by MS/MS ($TiO_2$-enriched vs. non-$TiO_2$)**

| **α-casein phosphopeptides detected without $TiO_2$ (Total no. of peptides detected = 839; No. of phosphopeptides detected = 158)** | **α -casein phosphopeptides detected post-$TiO_2$ (Total no. of peptides detected = 91; No. of phosphopeptides detected = 84)** |
|---|---|
| K.DIGSES(+79.97)TEDQAMEDIK.Q | |
| K.DIGS(+79.97)ESTEDQAMEDIK.Q | |
| K.DIGS(+79.97)ES(-18.01)TEDQAMEDIK.Q | |
| K.DIGS(-18.01)ES(+79.97)TEDQAMEDIK.Q | |
| K.DIGS(+79.97)ESTEDQAM(+31.99)EDIK.Q | K.DIGS(+79.97)ESTEDQAM(+31.99)EDIK.Q |
| K.DIGS(+79.97)ESTEDQAM(+15.99)EDIK.Q | K.DIGS(+79.97)ESTEDQAM(+15.99)EDIK.Q |
| K.Y(+27.99)KVPQLEIVPNS(+79.97)AEER.L | |
| K.DIGS(+79.97)ES(+79.97)TEDQAMEDIK.Q | K.DIGS(+79.97)ES(+79.97)TEDQAMEDIK.Q |
| K.DIGS(+79.97)E(+14.02)STEDQAM(+15.99)EDIK.Q | K.DIGS(+79.97)E(+14.02)STEDQAM(+15.99)EDIK .Q |
| K.D(+57.02)IGSES(+79.97)TEDQAM(+15.99)EDIK.Q | K.D(+57.02)IGSES(+79.97)TEDQAM(+15.99)EDIK .Q |
| K.DIGSES(+79.97)TEDQAM(+31.99)EDIK.Q | K.DIGSES(+79.97)TEDQAM(+31.99)EDIK.Q |
| K.DIGSES(+79.97)TEDQAM(+15.99)EDIK.Q | K.DIGSES(+79.97)TEDQAM(+15.99)EDIK.Q |
| K.DIGS(+79.97)ESTEDQAM(+15.99)EDIK(+57.02).Q | K.DIGS(+79.97)ESTEDQAM(+15.99)EDIK(+57.02) .Q |
| K.YKVPQLEIVPNS(+79.97)AEE(+14.02)R.L | |
| K.YKVPQLEIVPN(+.98)S(+79.97)AEER.L | K.YKVPQLEIVPN(+.98)S(+79.97)AEER.L |
| K.YK(+27.99)VPQLEIVPNS(+79.97)AEER.L | |
| K.DIGSEST(+79.97)EDQAM(+15.99)EDIK.Q | K.DIGSEST(+79.97)EDQAM(+15.99)EDIK.Q |
| K.YKVPQLEIVPNS(+79.97)AE(+14.02)ER.L | |
| K.YK(+42.01)VPQLEIVPNS(+79.97)AEER.L | |
| K.YKVPQLEIVPNS(+79.97)AEER.L | K.YKVPQLEIVPNS(+79.97)AEER.L |
| K.D(+57.02)IGS(+79.97)ESTEDQAM(+15.99)EDIK.Q | K.D(+57.02)IGS(+79.97)ESTEDQAM(+15.99)EDIK .Q |

| | |
|---|---|
| K.Y(+57.02)KVPQLEIVPNS(+79.97)AEER.L | K.Y(+57.02)KVPQLEIVPNS(+79.97)AEER.L |
| K.VPQLEIVPNS(+79.97)AEER.L | K.VPQLEIVPNS(+79.97)AEER.L |
| K.VPQLEIVPN(+.98)S(+79.97)AEER.L | K.VPQLEIVPN(+.98)S(+79.97)AEER.L |
| K.VPQLE(+14.02)IVPNS(+79.97)AEER.L | K.VPQLE(+14.02)IVPNS(+79.97)AEER.L |
| K.DIGS(+79.97)ES(+79.97)TEDQAM(+15.99)EDIK.Q | K.DIGS(+79.97)ES(+79.97)TEDQAM(+15.99)EDIK .Q |
| K.DIGS(+79.97)EST(+79.97)EDQAM(+15.99)EDIK.Q | K.DIGS(+79.97)EST(+79.97)EDQAM(+15.99)EDIK .Q |
| K.VPQLEIVPNS(+79.97)AE(+14.02)ER.L | K.VPQLEIVPNS(+79.97)AE(+14.02)ER.L |
| K.VPQLEIVPNS(+79.97)AE(+57.02)ER.L | K.VPQLEIVPNS(+79.97)AE(+57.02)ER.L |
| K.DIGSES(+79.97)TEDQ(+.98)AMEDIK.Q | |
| K.DIGSEST(+79.97)EDQAMEDIK.Q | |
| K.Y(+43.01)KVPQLEIVPNS(+79.97)AEER.L | |
| K.YKVPQ(+.98)LEIVPNS(+79.97)AEER.L | |
| K.VPQLEIVPNS(+79.97)AEE(+14.02)R.L | K.VPQLEIVPNS(+79.97)AEE(+14.02)R.L |
| K.Y(+57.02)KVPQLEIVPN(+.98)S(+79.97)AEER.L | |
| K.VPQLE(+21.98)IVPNS(+79.97)AEER.L | K.VPQLE(+21.98)IVPNS(+79.97)AEER.L |
| K.D(+21.98)IGS(+79.97)ESTEDQAMEDIK.Q | |
| K.VPQLEIVP(+31.99)NS(+79.97)AEER.L | |
| K.V(+57.02)PQLEIVPN(+.98)S(+79.97)AEER.L | K.V(+57.02)PQLEIVPN(+.98)S(+79.97)AEER.L |
| K.YK(+57.02)VPQLEIVPNS(+79.97)AEER.L | K.YK(+57.02)VPQLEIVPNS(+79.97)AEER.L |
| K.DIGSES(+79.97)T(+79.97)EDQAM(+15.99)EDIK.Q | K.DIGSES(+79.97)T(+79.97)EDQAM(+15.99)EDIK .Q |
| K.YK(+43.01)VPQLEIVPNS(+79.97)AEER.L | |
| K.VNELS(+79.97)KDIGSES(+79.97)TEDQAM(+15.99) EDIK.Q | K.VNELS(+79.97)KDIGSES(+79.97)TEDQAM(+15. 99)EDIK.Q |
| K.V(+27.99)PQLEIVPNS(+79.97)AEER.L | K.V(+27.99)PQLEIVPNS(+79.97)AEER.L |
| K.V(+57.02)PQLEIVPNS(+79.97)AEER.L | K.V(+57.02)PQLEIVPNS(+79.97)AEER.L |

| | |
|---|---|
| K.VPQLEIVPNS(+79.97)AEE(+57.02)R.L | K.VPQLEIVPNS(+79.97)AEE(+57.02)R.L |
| K.DIGSES(+79.97)TEDQAM(+15.99)EDIK(+57.02).Q | K.DIGSES(+79.97)TEDQAM(+15.99)EDIK(+57.02).Q |
| K.YKVPQLE(+57.02)IVPNS(+79.97)AEER.L | K.YKVPQLE(+57.02)IVPNS(+79.97)AEER.L |
| K.DIGSES(+79.97)TEDQAMEDIK(+43.01).Q | |
| K.KYKVPQLEIVPNS(+79.97)AEER.L | |
| K.DIGS(+79.97)ES(+79.97)TEDQAME(+57.02)DIK.Q | |
| K.DIGS(+79.97)EST(+79.97)EDQAMEDIK.Q | |
| K.DIGSES(+79.97)TE(+57.02)DQAM(+15.99)EDIK.Q | |
| K.V(+57.02)PQ(+.98)LEIVPNS(+79.97)AEER.L | K.V(+57.02)PQ(+.98)LEIVPNS(+79.97)AEER.L |
| K.DIGS(+79.97)E(+57.02)STEDQAM(+15.99)EDIK.Q | |
| K.YK(+57.02)VPQLEIVPN(+.98)S(+79.97)AEER.L | |
| K.VNELSKDIGS(+79.97)ES(+79.97)TEDQAM(+15.99)EDIK.Q | K.VNELSKDIGS(+79.97)ES(+79.97)TEDQAM(+15.99)EDIK.Q |
| D.IGS(+79.97)ES(+79.97)TEDQAM(+15.99)EDIK.Q | D.IGS(+79.97)ES(+79.97)TEDQAM(+15.99)EDIK.Q |
| G.S(+79.97)ES(+79.97)TEDQAM(+15.99)EDIK.Q | G.S(+79.97)ES(+79.97)TEDQAM(+15.99)EDIK.Q |
| K.VPQLEIVPNS(+79.97)AEER(-.98).L | K.VPQLEIVPNS(+79.97)AEER(-.98).L |
| K.VPQLE(+57.02)IVPNS(+79.97)AEER.L | K.VPQLE(+57.02)IVPNS(+79.97)AEER.L |
| K.YKVPQLE(+21.98)IVPNS(+79.97)AEER.L | |
| K.Y(+57.02)K(+57.02)VPQLEIVPNS(+79.97)AEER.L | |
| K.DIGS(+79.97)ES(+79.97)TEDQAMEDIK(+57.02).Q | |
| K.DIGS(+79.97)ESTEDQAMEDIK(+43.01).Q | |
| K.VPQLEIVP(+15.99)NS(+79.97)AEER.L | |
| K.VNELS(+79.97)KDIGS(+79.97)ESTEDQAM(+15.99)EDIK.Q | K.VNELS(+79.97)KDIGS(+79.97)ESTEDQAM(+15.99)EDIK.Q |
| K.YKVPQLE(+14.02)IVPNS(+79.97)AEER.L | |
| K.VNELSKDIGS(+79.97)EST(+79.97)EDQAM(+15.99)EDIK.Q | K.VNELSKDIGS(+79.97)EST(+79.97)EDQAM(+15.99)EDIK.Q |
| K.D(+21.98)IGSES(+79.97)TEDQAMEDIK.Q | |

| | |
|---|---|
| Y.KVPQLEIVPNS(+79.97)AEER.L | |
| K.DIGSE(+21.98)S(+79.97)TEDQAMEDIK.Q | |
| K.VPQLEIVPNS(+79.97)AEER(+15.99).L | |
| K.YK(+57.02)VPQ(+.98)LEIVPNS(+79.97)AEER.L | |
| R.LK(+43.99)KYKVPQLEIVPNS(+79.97)AEER.L | |
| K.DIGS(+79.97)EST(+79.97)EDQAME(+57.02)DIK.Q | |
| K.KYK(+57.02)VPQLEIVPNS(+79.97)AEER.L | |
| K.VPQLEIVPN(+15.99)S(+79.97)AEER.L | |
| K.Y(+57.02)KVPQ(+.98)LEIVPNS(+79.97)AEER.L | |
| K.DIGS(+79.97)ES(+79.97)TE(+57.02)DQAMEDIK.Q | K.DIGS(+79.97)ES(+79.97)TE(+57.02)DQAMEDIK .Q |
| R.L(+27.99)KKYKVPQLEIVPNS(+79.97)AEER.L | |
| K.DIGSEST(+79.97)E(+21.98)DQAMEDIK.Q | |
| K.VNELSK(+57.02)DIGS(+79.97)ES(+79.97)TEDQAM EDIK.Q | |
| K.YKVPQLEIVPNS(+79.97)A.E | |
| K.VPQLEIVPNS(+79.97)AE(+57.02)E(+57.02)R.L | |
| K.YKVPQ(+.98)LE(+14.02)IVPNS(+79.97)AEER.L | |
| K.DIGSES(+79.97)T(+79.97)EDQAMEDIK.Q | |
| K.VNELS(+79.97)KDIGS(+79.97)ES(+79.97)TEDQAM (+15.99)EDIK.Q | K.VNELS(+79.97)KDIGS(+79.97)ES(+79.97)TEDQ AM(+15.99)EDIK.Q |
| K.DIGS(+79.97)ES(+79.97)TEDQAMED(+57.02)IK.Q | |
| K.YKVPQLEIVPNS(+79.97)AE(+57.02)ER.L | |
| Q.LE(+14.02)IVPNS(+79.97)AEER.L | Q.LE(+14.02)IVPNS(+79.97)AEER.L |
| P.QLEIVPNS(+79.97)AEER.L | P.QLEIVPNS(+79.97)AEER.L |
| K.VPQLEIVPNSAEER(+79.97).L | K.VPQLEIVPNSAEER(+79.97).L |
| K.Y(+15.99)KVPQLEIVPNS(+79.97)AEER.L | |
| K.Y(+42.01)KVPQLEIVPNS(+79.97)AEER.L | |

| | |
|---|---|
| K.VNELSKD(+57.02)IGSEST(+79.97)EDQAM(+15.99)EDIK.Q | |
| K.YKVP(+15.99)QLEIVPNS(+79.97)AEER.L | |
| K.DIGSES(+79.97)TEDQ(+.98)AM(+15.99)EDIK.Q | K.DIGSES(+79.97)TEDQ(+.98)AM(+15.99)EDIK.Q |
| S.ES(+79.97)TEDQAM(+15.99)EDIK.Q | S.ES(+79.97)TEDQAM(+15.99)EDIK.Q |
| K.VPQ(+.98)LE(+14.02)IVPNS(+79.97)AEER.L | |
| I.GS(+79.97)ES(+79.97)TEDQAM(+15.99)EDIK.Q | I.GS(+79.97)ES(+79.97)TEDQAM(+15.99)EDIK.Q |
| K.K(+57.02)(+57.02)YKVPQLEIVPNS(+79.97)AEER.L | |
| K.VPQLEIVPN(+.98)S(+79.97)AE(+57.02)ER.L | |
| K.Y(+57.02)KVPQLEIVPNS(+79.97)AE(+14.02)ER.L | |
| L.EIVPNS(+79.97)AEER.L | L.EIVPNS(+79.97)AEER.L |
| K.VPQLEIVPNS(+79.97)AE(+21.98)ER.L | K.VPQLEIVPNS(+79.97)AE(+21.98)ER.L |
| K.YKVPQLEIVPN(+.98)S(+79.97)AE(+14.02)ER.L | |
| K.EKVNELS(+79.97)KDIGSES(+79.97)TEDQAM(+15.99)EDIK.Q | |
| K.YKVPQLEIVP(+31.99)NS(+79.97)AEER.L | |
| L.S(+79.97)KDIGS(+79.97)ESTEDQAM(+15.99)EDIK.Q | |
| K.DIGS(+79.97)ES(+79.97)TED(+57.02)QAMEDIK.Q | |
| Q.LEIVPNS(+79.97)AEER.L | Q.LEIVPNS(+79.97)AEER.L |
| Y.K(+57.02)VPQLEIVPNS(+79.97)AEER.L | |
| R.LK(+27.99)KYKVPQLEIVPNS(+79.97)AEER.L | |
| L.SKDIGS(+79.97)ES(+79.97)TEDQAM(+15.99)EDIK.Q | |
| K.YK(+15.99)VPQLEIVPNS(+79.97)AEER.L | |
| K.YKVPQLEIVPNS(+79.97)AEER(+15.99).L | |
| K.V(+42.01)PQLEIVPNS(+79.97)AEER.L | |
| K.V(+57.02)PQLEIVPNS(+79.97)AE(+57.02)ER.L | |
| K.YKVPQLEIVPNS(+79.97)AE(+21.98)ER.L | |

| | |
|---|---|
| K.YKVPQLE(+57.02)IVPN(+.98)S(+79.97)AEER.L | |
| I.VPNS(+79.97)AEER.L | |
| L.EIVPN(+.98)S(+79.97)AEER.L | |
| K.DIGS(+79.97)ES(+79.97)TEDQN(sub A)MEDIK.Q | |
| K.YKVPQLEIVPNS(+79.97)AEER(-.98).L | |
| S.EEIVPNS(+79.97)VEQK.H | S.EEIVPNS(+79.97)VEQK.H |
| K.QM(+15.99)EAESIS(+79.97)S(+79.97)SEEIVPNSVEQK.H | |
| K.VNELS(+79.97)KDIGS(+79.97)EST(+79.97)EDQAM(+15.99)EDIK.Q | K.VNELS(+79.97)KDIGS(+79.97)EST(+79.97)EDQAM(+15.99)EDIK.Q |
| K.EKVNELS(+79.97)KDIGS(+79.97)ESTEDQAM(+15.99)EDIK.Q | |
| K.YKVPQLE(+14.02)IVPN(+.98)S(+79.97)AEER.L | |
| K.YKVPQLE(+57.02)IVPNS(+79.97)AEE(+14.02)R.L | |
| K.E(sub K)YA(sub K)VPQLEIVPNS(+79.97)AEER.L | |
| K.EKVNELS(+79.97)KDIGS(+79.97)ES(+79.97)TEDQAM(+15.99)EDIK.Q | |
| K.YKVPQLEIVPNS(+79.97)AE(+57.02)E(+57.02)R.L | |
| K.Y(+162.05)KVPQLEIVPNS(+79.97)AEER.L | |
| Q.LEIVPN(+.98)S(+79.97)AEER.L | |
| K.EKVNELS(+79.97)KDIGS(+79.97)ES(+79.97)TEDQAMEDIK.Q | |
| K.VP(+15.99)QLEIVPNS(+79.97)AEER.L | |
| K.V(+42.01)PQLE(+14.02)IVPNS(+79.97)AEER.L | |
| R.LKKYK(+27.99)VPQLEIVPNS(+79.97)AEER.L | |
| K.VNELS(+79.97)KDIG.S | |
| K.DIGSEST(+79.97)EDQAME(+57.02)DIK.Q | |
| K.QMEAESISSS(+79.97)EE(+14.02)I.V | |
| K.DIGS(+79.97)ESTE(+21.98)DQAMEDIK.Q | |
| K.DIGS(+79.97)ESTEDQAME(+57.02)DIK.Q | |

| | |
|---|---|
| Q.LE(+57.02)IVPNS(+79.97)AEER.L | |
| V.PN(+.98)S(+79.97)AEER.L | |
| K.EPQ(sub M)IGVNQELAY(+79.97)FYPELFR.Q | |
| E.EIVPNS(+79.97)VEQK.H | E.EIVPNS(+79.97)VEQK.H |
| K.DIGS(+79.97)ES(+79.97)TEDQG(sub A)P(sub M)EDIK.Q | |
| V.PNS(+79.97)AEER.L | V.PNS(+79.97)AEER.L |
| K.YKVPQLEIVPN(+.98)S(+79.97)AEER(+14.02).L | |
| K.QMEAESISSS(+79.97)EEI(+14.02).V | |
| K.VNELS(+79.97)KDIN(sub G)SES(+79.97)TEDQAM(+15.99)EDIK.Q | |
| K.VNELS(+79.97)K.D | |
| K.DIGSEST(+79.97)E(+57.02)DQAMEDIK.Q | |
| K.DIGS(+79.97)ESTEDQAMED(+57.02)IK.Q | |
| K.DIGS(+79.97)ES(+79.97)TEDQAME(+14.02)DIK.Q | |
| | K.DIGSEST(+79.97)EDQAM(+15.99)EDIK(+57.02).Q |
| | K.DIGS(+79.97)ESTED(+57.02)QAM(+15.99)EDIK.Q |
| | E.S(+79.97)TEDQAM(+15.99)EDIK.Q |
| | E.ST(+79.97)EDQAM(+15.99)EDIK.Q |
| | K.VPQLEIVPNS(+79.97)AEER(+14.02).L |
| | K.VPQ(+.98)LEIVPN(+.98)S(+79.97)AEER.L |
| | K.DIGS(+79.97)EST(+79.97)EDQAM(+15.99).E |
| | K.DIGS(+79.97)ES(+79.97)TEDQAM(+15.99).E |
| | K.DIGS(+79.97)EST(+79.97)EDQ.A |
| | K.QM(+15.99)EAESISS(+79.97)S(+79.97)EEIVPNSVEQK.H |
| | K.DIGS(+79.97)ES(+79.97)TEDQAN(sub M)EDIK.Q |

| | |
|---|---|
| | K.DIGS(+79.97)ES(+79.97)TEDQAT(sub M)EDIK.Q |
| | K.DIGS(+79.97)ES(+79.97)T(+79.97)EDQAM(+15.99)EDIK.Q |
| | S.S(+79.97)EEIVPNS(+79.97)VEQK.H |
| | E.IVPNS(+79.97)VEQK.H |
| | S.EST(+79.97)EDQAM(+15.99)EDIK.Q |
| | K.DIGS(+79.97)Q(sub E)STEDQAM(+15.99)EDIK.Q |
| | K.DIGS(+79.97)ES(+79.97)TEDQAM(+15.99)ED.I |
| | S.SEEIVPNS(+79.97)VEQK.H |
| | G.S(+79.97)EST(+79.97)EDQAM(+15.99)EDIK.Q |
| | K.DIGS(+79.97)ES(+79.97)TEDQ.A |
| | K.DIGS(+79.97)ES(+79.97)TEDQAM(+15.99)E.D |
| | K.DIGS(+79.97)ESTED(+14.02)Q(+.98)AMEDIK.Q |
| | K.QM(+15.99)EAES(+79.97)ISSS(+79.97)EEIVPNSVEQK.H |
| | K.DIGS(+79.97)ESTEDQ.A |
| | K.DIGSEST(+79.97)EDQAMED(+57.02)IK.Q |
| | S.S(+79.97)S(+79.97)EEIVPNS(+79.97)VEQK.H |
| | K.DIGS(+79.97)ES(+79.97)TEDQAL(sub M)EDIK.Q |
| | K.DIGS(+79.97)ES(+79.97)TEDQC(sub A)MEDIK.Q |
| | K.DIGSES(+79.97)TEDQ.A |

### Table S4. SLiMs in *Drosophila melanogaster* HOX proteins

| Protein | Protein (disordered) size (aa) | No. of SLiMs | SLiM size (aa) | % SLiM | No. of conserved SLiMs | Conserved SLiM size (aa) | % Conserved SLiM |
|---|---|---|---|---|---|---|---|
| LAB | 576 | 139 | 366 | 64 | 57 | 149 | 26 |
| PB | 724 | 169 | 461 | 64 | 91 | 255 | 35 |
| DFD | 533 | 84 | 287 | 54 | 55 | 188 | 35 |
| SCR | 364 | 81 | 239 | 66 | 51 | 129 | 35 |
| ANTP | 325 | 57 | 175 | 54 | 48 | 140 | 43 |
| UBX | 336 | 51 | 156 | 46 | 42 | 133 | 40 |
| ABD-A | 277 | 45 | 156 | 56 | 35 | 131 | 47 |
| ABD-B(m) | 440 | 83 | 251 | 57 | 10 | 34 | 8 |
| ABD-B(r)* | 217 | 34 | 125 | 58 | - | - | - |

* - Conserved SLiM analysis was not performed for ABD-B(r).

### Table S5. S, T and Y distribution in SLiMs vs. non-SLiMs and their significance

| Protein | Percentage of amino acid that are S, T and Y | | | | *p*-value | |
|---|---|---|---|---|---|---|
| | Total SLiMs | | Conserved SLiMs | | | |
| | SLiMs | Non-SLiMs | SLiMs | Non- SLiMs | Total SLiMs | Conserved SLiMs |
| LAB | 33 | 6 | 34 | 19 | $3.45 \times 10^{-15}$ | $4.1 \times 10^{-4}$ |
| PB | 26 | 8 | 29 | 14 | $5.73 \times 10^{-10}$ | $2.62 \times 10^{-6}$ |
| DFD | 25 | 13 | 22 | 17 | $3.9 \times 10^{-4}$ | 0.169 |
| SCR | 25 | 5 | 28 | 13 | $3.38 \times 10^{-7}$ | $5.7 \times 10^{-4}$ |
| ANTP | 23 | 5 | 25 | 8 | $3.77 \times 10^{-6}$ | $2 \times 10^{-5}$ |
| UBX | 26 | 9 | 28 | 10 | $3 \times 10^{-5}$ | $3 \times 10^{-5}$ |
| ABD-A | 21 | 4 | 22 | 6 | $3 \times 10^{-5}$ | $1.7 \times 10^{-4}$ |
| ABD-B(m) | 31 | 5 | 26 | 19 | $4.02 \times 10^{-13}$ | 0.368 |
| ABD-B(r) | 30 | 12 | - | - | $1.63 \times 10^{-3}$ | - |

### Table S6. SLiMs in *D. melanogaster* SCR

| Short Linear Motifs (SLiMs) in SCR | Amino acids | Putative Function |
|---|---|---|
| MDP | 1-3 | N-terminal motif that initiates protein degradation by binding to the UBR-box of N-recognins. This N-degron variant comprises N-terminal Asp or Glu as destabilizing residue. |
| FAMSSY | 6-11 | NEK2 phosphorylation motif with preferred Phe, Leu or Met in the -3 position to compensate for less favorable residues in the +1 and +2 position. |
| AMSSY | 7-11 | The USP7 MATH domain binding motif variant based on the MDM2 and p53 interactions. |
| SSYQF | 9-13 | Phosphopeptide motif which directly interacts with the BRCT (carboxy-terminal) domain of the Breast Cancer Gene BRCA1 with low affinity |
| SSYQFV | 9-14 | Canonical LIR motif that binds to Atg8 protein family members to mediate processes involved in autophagy. |
| SYQFV | 10-14 | Canonical LIR motif that binds to Atg8 protein family members to mediate processes involved in autophagy. |
| YQFV | 11-14 | Tyrosine-based sorting signal responsible for the interaction with mu subunit of AP (Adaptor Protein) complex |
| FVNSLA | 13-18 | NEK2 phosphorylation motif with preferred Phe, Leu or Met in the -3 position to compensate for less favorable residues in the +1 and +2 position. |
| SLASCYP | 16-22 | CK1 phosphorylation site |
| YPQQ | 21-24 | YXXQ motif found in the cytoplasmic region of cytokine receptors that bind STAT3 SH2 domain. |
| AGNSS | 34-38 | The USP7 MATH domain binding motif variant based on the MDM2 and p53 interactions. |
| GNSSAGGS | 35-42 | GSK3 phosphorylation recognition site |
| AGGSG | 39-43 | The USP7 MATH domain binding motif variant based on the MDM2 and p53 interactions. |
| AGGSG | 46-50 | The USP7 MATH domain binding motif variant based on the MDM2 and p53 interactions. |
| SGGTNGG | 55-61 | CK1 phosphorylation site |
| GAATPG | 66-71 | The Class IV WW domain interaction motif is recognised primarily by the Pin1 phosphorylation-dependent prolyl isomerase. |
| GAATPGA | 66-72 | Proline-Directed Kinase (e.g. MAPK) phosphorylation site in higher eukaryotes. |
| YFPA | 75-78 | STAT5 Src Homology 2 (SH2) domain binding motif. |
| AAYTPN | 80-85 | The Class IV WW domain interaction motif is recognised primarily by the Pin1 phosphorylation-dependent prolyl isomerase. |
| AAYTPNL | 80-86 | Proline-Directed Kinase (e.g. MAPK) phosphorylation site in higher eukaryotes. |
| AYTPNLY | 81-87 | Phosphothreonine motif binding a subset of FHA domains that show a preference for a large aliphatic amino acid at the pT+3 position. |
| YTPN | 82-85 | STAT5 Src Homology 2 (SH2) domain binding motif. |

| | | |
|---|---|---|
| PNLYP | 84-88 | PxLxP motif is recognized by a subset of MYND domain containing proteins. |
| NLYPNTP | 85-91 | This is the motif recognized by those SH3 domains with a non-canonical class I recognition specificity. |
| YPNT | 87-90 | GRB2-like Src Homology 2 (SH2) domains binding motif. |
| YPNTPQ | 87-92 | The Class IV WW domain interaction motif is recognised primarily by the Pin1 phosphorylation-dependent prolyl isomerase. |
| YPNTPQA | 87-93 | Proline-Directed Kinase (e.g. MAPK) phosphorylation site in higher eukaryotes. |
| PNTPQA | 88-93 | Phospho-dependent motif that mediates docking of CDK substrates and regulators to cyclin-CDK-bound Cks1. |
| YANQ | 95-98 | GRB2-like Src Homology 2 (SH2) domains binding motif; YXXQ motif found in the cytoplasmic region of cytokine receptors that bind STAT3 SH2 domain. |
| YGGQ | 101-104 | YXXQ motif found in the cytoplasmic region of cytokine receptors that bind STAT3 SH2 domain. |
| VDYTQLQ | 110-116 | (ST)Q motif which is phosphorylated by PIKK family members; Ser/Thr residue phosphorylated by the Plk1 kinase |
| DYTQL | 111-115 | Canonical LIR motif that binds to Atg8 protein family members to mediate processes involved in autophagy. |
| YTQL | 112-115 | STAT5 Src Homology 2 (SH2) domain binding motif; Tyrosine-based sorting signal responsible for the interaction with mu subunit of AP (Adaptor Protein) complex |
| RLLL | 119-122 | Substrate recognition site that interacts with cyclin and thereby increases phosphorylation by cyclin/cdk complexes. Predicted proteins should have a CDK phosphorylation site. Also used by cyclin/cdk inhibitors. |
| ANISCK | 163-168 | NEK2 phosphorylation motif with specific set of residues in the +1 and +2 position to compensate for less favorable residues in the -3 position. |
| YAND | 169-172 | GRB2-like Src Homology 2 (SH2) domains binding motif. |
| ANDPVTP | 170-176 | This is the motif recognized by those SH3 domains with a non-canonical class I recognition specificity. |
| DPVTPG | 172-177 | The Class IV WW domain interaction motif is recognised primarily by the Pin1 phosphorylation-dependent prolyl isomerase. |
| DPVTPGG | 172-178 | Proline-Directed Kinase (e.g. MAPK) phosphorylation site in higher eukaryotes. |
| DPVTPGGS | 172-179 | GSK3 phosphorylation recognition site |
| PVTPGG | 173-178 | Phospho-dependent motif that mediates docking of CDK substrates and regulators to cyclin-CDK-bound Cks1. |
| PVTPGGS | 173-179 | The TPxxS phospho-dependent degron binds the FBW7 F box proteins of the SCF (Skp1_Cullin-Fbox) complex. |
| VTPGGS | 174-179 | The TPxxS phospho-dependent degron binds the FBW7 F box proteins of the SCF (Skp1_Cullin-Fbox) complex. |
| PGGSG | 176-180 | The USP7 MATH domain binding motif variant based on the MDM2 and p53 interactions. |
| SANSNNN | 193-199 | CK1 phosphorylation site |

| | | |
|---|---|---|
| NNNSQSL | 198-204 | (ST)Q motif which is phosphorylated by PIKK family members. |
| SLASPQ | 203-208 | The Class IV WW domain interaction motif is recognised primarily by the Pin1 phosphorylation-dependent prolyl isomerase. |
| SLASPQD | 203-209 | CK1 phosphorylation site; Proline-Directed Kinase (e.g. MAPK) phosphorylation site in higher eukaryotes |
| DLSTRDIS | 209-216 | GSK3 phosphorylation recognition site |
| LSTRDIS | 210-216 | Phosphothreonine motif binding a subset of FHA domains that show a preference for a large aliphatic amino acid at the pT+3 position. |
| RDISPK | 213-218 | The Class IV WW domain interaction motif is recognised primarily by the Pin1 phosphorylation-dependent prolyl isomerase; Short version of the CDK phosphorylation site which shows specificity towards a lysine/arginine residue at the [ST] +2 position. |
| RDISPKL | 213-219 | Proline-Directed Kinase (e.g. MAPK) phosphorylation site in higher eukaryotes. |
| RDISPKLS | 213-220 | GSK3 phosphorylation recognition site |
| PKLSP | 217-221 | PxLxP motif is recognized by a subset of MYND domain containing proteins. |
| PKLSPS | 217-222 | The Class IV WW domain interaction motif is recognised primarily by the Pin1 phosphorylation-dependent prolyl isomerase. |
| PKLSPSS | 217-223 | Proline-Directed Kinase (e.g. MAPK) phosphorylation site in higher eukaryotes. |
| SPSSVVE | 220-226 | CK1 phosphorylation site; CK2 phosphorylation site; Ser/Thr residue phosphorylated by Plk4 |
| SPSSVVES | 220-227 | GSK3 phosphorylation recognition site |
| VVESVARS | 224-231 | GSK3 phosphorylation recognition site |
| LGGSLA | 237-242 | NEK2 phosphorylation motif with preferred Phe, Leu or Met in the -3 position to compensate for less favorable residues in the +1 and +2 position. |
| SGVSGGP | 257-263 | CK1 phosphorylation site |
| PGNVNVP | 263-269 | This is the motif recognized by those SH3 domains with a non-canonical class I recognition specificity. |
| PMHSPG | 269-274 | The Class IV WW domain interaction motif is recognised primarily by the Pin1 phosphorylation-dependent prolyl isomerase. |
| PMHSPGG | 269-275 | Proline-Directed Kinase (e.g. MAPK) phosphorylation site in higher eukaryotes. |
| GGDSDSE | 275-281 | CK2 phosphorylation site |
| GGDSDSES | 275-282 | GSK3 phosphorylation recognition site |
| DSDSESDS | 277-284 | GSK3 phosphorylation recognition site |
| SESDS | 280-284 | Caspase-3 and Caspase-7 cleavage site. |
| ESDSGNE | 281-287 | CK2 phosphorylation site |
| EAGSSQNS | 287-294 | GSK3 phosphorylation recognition site |
| AGSSQ | 288-292 | The USP7 MATH domain binding motif variant based on the MDM2 and p53 interactions. |
| AGSSQNS | 288-294 | (ST)Q motif which is phosphorylated by PIKK family members. |
| SQNSGNG | 291-297 | CK1 phosphorylation site |

| | | |
|---|---|---|
| YPWM | 305-308 | The YPWM motif confers binding to the PBX homeobox domain; Tyrosine-based sorting signal responsible for the interaction with mu subunit of AP (Adaptor Protein) complex |
| KRVHLGTSTVNANGETKRQRT | 309-329 | Bipartite variant of the classical basically charged NLS. |
| LGTSTV | 313-318 | NEK2 phosphorylation motif with preferred Phe, Leu or Met in the -3 position to compensate for less favorable residues in the +1 and +2 position. |
| LGTSTVN | 313-319 | Phosphothreonine motif binding a subset of FHA domains that show a preference for a large aliphatic amino acid at the pT+3 position. |
| YHQF | 402-405 | Tyrosine-based sorting signal responsible for the interaction with mu subunit of AP (Adaptor Protein) complex |
| IHPSQFA | 407-413 | (ST)Q motif which is phosphorylated by PIKK family members. |
| SQFAHL | 410-415 | Canonical LIR motif that binds to Atg8 protein family members to mediate processes involved in autophagy. |
| FAHLSA | 412-417 | The C-terminal class 2 PDZ-binding motif is classically represented by a pattern such as (VYF)X(VIL)* |

Red - Amino acid residues that are phosphorylated

Blue - Amino acid residues that are acetylated, formylated, methylated, carboxylated or hydroxylated

**Table S7. Conserved SLiMs in *D. melanogaster* SCR**

| Short Linear Motifs (SLiMs) in SCR | Amino acids | Taxonomic level of conservation | Putative Function |
|---|---|---|---|
| FAMSSY | 6-11 | *Drosophila* | NEK2 phosphorylation motif with preferred Phe, Leu or Met in the -3 position to compensate for less favorable residues in the +1 and +2 position. |
| AMSSY | 7-11 | *Drosophila* | The USP7 MATH domain binding motif variant based on the MDM2 and p53 interactions. |
| SSYQF | 9-13 | Arthropoda | Phosphopeptide motif which directly interacts with the BRCT (carboxy-terminal) domain of the Breast Cancer Gene BRCA1 with low affinity |
| SSYQFV | 9-14 | Arthropoda | Canonical LIR motif that binds to Atg8 protein family members to mediate processes involved in autophagy. |
| SYQFV | 10-14 | Arthropoda | Canonical LIR motif that binds to Atg8 protein family members to mediate processes involved in autophagy. |
| YQFV | 11-14 | Arthropoda | Tyrosine-based sorting signal responsible for the interaction with mu subunit of AP (Adaptor Protein) complex |
| FVNSLA | 13-18 | Arthropoda | NEK2 phosphorylation motif with preferred Phe, Leu or Met in the -3 position to compensate for less favorable residues in the +1 and +2 position. |
| SLASCYP | 16-22 | *Drosophila* | CK1 phosphorylation site |
| GAATPG | 66-71 | *Drosophila* | The Class IV WW domain interaction motif is recognised primarily by the Pin1 phosphorylation-dependent prolyl isomerase. |
| GAATPGA | 66-72 | *Drosophila* | Proline-Directed Kinase (e.g. MAPK) phosphorylation site in higher eukaryotes. |
| AAYTPN | 80-85 | *Drosophila* | The Class IV WW domain interaction motif is recognised primarily by the Pin1 phosphorylation-dependent prolyl isomerase. |
| AAYTPNL | 80-86 | *Drosophila* | Proline-Directed Kinase (e.g. MAPK) phosphorylation site in higher eukaryotes. |
| AYTPNLY | 81-87 | *Drosophila* | Phosphothreonine motif binding a subset of FHA domains that show a preference for a large aliphatic amino acid at the pT+3 position. |
| PNLYP | 84-88 | *Drosophila* | PxLxP motif is recognized by a subset of MYND domain containing proteins. |
| NLYPNTP | 85-91 | *Drosophila* | This is the motif recognized by those SH3 domains with a non-canonical class I recognition specificity. |
| YPNTPQ | 87-92 | *Drosophila* | The Class IV WW domain interaction motif is recognised primarily by the Pin1 phosphorylation-dependent prolyl isomerase. |
| YPNTPQA | 87-93 | *Drosophila* | Proline-Directed Kinase (e.g. MAPK) phosphorylation site in higher eukaryotes. |

| | | | |
|---|---|---|---|
| PNTPQA | 88-93 | *Drosophila* | Phospho-dependent motif that mediates docking of CDK substrates and regulators to cyclin-CDK-bound Cks1. |
| VDYTQLQ | 110-116 | Diptera | (ST)Q motif which is phosphorylated by PIKK family members. |
| DYTQL | 111-115 | Insecta | Canonical LIR motif that binds to Atg8 protein family members to mediate processes involved in autophagy. |
| YTQL | 112-115 | Insecta | STAT5 Src Homology 2 (SH2) domain binding motif; Tyrosine-based sorting signal responsible for the interaction with mu subunit of AP (Adaptor Protein) complex |
| PGGSG | 176-180 | *Drosophila* | The USP7 MATH domain binding motif variant based on the MDM2 and p53 interactions. |
| SLASPQ | 203-208 | *Drosophila* | The Class IV WW domain interaction motif is recognised primarily by the Pin1 phosphorylation-dependent prolyl isomerase. |
| SLASPQD | 203-209 | *Drosophila* | CK1 phosphorylation site; Proline-Directed Kinase (e.g. MAPK) phosphorylation site in higher eukaryotes |
| DLSTRDIS | 209-216 | *Drosophila* | GSK3 phosphorylation recognition site |
| LSTRDIS | 210-216 | *Drosophila* | Phosphothreonine motif binding a subset of FHA domains that show a preference for a large aliphatic amino acid at the pT+3 position. |
| RDISPK | 213-218 | *Drosophila* | The Class IV WW domain interaction motif is recognised primarily by the Pin1 phosphorylation-dependent prolyl isomerase; Short version of the CDK phosphorylation site which shows specificity towards a lysine/arginine residue at the [ST] +2 position. |
| RDISPKL | 213-219 | *Drosophila* | Proline-Directed Kinase (e.g. MAPK) phosphorylation site in higher eukaryotes. |
| RDISPKLS | 213-220 | *Drosophila* | GSK3 phosphorylation recognition site |
| PKLSP | 217-221 | *Drosophila* | PxLxP motif is recognized by a subset of MYND domain containing proteins. |
| PKLSPS | 217-222 | *Drosophila* | The Class IV WW domain interaction motif is recognised primarily by the Pin1 phosphorylation-dependent prolyl isomerase. |
| PKLSPSS | 217-223 | *Drosophila* | Proline-Directed Kinase (e.g. MAPK) phosphorylation site in higher eukaryotes. |
| SPSSVVE | 220-226 | *Drosophila* | CK1 phosphorylation site; CK2 phosphorylation site |
| SPSSVVES | 220-227 | *Drosophila* | GSK3 phosphorylation recognition site |
| VVESVARS | 224-231 | *Drosophila* | GSK3 phosphorylation recognition site |
| LGGSLA | 237-242 | *Drosophila* | NEK2 phosphorylation motif with preferred Phe, Leu or Met in the -3 position to compensate for less favorable residues in the +1 and +2 position. |
| PMHSPG | 269-274 | *Drosophila* | The Class IV WW domain interaction motif is recognised primarily by the Pin1 phosphorylation-dependent prolyl isomerase. |

| | | | |
|---|---|---|---|
| PMHSPGG | 269-275 | *Drosophila* | Proline-Directed Kinase (e.g. MAPK) phosphorylation site in higher eukaryotes. |
| GGDSDSE | 275-281 | *Drosophila* | CK2 phosphorylation site |
| GGDSDSES | 275-282 | *Drosophila* | GSK3 phosphorylation recognition site |
| DSDSESDS | 277-284 | *Drosophila* | GSK3 phosphorylation recognition site |
| SESDS | 280-284 | *Drosophila* | Caspase-3 and Caspase-7 cleavage site. |
| ESDSGNE | 281-287 | *Drosophila* | CK2 phosphorylation site |
| EAGSSQNS | 287-294 | *Drosophila* | GSK3 phosphorylation recognition site |
| AGSSQ | 288-292 | *Drosophila* | The USP7 MATH domain binding motif variant based on the MDM2 and p53 interactions. |
| AGSSQNS | 288-294 | *Drosophila* | (ST)Q motif which is phosphorylated by PIKK family members. |
| SQNSGNG | 291-297 | *Drosophila* | CK1 phosphorylation site |
| YPWM | 305-308 | Bilateria | The YPWM motif confers binding to the PBX homeobox domain; Tyrosine-based sorting signal responsible for the interaction with mu subunit of AP (Adaptor Protein) complex |
| IHPSQFA | 407-413 | Diptera | (ST)Q motif which is phosphorylated by PIKK family members. |
| SQFAHL | 410-415 | Insecta | Canonical LIR motif that binds to Atg8 protein family members to mediate processes involved in autophagy. |
| FAHLSA | 412-417 | Insecta | The C-terminal class 2 PDZ-binding motif is classically represented by a pattern such as (VYF)X(VIL)* |

Red - Amino acid residues that are phosphorylated

Blue - Amino acid residues that are acetylated, formylated, methylated, carboxylated or hydroxylated

**Figure S9. Alignment of SLiMs of animal SCR homologs.** SLiMs of *D. melanogaster* SCR which are conserved across various taxonomic groups were aligned using multiple sequence alignment tools, MAFFT v. 7 and Clustal Omega. The phylogenetic tree on the left is not drawn to scale and it merely depicts the relationship among the organisms and not evolutionary time of divergence. The block diagram below the aligned sequences shows the HD and four SLiMs conserved beyond Diptera.

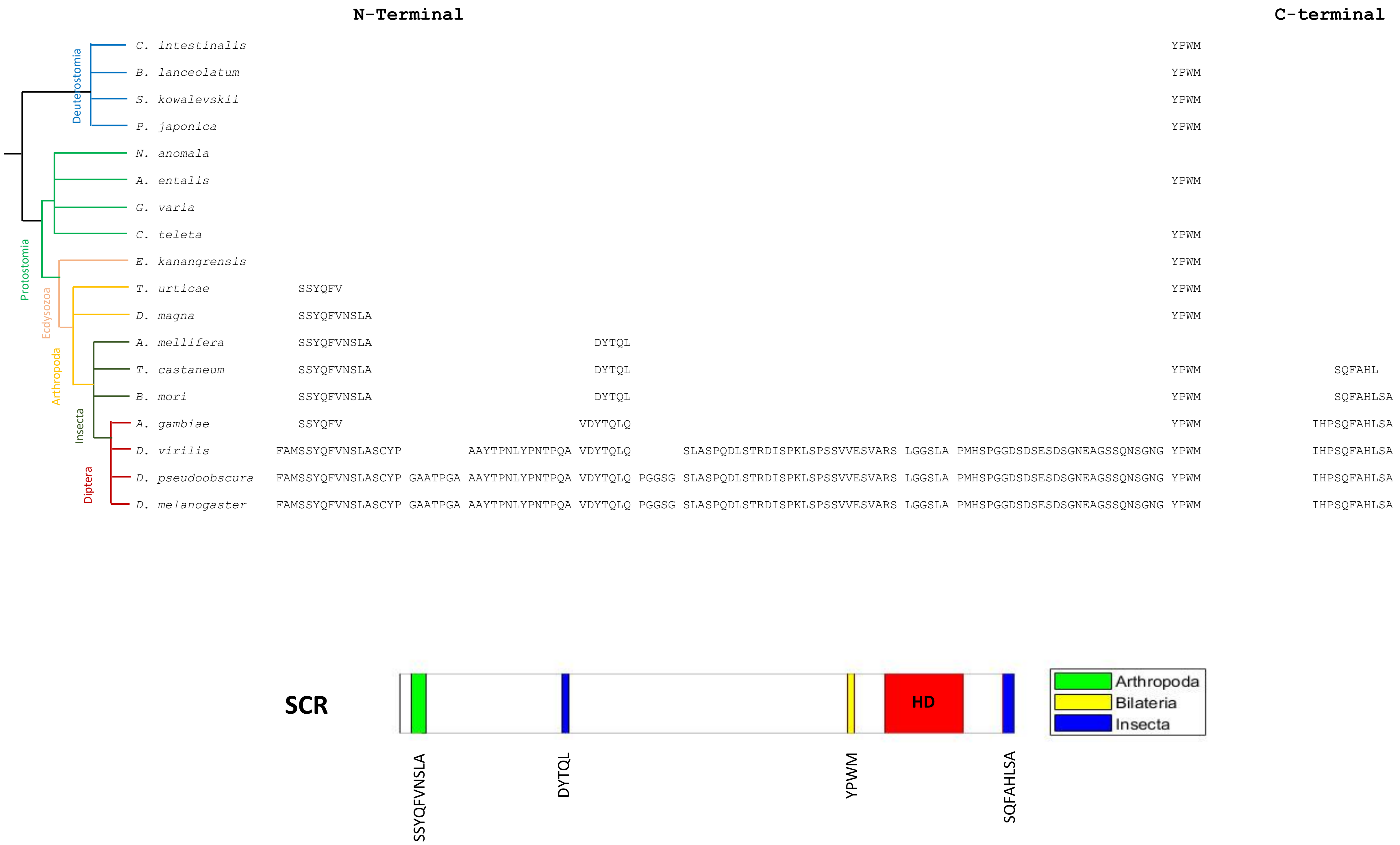
N-Terminal
C-terminal
Deuterostomia
Protostomia
Ecdysozoa
Arthropoda
Insecta
Diptera
C. intestinalis YPWM
B. lanceolatum YPWM
S. kowalevskii YPWM
P. japonica YPWM
N. anomala
A. entalis YPWM
G. varia
C. teleta YPWM
E. kanangrensis YPWM
T. urticae SSYQFV YPWM
D. magna SSYQFVNSLA YPWM
A. mellifera SSYQFVNSLA DYTQL
T. castaneum SSYQFVNSLA DYTQL YPWM SQFAHL
B. mori SSYQFVNSLA DYTQL YPWM SQFAHLSA
A. gambiae SSYQFV VDYTQLQ YPWM IHPSQFAHLSA
D. virilis FAMSSYQFVNSLASCYP AAYTPNLYPNTPQA VDYTQLQ SLASPQDLSTRDISPKLSPSSVVESVARS LGGSLA PMHSPGGDSDSESDSGNEAGSSQNSGNG YPWM IHPSQFAHLSA
D. pseudoobscura FAMSSYQFVNSLASCYP GAATPGA AAYTPNLYPNTPQA VDYTQLQ PGGSG SLASPQDLSTRDISPKLSPSSVVESVARS LGGSLA PMHSPGGDSDSESDSGNEAGSSQNSGNG YPWM IHPSQFAHLSA
D. melanogaster FAMSSYQFVNSLASCYP GAATPGA AAYTPNLYPNTPQA VDYTQLQ PGGSG SLASPQDLSTRDISPKLSPSSVVESVARS LGGSLA PMHSPGGDSDSESDSGNEAGSSQNSGNG YPWM IHPSQFAHLSA
SCR
HD
Arthropoda
Bilateria
Insecta
SSYQFVNSLA
DYTQL
YPWM
SQFAHLSA

**Table S8. Accession numbers of animal HOX protein homolog sequences (retrieved from NCBI)**

| **Organism** | **Taxonomic group** | **LAB** | **PB** | **DFD** | **SCR** | **ANTP** | **UBX** | **ABD-A** | **ABD-B** |
|---|---|---|---|---|---|---|---|---|---|
| *Drosophila melanogaster* | Diptera | AAF54098.1 | NP_996163.1 | AAF54083.2 | NP_524248.2 | NP_996175.1 | NP_536752.1 | NP_476693.1 | NP_524896.2 (isoform m) NP_650577.1 (isoform r) |
| **Organism** | **Taxonomic group** | ***Drosophila melanogaster* HOX homologs** | | | | | | | |
| *Drosophila pseudoobscura* | Diptera | EDY68244.1 | EIM52773.2 | EAL28360.2 | EAL28358.3 | EIM52770.2 | EAL28209.2 | EDY68153.2 | XP_003736655.2 |
| *Drosophila virilis* | Diptera | EDW67051.1 | XP_002053521.2 | XP_002053514.2 | AAO01077.1 (partial) +XP_015027472.1 | KRF83066.1 | EDW67025.1 | EDW58935.2 | KRF78610.1 |
| *Anopheles gambiae* | Diptera | XP_001688960.2 | XP_001688961.2 | XP_001688962.1 | AAC31944.1 | AAC31945.1 | AAC31942.1 | AAC31946.1 | ABD38144.1 |
| *Bombyx mori* | Lepidoptera | - | XP_021206544.1 | NP_001037341.1 | BAA76868.1 | NP_001037319.1 | NP_001107632.1 | ACD10794.1 | BAH30152.1 |
| *Tribolium castaneum* | Coleoptera | EEZ99257.1 | NP_001107807.1 (Maxillopedia) | AAK16423.1 | AAK16422.1 | EEZ99250.1 | EEZ99249.1 | EEZ99248.1 | EEZ99247.1 |
| *Apis mellifera* | Hymenoptera | XP_006559411.1 | XP_006559426.1 | XP_001120045.2 | XP_623903.2 | CAC06383.1 | NP_001162171.1 | XP_394120.4 | XP_006559430.1 |
| *Schistocerca americana* | Orthoptera | - | - | - | - | AAB03236.1 | - | - | - |
| *Daphnia pulex/magna* | Crustacea | EFX86813.1 | EFX86812.1 | EFX86808.1 | BAJ05331.1 | BAE96993.1 | BAE96992.1 | EFX86800.1 | EFX86798.1 |
| *Tetranychus urticae** | Chelicarata | tetur11g05940 | tetur11g05920 | tetur20g02580 | tetur20g02540 | tetur20g02430; | tetur20g02400 | - | tetur20g02350 |

| | | | | | | | | | |
|---|---|---|---|---|---|---|---|---|---|
| | | | | | | tetur20g02440 | | | |
| *Strigamia maritima* | Myriapoda | - | - | - | - | - | - | ABD16213.1 | - |
| *Euperipatoides kanangrensis* | Onychophora | CCK73369.1 | CCK73370.1 | CCK73372.1 | CCK73373.1 | CCK73375.2 | CCK73376.1 | CCK73377.1 | CCK73379.1 |
| *Chaetopterus variopedatus* | Annelida | AAD55932.1 (Hox1) | AAD55933.1 (Hox2) | AAD55935.1 (partial) (Hox4) | - | - | - | - | - |
| *Capitella teleta* | Annelida | - | - | ABY67955.1 | ABY67956.1 | ABY67962.1 | ABY67958.1 (Lox4); ABY67959.1 (Lox2) | ABY67958.1 (Lox4); ABY67959.1 (Lox2) | ABY67961.1 (Post1); ABY67960.1 (Post2) |
| *Helobdella triserialis/ robusta* | Annelida | CAA71835.1 (Lox7) | - | - | - | AAB61442.1 (partial) (Lox5) | - | - | - |
| *Gibbula varia* | Mollusca | ACX84671.1 (Hox1) | ADJ18233.1 (Hox2) | ACX84672.1 (partial) (Hox4) | ADJ18234.1 (Hox5) | ADJ18236.1 (Hox7) | ADJ18237.1 (Lox4); ADJ18238.1 (Lox2) | ADJ18237.1 (Lox4); ADJ18238.1 (Lox2) | - |
| *Antalis entalis* | Mollusca | APD15651.1 (Hox1) | APD15652.1 (partial) (Hox2) | APD15654.1 (Hox4) | APD15655.1 (Hox5) | - | APD15658.1 (Lox4); APD15657.1 (Lox2) | APD15658.1 (Lox4); APD15657.1 (Lox2) | APD15659.1 (Post1); APD15660.1 (Post2) |
| *Gymnomenia pellucida* | Mollusca | - | - | - | - | APD15669.1 (Hox7) | - | - | APD15672.1 (Post1); APD15673.1 (Post2) |
| *Novocrania anomala* | Brachiopoda | ANO46564.1 | ANO46565.1 | ANO46577.1 (partial) | ANO46567.1 | - | - | - | - |
| *Peronella japonica* | Echinodermata | BAO57695.1 (Hox1) | BAO57696.1 (Hox2) | - | BAO57698.1 (Hox5) | BAO57700.1 (Hox7) | BAO57701.1 (Hox8) | BAO57701.1 (Hox8) | BAO57703.1 (Hox11/13a); BAO57704.1 (Hox11/13b) BAO57705.1 (Hox11/13c) |

| | | | | | | | | | |
|---|---|---|---|---|---|---|---|---|---|
| *Saccogloccus kowalevskii* | Hemichordata | AAP79296.1 (Hox1) | ABK00018.1 (Hox2) | AAP79297.1 (Hox4) | ABK00019.1 (Hox5) | NP_0011583 80.1 (Hox7) | - | - | ABK00021.1 (Hox9/10); ABK00022.1 (Hox11/13a); ABK00023.1 (Hox11/13b); AAP79288.1 (Hox11/13c) |
| *Balanoglossus misakiensis* | Hemichordata | - | - | - | - | - | BAH23877.1 (Hox8) | BAH23877.1 (Hox8) | - |
| *Branchiostoma floridae/ lanceolatum* | Cephalochordata /Amphioxus | BAA78620.2 (Hox1) | BAA78621.1 (Hox2) | BAA78622.1 (Hox4) | ACJ74385.1 (Hox5) | ACJ74388.1 (Hox7) | ACJ74387.1 (Hox8) | ACJ74387.1 (Hox8) | - |
| *Ciona intestinalis* | Urochordata/ Tunicata | NP_0011223 33.1 (Hox1) | CAD59668.1 (partial) (Hox2) | NP_001027 781.2 (Hox4) | NP_0010276 65.1 (Hox5) | - | - | - | - |

***** *Tetranychus urticae* HOX protein sequences were retrieved from the *Tetranychus urticae* database in Online Resource for Community Annotation of Eukaryotes (ORCAE) - http://bioinformatics.psb.ugent.be/orcae/overview/Tetur
